# A K27-linked Ubiquitin Checkpoint Controls NOTCH Homeostasis

**DOI:** 10.64898/2025.12.18.695023

**Authors:** Behzad Mansoori, Ying Song, Tian Zhang, Zihan Zheng, Qing Zhu, Shun Li, Mckenna Reale, Christian Pangilinan, Janvhi Suresh Machhar, Jinghui Liang, Jason Chang, Young-Kwon Hong, Gerald B. Wertheim, Maureen E. Murphy, Yong-Mi Kim, Markus Muschen, Michaela U Gack, Andrew V. Kossenkov, Qin Liu, Noam Auslander, Iannis Aifantis, Chintan Parekh, Chengyu Liang

## Abstract

Cell surface receptors such as NOTCH1 must be tightly regulated to ensure developmental fidelity and prevent pathological activation. Although the proteolytic steps culminating in nuclear NOTCH1 signaling are established, how cells prevent excessive or uncontrolled activation has remained unresolved. Here we identify the autophagy-related protein UVRAG as a negative regulator of NOTCH1. Upon receptor activation, UVRAG, acting independently of autophagy, recruits and activates the E3-ligase ITCH to catalyze K27-linked ubiquitination of membrane-tethered NOTCH1, thereby licensing ESCRT-dependent lysosomal degradation. Disruption of the UVRAG-ITCH-ESCRT axis stabilizes activated NOTCH1 intermediates and amplifies oncogenic signaling. In T-cell leukemia models driven by constitutive NOTCH1 activity, restoring UVRAG expression reinstates receptor turnover, suppresses disease progression, and improves therapeutic response. These findings define a ubiquitin-directed safeguard circuit that enforces NOTCH1 signaling homeostasis and reveals a tunable axis for intervention in NOTCH1-driven cancers.

## INTRODUCTION

The NOTCH signaling pathway is conserved across metazoans and plays a fundamental role in cell fate specification, stem cell maintenance, and tissue homeostasis (Kopan and Ilagan, 2009). In mammals, activation of the NOTCH1 receptor is triggered by binding of Delta-like or Jagged ligands presented on adjacent cells, which enables ADAM-family metalloprotease cleavage (S2) of the extracellular domain. This event generates a membrane-tethered intermediate, NOTCH1ΔE, which undergoes subsequent intramembrane proteolysis by the γ-secretase complex (S3), releasing the intracellular domain (NICD). NICD translocates to the nucleus and drives transcriptional programs that govern developmental and physiological processes (Ferrando, 2009; Kopan and Ilagan, 2009). Long-standing evidence indicates that NOTCH1 signaling is exquisitely dosage-sensitive; small changes in signal strength can drastically alter lineage trajectories during development or promote malignant transformation in disease (Chiang et al., 2008; Izon et al., 2001; Schweisguth, 1995). This is particularly evident in T-cell acute lymphoblastic leukemia (T-ALL), where activating mutations in NOTCH1 promote ligand-independent activation or impaired NICD turnover, leading to sustained transcriptional activation (Aifantis et al., 2008; Ferrando, 2009).

To maintain signaling fidelity, several negative regulatory mechanisms constrain NOTCH1 pathway output. Nuclear NICD is targeted for proteasomal degradation by the E3 ligase FBXW7, serving as a feedback mechanism to restrict transcriptional duration (O’Neil et al., 2007; Thompson et al., 2007). Separately, endocytic regulators such as Rab5 and ubiquitin ligases including Nedd4 and Deltex have been shown to modulate receptor internalization and sorting decisions in endosomes, buffering surface receptor levels (Fortini and Bilder, 2009; Fuwa et al., 2006; Wilkin et al., 2004). While these findings highlight the importance of endosomal compartments in the regulation of NOTCH homeostasis, they primarily affect unliganded, full-length receptors and do not account for how signaling is modulated after activation has been initiated. Once S2 cleavage occurs, the receptor is generally thought to passively undergo rapid γ-secretase processing to release NICD. However, whether and how endosomal compartments regulate the flux or fate of this activated intermediate remains unknown. This gap is particularly salient given the central positioning of NOTCH1ΔE as a bottleneck for signal propagation. Genetic studies in Drosophila have implicated the UV radiation resistance-associated gene (UVRAG), a regulator of autophagy and endosome function (Itakura et al., 2008; Liang et al., 2006; Liang et al., 2008), in modulating Notch activity (Lee et al., 2011), but the underlying mechanism and its relevance to mammalian NOTCH1 signaling remain undefined. We investigated whether UVRAG regulates NOTCH1 signaling by controlling the fate of the activated, S2-cleaved receptor intermediate prior to NICD release.

## RESULTS

### UVRAG Negatively Regulates NOTCH1 Activation

To investigate whether UVRAG regulates NOTCH1 activity in human cells, we used HeLa cells expressing wild-type NOTCH1 and stimulated them with the canonical ligand DLL4. NOTCH1 activity was measured using TP1- and CBF1-luciferase reporters, as well as endogenous target genes HES1 and HEY1. UVRAG depletion using two independent shRNAs significantly enhanced ligand-induced reporter activity and target gene expression, with no effect on baseline levels (Figures 1A, 1B, and S1A). Conversely, overexpression of UVRAG reduced ligand-induced NOTCH1 activity (Figures S1B and S1C), suggesting that UVRAG acts as a negative regulator of ligand-triggered NOTCH1 signaling.

**Figure 1.**
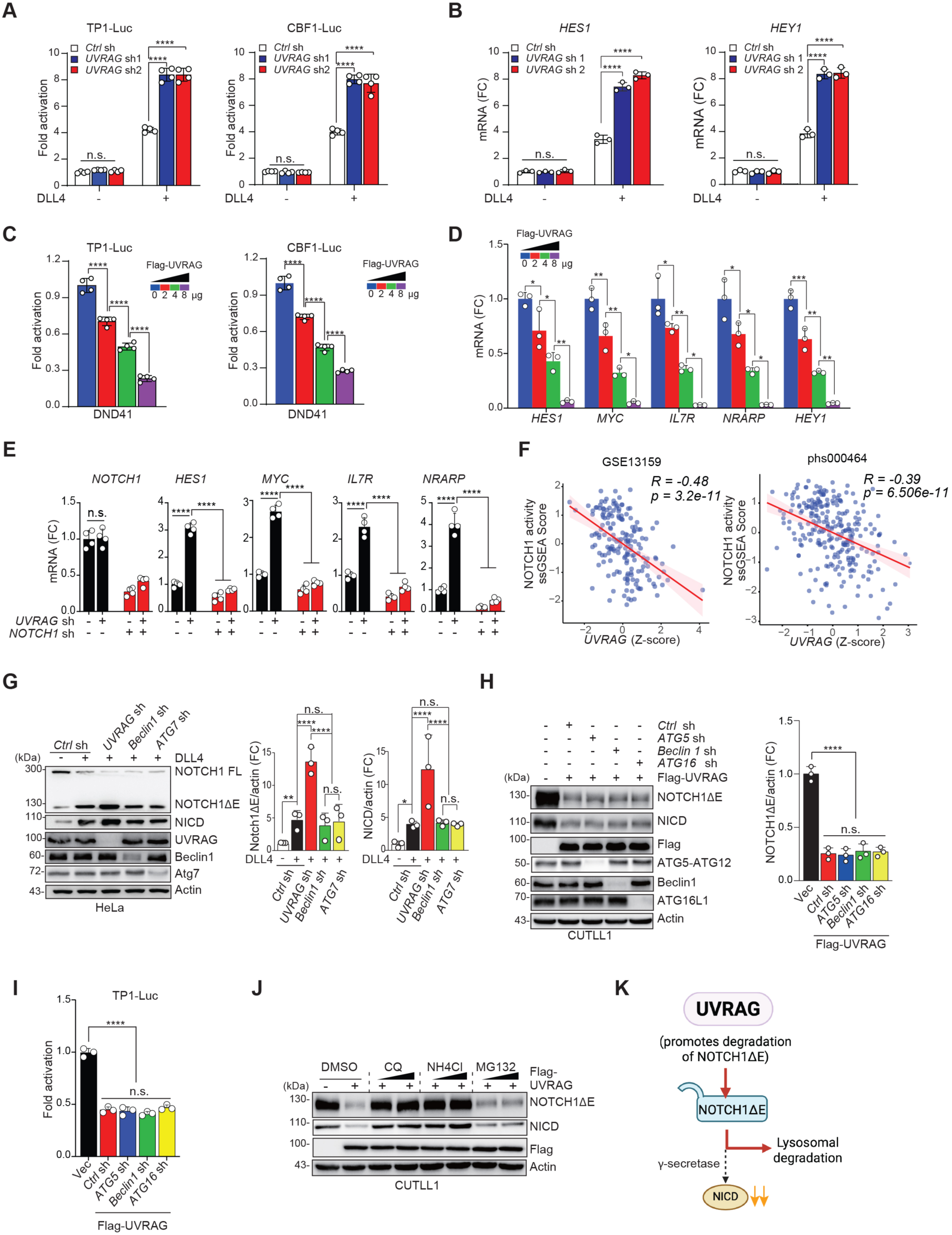
UVRAG Suppresses NOTCH1 Activation by Promoting Lysosomal Degradation of NOTCH1ΔE. (A) TP1- (left) and CBF1- (right) luciferase activity in HeLa cells expressing *Control* (*Ctrl*) or *UVRAG-*specific shRNAs (sh1 and sh2) with or without DLL4 stimulation (n = 4). See also Figure S1A for UVRAG expression in cells in (A). (B) RT-qPCR quantification of *HES1* and *HEY1* transcripts in cells from (A). n = 3. (C) Dose-dependent inhibition of NOTCH1 signaling measured by TP1- (left) and CBF1- (right) luciferase activity in DND41 T-ALL cells transfected with increasing amounts of Flag-UVRAG. n = 4. (D) RT-qPCR quantification of indicated NOTCH1 target genes in cells from (C), normalized to control (Flag-UVRAG 0 μg). n = 3. (E) RT-qPCR quantification of indicated gene transcripts in *Ctrl* or *UVRAG*-knockdown DND41 cells expressing *NOTCH1-*specific shRNA. n = 4. (F) Scatter plot correlating UVRAG expression (Z-score normalized) with the ssGSEA scores of NOTCH1 signature (Palomero et al., 2006b) in T-ALL patient samples from two datasets (GSE13159, n = 174; dbGaP phs000464 T-ALL, n = 264). (G) Immunoblotting (IB, left) and densitometric analysis (right) of NOTCH1ΔE/actin and NICD/actin in HeLa cells transduced with indicated shRNAs after DLL4 stimulation. Actin serves as loading control. See also Figure S2A for NOTCH1 mRNA expression. (H) IB analysis (left) and densitometric quantification (right) of NOTCH1ΔE/actin ratios in CUTLL1 cells stably expressing Flag-UVRAG and/or transduced with indicated shRNAs. (I) TP1-luciferase activity in cells from (H). n = 3. (J) IB analysis of NOTCH1ΔE and NICD in CUTLL1 cells stably expressing Vec (-) or Flag-UVRAG (+) treated with DMSO, lysosomal inhibitors (chloroquine, CQ, 20 mM; NH4Cl, 10 mM) or proteasomal inhibitor (MG132, 10 mM). (K) Working model depicting UVRAG-directed lysosomal degradation of membrane-bound NOTCH1ΔE upstream of γ-secretase cleavage. Data in (G), (H), and (J) are representative of three independent experiments. Data in (A-E) and (G-I) represent mean ± s.d. from biologically independent samples (n ≥ 3), analyzed by one-way ANOVA with Tukey’s *post hoc* test. *, *p* < 0.05; **, *p* < 0.01; ***, *p* < 0.001; ****, *p* < 0.0001; ns, not significant. See also Figure S1 and Figure S2.

To determine whether UVRAG also restrains ligand-independent NOTCH1 activation driven by oncogenic mutations, we analyzed human T-ALL cell lines harboring activating mutations in *NOTCH1* commonly found in T-ALL patients (Sanchez-Irizarry et al., 2004). In DND41 cells, which carry the L1594P mutation that permits S2 cleavage in the absence of ligand (Sanchez-Irizarry et al., 2004), UVRAG overexpression suppressed reporter activity and expression of NOTCH1 target genes in a dose-dependent manner (Figures 1C and 1D). Knockdown of *UVRAG* increased target gene expression, and this effect was reversed by silencing *NOTCH1* itself (Figure 1E). Similar results were observed in CUTLL1 cells, which express a membrane-tethered, constitutively active NOTCH1 mutant (Palomero et al., 2006a) (Figures S1D and S1E). Reintroduction of wild-type (WT) UVRAG into *UVRAG*-deficient CUTLL1 cells restored repression of NOTCH1 target genes (Figure S1E), directly linking UVRAG loss to enhanced signaling. Consistent with these findings, single-sample gene set enrichment analysis (ssGSEA) of human T-ALL patient datasets revealed an inverse correlation between UVRAG expression and NOTCH1 target gene signatures (Palomero et al., 2006b) (Figure 1F). These findings collectively underscore the general role of UVRAG in restraining both ligand-induced and mutation-driven NOTCH1 signaling activation.

### UVRAG Targets Membrane-bound NOTCH1ΔE for Lysosomal Degradation Independent of Autophagy

*NOTCH1* transcript levels remained unchanged following *UVRAG* depletion or overexpression in both HeLa and T-ALL cells (Figures S2A and S2B), suggesting post-transcriptional regulation. To determine whether this regulation occurs at the protein level, we examined the effect of UVRAG on ligand-induced NOTCH1 processing. In DLL4-stimulated HeLa cells, UVRAG overexpression reduced levels of the membrane-tethered NOTCH1ΔE and cleaved NICD in a dose-dependent manner, without accumulation of full-length receptor (Figure S2C). NICD was detected using a V1744-specific antibody, which marks γ-secretase activity (Schroeter et al., 1998). Conversely, *UVRAG* knockdown increased levels of both NOTCH1ΔE and NICD upon ligand stimulation (Figure 1G), consistent with enhanced signaling (Figures 1A and 1B). Similar effects were observed in T-ALL cells harboring active *NOTCH1* mutants, where UVRAG overexpression suppressed NOTCH1ΔE and NICD abundance (Figure S2D). Across a panel of T-ALL cell lines and primary samples, UVRAG expression inversely correlated with NICD levels (Figures S2E and S2F), supporting a broader regulatory relationship. Cycloheximide (CHX) chase experiments revealed that UVRAG selectively accelerated degradation of NOTCH1ΔE but had no effect on NICD stability (Figures S2G and S2H), suggesting that reduced NICD levels result from diminished precursor availability. Consistently, UVRAG reduced reporter activity driven by NOTCH1ΔE, but not when NICD was directly expressed (Figure S2I). Furthermore, UVRAG depleted NOTCH1ΔE even in the presence of γ-secretase inhibition by DAPT (Dovey et al., 2001) or when using a cleavage-resistant V1744L mutant (Schroeter et al., 1998) (Figure S2J), indicating that UVRAG acts upstream of S3 cleavage.

Although UVRAG is classically associated with autophagy through its interaction with the Beclin1-PI3KC3/VPS34 complex (Itakura et al., 2008; Liang et al., 2006; Liang et al., 2008), this regulatory function appears mechanistically distinct. Silencing core autophagy genes, including *Beclin1* or *ATG7*, did not recapitulate the accumulation of NOTCH1ΔE seen with *UVRAG* loss in DLL4-stimulated HeLa cells (Figure 1G). Similarly, knockdown of *ATG5*, *ATG16*, or *Beclin 1* failed to disrupt UVRAG-mediated suppression of NOTCH1 activity in T-ALL cells (Figures 1H and 1I). These data indicate that UVRAG regulates NOTCH1 activity via an autophagy-independent mechanism. To define the degradation route, we tested the involvement of lysosomal vs. proteasomal pathways. UVRAG-mediated reduction of NOTCH1ΔE and NICD was unaffected by proteasome inhibition (MG132) but reversed by lysosomal inhibitors such as chloroquine (CQ) and NH4Cl in both HeLa and T-ALL cells (Figures 1J and S2K). CQ treatment also restored NOTCH1 reporter activity and target gene expression suppressed by UVRAG (Figures S2L and S2M). This lysosome-dependent effect was confirmed in SW480 colorectal cancer cells expressing WT NOTCH1, where UVRAG overexpression reduced JAG2-induced NOTCH1ΔE and NICD levels in a dose-dependent manner, and this suppression was reversed by lysosomal inhibition (Figures S2N-S2P). Together, these results demonstrate that UVRAG selectively targets membrane-tethered NOTCH1ΔE for lysosomal degradation in an autophagy-independent manner, thereby constraining NICD production and downstream signaling output across multiple cell types (Figure 1K).

### UVRAG Selectively Binds Membrane-tethered NOTCH1ΔE

To determine how UVRAG regulates NOTCH1ΔE stability, we examined its interaction with the NOTCH1 receptor. In the absence of ligand, when S2 cleavage is minimal, UVRAG did not associate with NOTCH1 (Figure 2A). Upon DLL4 stimulation, UVRAG selectively co-immunoprecipitated with the membrane-tethered intermediate NOTCH1ΔE, but not with the cleaved NICD (Figure 2A). A similar pattern was observed in CUTLL1 T-ALL cells, which express a ligand-independent, constitutively active *NOTCH1* mutant: UVRAG robustly associated with endogenous NOTCH1ΔE but not NICD (Figure 2B). In co-expression assays, UVRAG bound to NOTCH1ΔE but failed to interact with NICD or NICD variants lacking the PEST domain (NICDΔ82 and NICDΔ162) (Figures S3A and S3B). Treatment with the γ-secretase inhibitor (GSI) DAPT, which blocks NICD production, did not disrupt UVRAG-NOTCH1ΔE binding (Figure S3C), indicating that UVRAG recognizes the membrane-tethered intermediate irrespective of downstream cleavage.

**Figure 2.**
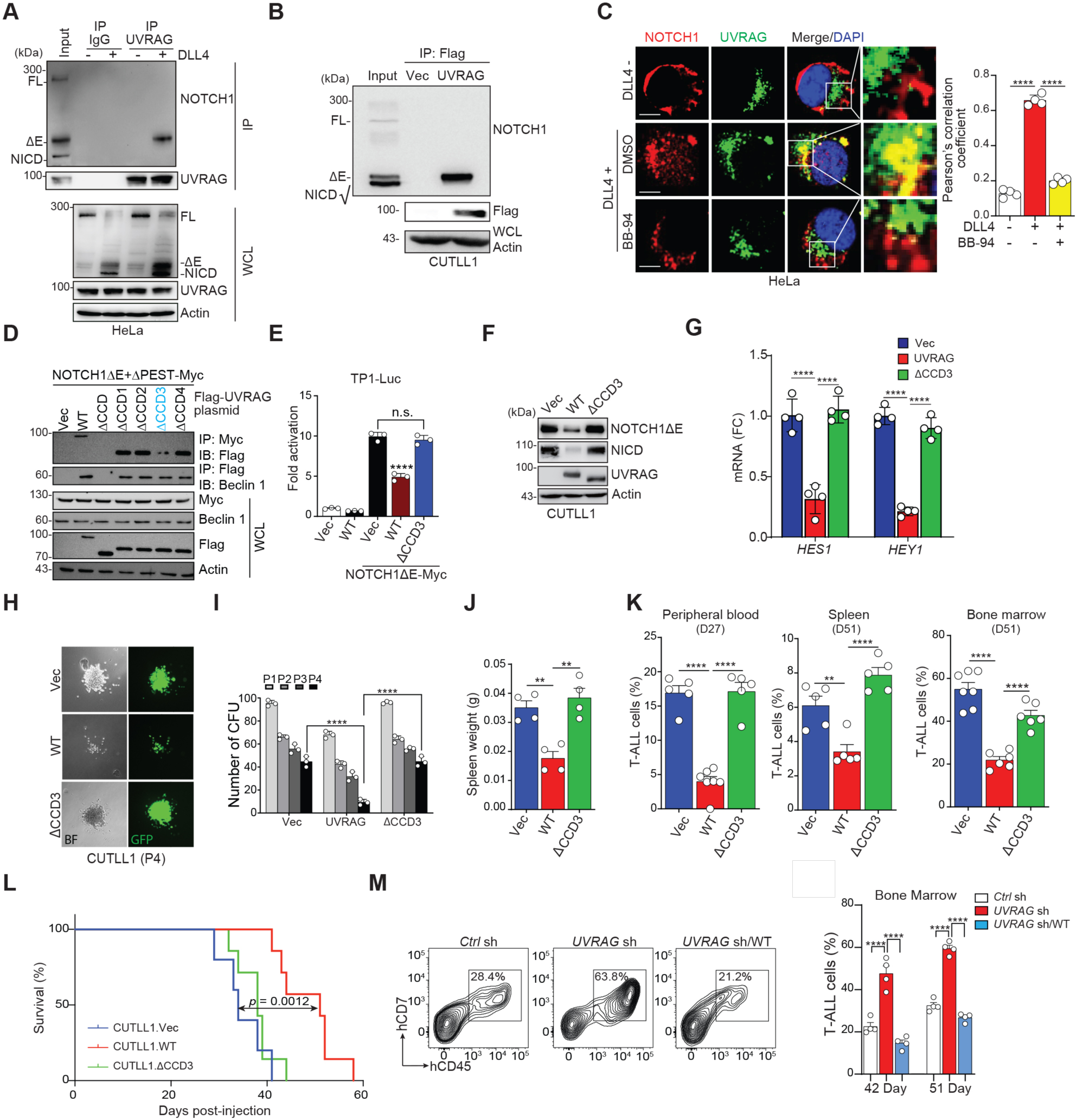
UVRAG Selectively Binds Membrane-bound NOTCH1 ΔE and Suppresses T-ALL. (A) Immunoprecipitation (IP) of NOTCH1 with UVRAG in HeLa cells, with or without DLL4 stimulation. FL, full-length. (B) IP of NOTCH1 with UVRAG in CUTLL1 cells expressing empty vector (Vec) or Flag-UVRAG. (C) Confocal imaging showing colocalization of endogenous NOTCH1 (red) and UVRAG (green) in DLL4-treated HeLa cells, with or without batimastat (BB-94) treatment. Nuclei stained with DAPI (blue). UVRAG-NOTCH1 colocalization quantified using Pearson’s correlation coefficient (n = 50 cells per group, 3 independent experiments). Scale bars, 20 mm. (D) Co-IP of UVRAG with NOTCH1ΔE or Beclin1 in HEK293T cells co-transfected with NOTCH1ΔE+ΔPEST-Myc and indicated UVRAG constructs. See also Figure S3H-J for UVRAG domain mapping. (E) TP1-luciferase activity in HEK293T cells co-transfected with Vec, UVRAG WT, or ΔCCD3 and NOTCH1ΔE-Myc. (F) IB of indicated proteins in CUTLL1 cells expressing Vec, UVRAG WT or ΔCCD3. (G) RT-qPCR of *HES1* and *HEY1* transcripts in cells from (F). n = 4. (H) Representative bright-field (BF) and GFP images of colony formation after 4 passages (P4) with CUTLL1 cells expressing Vec, UVRAG WT, or ΔCCD3 along with GFP. (I) Quantification of colony-forming units (CFU) from cells in (H). (J) Spleen weights from mice transplanted with CUTLL1 cells expressing Vec, UVRAG WT, or ΔCCD3 at day 51 post-injection (n = 4). (K) Quantification of T-ALL cells (hCD45^+^hCD7^+^) in peripheral blood (left), spleen (middle), and bone marrow (right) from mice in (J) at indicated time points (n = 5~6 per group). (L) Kaplan-Meier survival curves for mice transplanted as described in (J) (n = 5~6 per group). (M) Flow cytometry plots of bone marrow from mice transplanted with CUTLL1 cells expressing *Ctrl* sh or *UVRAG* shRNA complemented with Vec (*UVRAG* sh) or WT UVRAG (*UVRAG* sh/WT), analyzed for human CD45/CD7 at day 51 post-injection (n = 5). See also Figure S4C for UVRAG expression. (N) Quantification of T-ALL cells (hCD45^+^hCD7^+^) in bone marrow from mice in (M) at day 42 and 51 post-injection (n = 7). See also Figure S4E and S4F for leukemia burden. Data in (A-D), (F), (H), and (M) are representative of three independent experiments. Data in (C), (E), (G), (I-K), and (N) represent mean ± s.d. from biologically independent samples (n ≥ 3), analyzed by one-way ANOVA with Tukey’s *post hoc* test. Data in (L) analyzed using log-rank (mantel-cox) test. **, *p* < 0.01; ***, *p* < 0.001; ****, *p* < 0.0001; ns, not significant. See also Figure S3 and S4.

Because ligand binding can induce NOTCH1 endocytosis—a step required for activation (Chapman et al., 2016; Yamamoto et al., 2010)—we assessed UVRAG localization relative to NOTCH1 in HeLa cells. At baseline, NOTCH1 localized to the plasma membrane with minimal colocalization with UVRAG. Following DLL4 stimulation, NOTCH1 redistributed to cytoplasmic vesicles and extensively colocalized with endogenous UVRAG (Figure 2C). This colocalization was dependent on endocytosis, as *RAB5* knockdown abolished UVRAG-NOTCH1 overlap (Figure S3D). Blocking S2 cleavage with the metalloprotease inhibitor batimastat (BB-94) (van Tetering et al., 2009) prevented NOTCH1ΔE formation without disrupting ligand-induced endocytosis; under these conditions, UVRAG no longer colocalized with internalized NOTCH1, suggesting UVRAG specifically recognizes NOTCH1ΔE (Figure 2C). Co-localization with EEA1 (early endosomes) and LAMP1 (late endosomes/lysosomes) further confirmed that UVRAG-NOTCH1ΔE interactions occur within endosomal compartments (Figure S3E).

To define the domains mediating this interaction, we mapped UVRAG-binding regions within NOTCH1 using truncation mutants. In HEK293T cells, the transmembrane (TM) plus RAM (TM+R) domain of NOTCH1ΔE was both necessary and sufficient for UVRAG binding, whereas deletion of either domain (ΔR or ΔTM) abolished the interaction (Figures S3F and S3G). Complementary mapping of UVRAG revealed that its coiled-coil domain (CCD) was required for binding to NOTCH1ΔE (Figures S3H-J), with amino acids 235-252 (CCD3 region) identified as critical (Figures 2D and S3H). Because UVRAG’s CCD also binds Beclin 1 and mediate autophagy (Liang et al., 2006), we assessed whether CCD3 is functionally separable. Unlike the full CCD deletion (ΔCCD), the CCD3 deletion mutant (ΔCCD3) retained Beclin 1 binding (Figure 2D) and preserved autophagy activity, as evidenced by enhanced autophagosome-associated LC3 (LC3-II) conversion and reduced p62 (an autophagy substrate) levels (Mizushima et al., 2010) following Torin1 treatment (Figures S3K and S3L). These findings establish that UVRAG interacts with NOTCH1ΔE in endosomal compartments and this regulatory interface is distinct from UVRAG’s canonical autophagy function.

### UVRAG-NOTCH1ΔE Interaction is Required for NOTCH1ΔE Degradation and T-ALL Suppression

To determine whether UVRAG must physically interact with NOTCH1ΔE to suppress signaling, we examined the ΔCCD3 mutant, which lacks the NOTCH1ΔE-binding region but retains other UVRAG functions. In HEK293T cells, ΔCCD3 failed to inhibit NOTCH1ΔE-driven reporter activity, whereas WT UVRAG strongly suppressed it (Figure 2E). Similarly, in T-ALL cells expressing constitutively active *NOTCH1* mutants, WT UVRAG reduced levels of NOTCH1ΔE, NICD, and downstream target gene expression, while ΔCCD3 had no inhibitory effect (Figures 2F and 2G). We next asked whether this interaction affects access of NOTCH1ΔE to the γ-secretase complex. In CUTLL1 cells, NOTCH1 co-immunoprecipitated with core γ-secretase components, including presenilin 1 (PSEN1), nicastrin, APH-1, and PEN-2 (Yang et al., 2019), but these interactions were reduced by WT UVRAG, consistent with degradation of the precursor pool (Figure S4A). In contrast, ΔCCD3 did not alter γ-secretase binding, suggesting that direct engagement of NOTCH1ΔE by UVRAG is required for precursor elimination and suppression of downstream processing.

To assess the functional consequences of this interaction on NOTCH1-controlled cellular responses, we evaluated cell proliferation and self-renewal in CUTLL1 cells. Expression of WT UVRAG impaired leukemia cell proliferation and clonogenic capacity, with serial replating assays revealing a progressive decline in colony formation and near-complete exhaustion by passages 4 (P4) (Figures S4B, 2H, and 2I). The ΔCCD3 mutant did not affect these phenotypes. *In vivo*, CUTLL1 cells expressing either WT UVRAG or ΔCCD3 were transplanted into NSG mice. WT UVRAG markedly reduced leukemic burden, as evidenced by decreased spleen weight, reduced leukemic infiltration of bone marrow (BM), spleen, and peripheral blood, and prolonged survival (Figures 2J-L). These effects were lost in mice engrafted with ΔCCD3-expressing cells, highlighting the requirement for UVRAG-NOTCH1ΔE binding *in vivo*.

To determine whether endogenous UVRAG restrains NOTCH1-driven leukemia growth, we depleted *UVRAG* in CUTLL1 cells and transplanted them into NSG mice. *UVRAG* loss accelerated disease progression, resulting in increased leukemic infiltration, elevated mitotic index, and splenomegaly (Figures 2M, 2N, and S4C-F). Reintroduction of shRNA-resistant WT UVRAG restored suppression of these phenotypes, confirming specificity (Figures 2M, 2N, S4C-F). Notably, treatment with the γ-secretase inhibitor DAPT, but not the autophagy activator Torin1, rescued the effects of *UVRAG* loss (Figures S4G and S4H), supporting a signaling-specific, autophagy-independent mechanism. Together, these findings establish that the UVRAG-NOTCH1 interaction is required for limiting downstream signaling, cellular fitness, and in vivo progression driven by dysregulated NOTCH1 activity (Figure S4I).

### UVRAG Promotes K27-linked Ubiquitination of NOTCH1ΔE

To elucidate how UVRAG facilitates degradation of NOTCH1ΔE, we examined whether it regulates its ubiquitination—a key signal for lysosomal targeting of membrane proteins (Piper and Lehner, 2011). In HEK293T cells, WT UVRAG robustly promoted NOTCH1ΔE ubiquitination, whereas the ΔCCD3 mutant, which cannot bind NOTCH1, had no effect (Figure S5A). Conversely, *UVRAG* knockdown reduced NOTCH1ΔE ubiquitination and stabilized the protein; both effects were reversed by re-expression of shRNA-resistant WT UVRAG (Figure S5B). Similar results were observed in CUTLL1 and DND41 T-ALL cells, as well as in ligand-stimulated HeLa and SW480 cells, where UVRAG enhances ubiquitination of endogenous NOTCH1ΔE in a CCD3-dependent manner (Figures 3A, S5C, and S5D). Conversely, *UVRAG* depletion consistently led to NOTCH1ΔE accumulation and loss of ubiquitination (Figures 3B and S5E). These effects were independent of autophagy, as *Beclin 1* knockdown had no impact, and Beclin 1 did not co-immunoprecipitate with NOTCH1ΔE (Figure S5F). These results indicate that UVRAG promotes lysosomal turnover of NOTCH1ΔE through direct, autophagy-independent ubiquitination.

**Figure 3.**
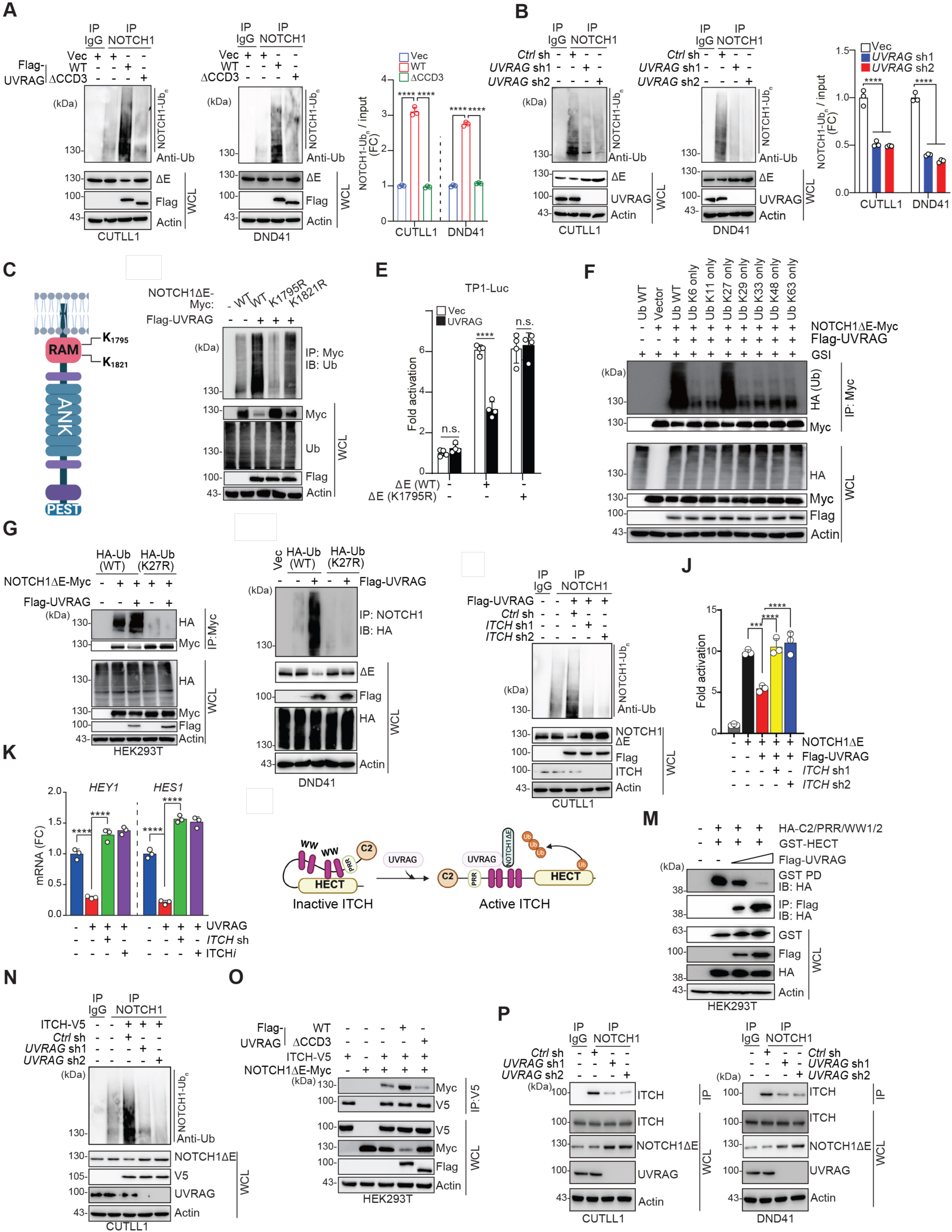
UVRAG Cooperates with ITCH to Promote K27-linked Ubiquitination of NOTCH1ΔE. (A and B) IP of NOTCH1 ubiquitination in CUTLL1 and DND41 cells expressing Vec, Flag-UVRAG WT or ΔCCD3 (A), or expressing *Ctrl* or *UVRAG*-specific shRNAs (sh1 and sh2) (B). Densitometric quantification of ubiquitinated NOTCH1 normalized to input (right; n = 3). IgG, negative control. (C) Schematic of ubiquitinated lysines in NOTCH1ΔE. RAM, RBPJk-associated molecule; ANK, ankyrin repeats; PEST, proline (P), glutamic acid (E), serine (S), and threonine (T) enriched region. See also Figure S5G-I. (D) Ubiquitination of Myc-tagged NOTCH1ΔE WT, K1795R, or K1821R mutant in HEK293T cells transfected with indicated plasmids. (E) TP1-luciferase activity in HEK293T cells co-transfected with WT or K1795R NOTCH1ΔE and Flag-UVRAG. n = 4. (F) Ubiquitination of NOTCH1ΔE-Myc in HEK293T cells expressing Flag-UVRAG and HA-Ub WT or mutants, treated with Compound E (GSI). (G and H) Ubiquitination of NOTCH1ΔE-Myc in HEK293T (G) and of endogenous NOTCH1 in DND41 (H) cells expressing Flag-UVRAG with HA-Ub WT or K27R. (I) Ubiquitination of NOTCH1 in CUTLL1 cells expressing Flag-UVRAG and *Ctrl* or *ITCH*-specific shRNAs. (J) CBF1-luciferase activity in HEK293T cells transfected with NOTCH1ΔE and Flag-UVRAG, expressing *Ctrl* or *ITCH* shRNAs (n = 3). (K) RT-qPCR of *HEY1* and *HES1* transcripts in CUTLL1 cells expressing Flag-UVRAG, treated with *ITCH* shRNA or ITCH inhibitor (ITCH*i*) Clomipramine (10 mM). n = 3. (L) Model depicting UVRAG-mediated relief of auto-inhibitory conformation in ITCH E3-ligase to facilitate NOTCH1ΔE ubiquitination. (M) GST pulldown (PD) assay showing binding between HA-tagged N-terminal domain (C2+PRR+WW1/2) and HECT domain of ITCH disrupted by UVRAG. (N) Ubiquitination of NOTCH1 in CUTLL1 cells expressing ITCH-V5 and *Ctrl* or *UVRAG* shRNAs. (O and P) Co-IP of NOTCH1ΔE with ITCH in HEK293T cells transfected with Vec or UVRAG (WT or ΔCCD3) (O), and of endogenous NOTCH1 with ITCH in cells expressing *Ctrl* or *UVRAG* shRNAs (P). Data in (A and B), (D), (F-I), and (M-P) are representative of three independent experiments. Data in (A), (B), (E), (J), and (K) represent mean ± s.d. from biologically independent samples, analyzed by one-way ANOVA with Tukey’s *post hoc* test. **, *p* < 0.01; ****, *p* < 0.0001; ns, not significant. See also Figure S5 and S6.

To identify the relevant modification sites of NOTCH1ΔE, we performed mass spectrometry following affinity purification of NOTCH1ΔE co-expressed with UVRAG in HEK293T cells treated with DAPT to prevent NICD production. Two conserved lysines in the RAM domain, K1795 and K1821, emerged as candidate ubiquitin acceptor sites (Figures 3C and S5G-I). Substitution of K1795 with arginine (K1795R) abolished UVRAG-dependent ubiquitination and stabilized NOTCH1ΔE, whereas the K1821R mutation had no effect (Figure 3D). The K1795R mutant also exhibited increased protein half-life and was refractory to *UVRAG* depletion (Figures S5J and S5K). Despite this stabilization, K1795R retained transcriptional competence—binding RBP-Jκ and activating downstream targets similarly to WT NOTCH1ΔE (Figures 3E, S5L, and S5M). However, it was resistant to UVRAG-mediated suppression (Figures 3E, S5M, and S5N), identifying K1795 as a functionally critical site for UVRAG-directed turnover.

To identify the ubiquitin linkage type, we co-expressed NOTCH1ΔE and Flag-UVRAG with a panel of HA-tagged ubiquitin (Ub) mutants, each retaining only a single lysine residue while all others were mutated, in GSI-treated HEK293T cells (Figure 3F). Among all variants, only K27-only Ub supported robust NOTCH1ΔE ubiquitination, matching levels seen with WT Ub (Figure 3F). This contrasts with canonical K48-linked ubiquitination of NICD by FBXW7 and other E3 ligases (Ma et al., 2023; Yeh et al., 2016). Ubiquitin restriction assay using chain-specific deubiquitinating enzymes (DUB) (Mevissen et al., 2013) further supported this specificity: YOD1 (K27/K29/K33-specific) and USP21 (broad spectrum) reduced NOTCH1ΔE ubiquitination, while DUBs targeting K48 (OTUB1), K63 (OTUB2), or K29/K33 (TRABID) had no effect (Figure S5O). Moreover, expression of a K27R Ub mutant blocked both ubiquitination and degradation of NOTCH1ΔE by UVRAG, whereas the K29R mutant did not (Figures 3G and S5P), suggesting that prior reports of K29-dependent regulation of non-activate NOTCH1 (Chastagner et al., 2008) do not extend to this signaling-committed intermediate. Likewise, in DND41 cells, HA-tagged WT Ub, but not K27R, supported endogenous NOTCH1ΔE ubiquitination (Figure 3H). These results establish K27-linked ubiquitination at lysine 1795 in the RAM domain as a critical signal directing NOTCH1ΔE to lysosomal degradation and limiting downstream signaling.

### UVRAG Activates ITCH to Catalyze NOTCH1ΔE Ubiquitination and Degradation

To identify the E3 ligase responsible for UVRAG-dependent ubiquitination of NOTCH1ΔE, we performed mass spectrometry on UVRAG-associated protein complexes in CUTLL1 cells. The HECT-type E3 ligase ITCH emerged as the top candidate (Figure S6A). While ITCH has been implicated in Notch signaling in *Drosophila* and in T cell regulation in mammals (Rathinam et al., 2011), its direct role in mammalian NOTCH1 degradation has not been established. We confirmed the interaction between UVRAG and ITCH through dose-dependent binding of recombinant proteins *in vitro* (Figure S6B). Functionally, ITCH was required for UVRAG-driven ubiquitination and degradation of NOTCH1ΔE. *ITCH* knockdown using two independent shRNAs attenuated UVRAG-induced NOTCH1ΔE ubiquitination and stabilized the protein in both T-ALL cells with *NOTCH1* mutations and ligand-stimulated SW480 cells expressing WT NOTCH1 (Figures 3I and S6C). Similarly, ITCH inhibition with clomipramine (Rossi et al., 2014) blocked UVRAG-mediated suppression of NOTCH1 signaling (Figures 3J and 3K). Knockdown of other factors implicated in NOTCH1 turnover, including *NEDD4* and *NUMB* (Kandachar and Roegiers, 2012; Wilkin et al., 2004), had no comparable effect (Figure S6D), highlighting the specificity of the UVRAG-ITCH axis. Consistent with UVRAG’s role in promoting K27-linked ubiquitination, mutation of lysine 27 on ubiquitin (K27R) abolished ITCH-dependent modification of NOTCH1ΔE (Figures S6E and S6F). In T-ALL patient datasets, ITCH expression inversely correlated with NOTCH1 target gene signatures, whereas no such relationship was observed for NEDD4 or NUMB (Figure S6G-I).

We next investigated how UVRAG regulates ITCH activity. Domain mapping in HEK293T cells revealed that both the proline-rich region (PRR) and WW1/2 domains of ITCH were each sufficient for UVRAG binding (Figures S6J and S6K). These domains are known to mediate intramolecular interactions with the C-terminal HECT domain, maintaining ITCH in an auto-inhibited state (Zhu et al., 2017). We hypothesized that UVRAG relieves this inhibition by competitively engaging the regulatory domains (Figure 3L). Co-immunoprecipitation confirmed strong binding between ITCH’s PRR-WW1/2 and HECT domains, which was diminished upon UVRAG expression (Figure 3L). *In vitro* ubiquitination assays further corroborated this mechanism: ITCH alone exhibited minimal activity toward NOTCH1, but addition of UVRAG enhanced ITCH-dependent ubiquitination in a dose-responsive manner (Figure S6L). In UVRAG-deficient T-ALL and SW480 cells, ITCH failed to promote NOTCH1ΔE ubiquitination or degradation (Figures 3N and S6M). Expression of WT UVRAG, but not ΔCCD3, restored ITCH association with NOTCH1ΔE and enhanced its ubiquitination, even as total NOTCH1ΔE levels declined (Figure 3O). Conversely, *UVRAG* depletion impaired ITCH-NOTCH1ΔE complex formation and led to accumulation of the substrate (Figure 3P). These results together identify UVRAG as a co-activator of ITCH that relieves autoinhibition and licenses K27-linked ubiquitination of membrane-tethered NOTCH1ΔE, thereby coupling substrate recognition to its selective lysosomal degradation (Figure S6N).

### ESCRT-Mediated Endo-lysosomal Sorting and Degradation of K27-Ubiquitinated NOTCH1ΔE

Membrane proteins marked for lysosomal degradation are typically routed through the ESCRT (endosomal sorting complexes required for transport) pathway, which facilitates their incorporation into intraluminal vesicles of multi-vesicular bodies (MVBs) (Henne et al., 2011; Hurley and Emr, 2006). In *Drosophila*, ESCRT deficiency leads to hyperactivation of Notch signaling and tissue overgrowth (Vaccari and Bilder, 2005), but whether this pathway regulates the degradation of activated, ubiquitinated NOTCH1ΔE in mammalian cells remains unclear. To test this, we depleted representative subunits from each ESCRT complex—*HRS* (ESCRT-0), *TSG101* (ESCRT-I), *VPS25* (ESCRT-II), *VPS24* (ESCRT-III)—in CUTLL1 cells stably expressing UVRAG. Loss of ESCRT function resulted in accumulation of ubiquitinated NOTCH1, increased NOTCH1ΔE and NICD levels, and loss of UVRAG-mediated suppression of signaling (Figure 4A-C). Correspondingly, UVRAG-dependent repression of reporter activity and target gene expression was abolished in *ESCRT*-deficient cells (Figures 4D and S7A). In contrast, knockdown of HOPS (homotypic fusion and protein sorting) complex components (*VPS16*, *VPS39*, or *VPS41*), which function in endolysosomal fusion and have been linked to activation—not repression—of NOTCH signaling (Wilkin et al., 2008), had no effect (Figure 4A-D and S7A). These results indicate that UVRAG-dependent degradation of NOTCH1ΔE proceeds specifically through the ESCRT pathway. Phenotypically, *ESCRT* loss phenocopied *UVRAG* depletion, leading to increased NOTCH1 target gene expression in both T-ALL and ligand-stimulated SW480 cells (Figures S7B and S7C). Functionally, *ESCRT* knockdown attenuated UVRAG’s ability to suppress colony formation in CUTLL1 cells (Figure 4E), suggesting that ESCRT acts downstream of UVRAG to constrain NOTCH1-driven growth. Confocal imaging in HeLa cells revealed that *ESCRT* depletion impaired colocalization of NOTCH1ΔE with the lysosomal marker LAMP1, consistent with defective trafficking (Figures 4F and 4G). Critically, this dependence on ESCRT was contingent on upstream ubiquitination. The NOTCH1ΔE K1795R mutant, which is resistant to K27-linked modification, exhibited elevated reporter activity (compared to WT) and was unresponsive to *ESCRT* depletion (Figure S7D). Similarly, *ESCRT* loss had no additive effect in *UVRAG-* or *ITCH-*deficient cells, where ubiquitination is already impaired (Figure S7E). Together, these results support a model in which K27-linked ubiquitination of NOTCH1ΔE by the UVRAG-ITCH complex licenses its recognition and sorting by the ESCRT machinery for lysosomal degradation.

**Figure 4.**
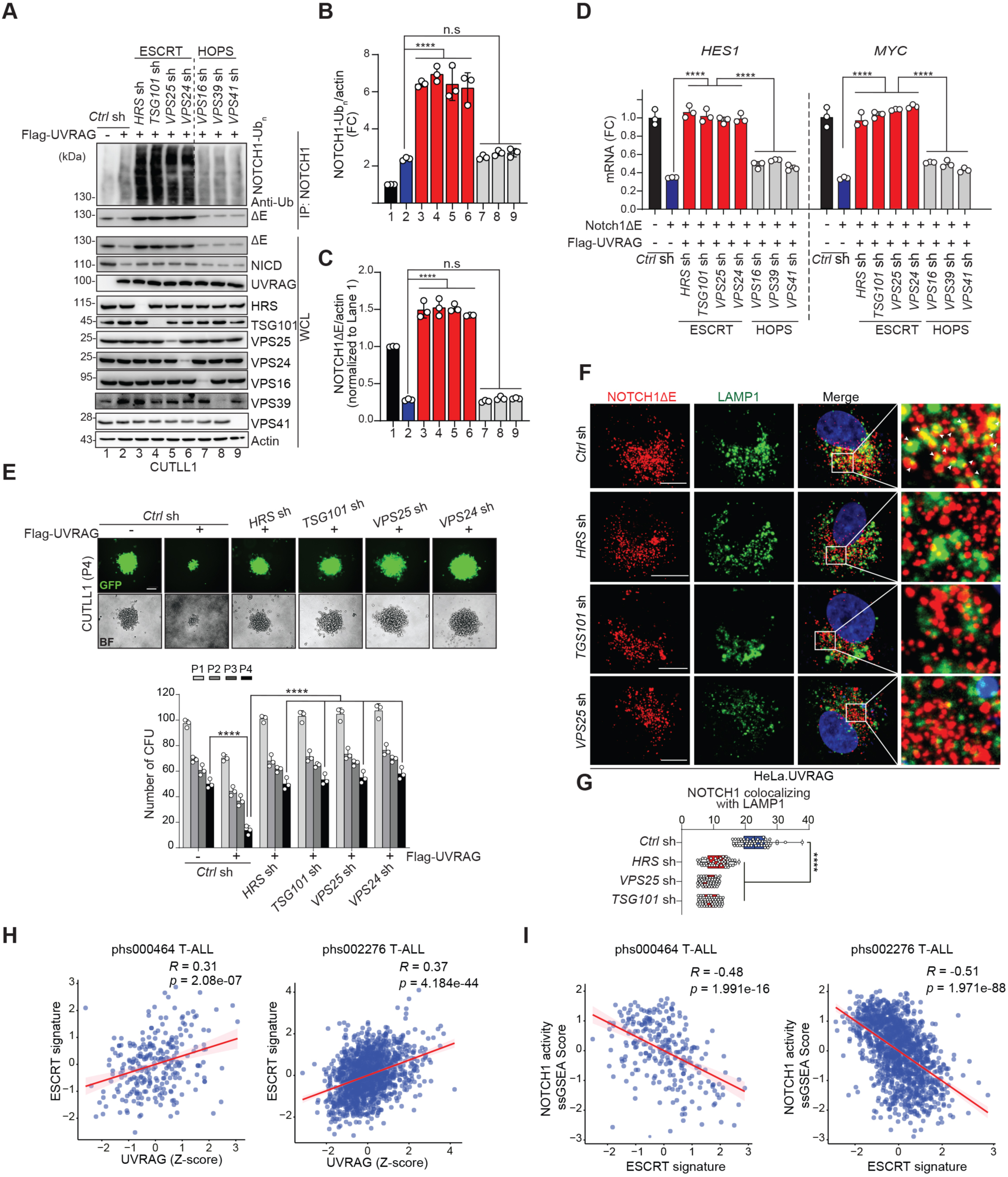
ESCRT-mediated Endo-lysosomal Degradation of K27-Ubiquitinated NOTCH1ΔE. (A) Ubiquitination of NOTCH1 immunoprecipitated in CUTLL1 cells expressing Flag-UVRAG and *Ctrl* shRNA or indicated gene-specific shRNAs. IgG, negative control. (B and C) Densitometric quantification of ubiquitinated NOTCH1 (NOTCH1-Ub_n_; B) and NOTCH1ΔE (C) normalized to actin (*n* = 3). (D) RT-qPCR analysis of *HES1* and *MYC* transcripts in cells from (A). n = 3. See also Figure S7A for reporter activity. (E) Representative GFP and BF images (top) showing colony formation after P4 in CUTLL1 cells expressing Vec or UVRAG and indicated shRNAs; CFU quantified (bottom; n = 3). (F) Confocal micrographs of NOTCH1ΔE-Myc (red) colocalization with LAMP1 (green) in HeLa.UVRAG cells expressing NOTCH1ΔE-Myc and *Ctrl* shRNA or indicated shRNAs. Insets highlight reduced co-distribution of NOTCH1ΔE with LAMP1 upon *ESCRT* deficiency. Arrowheads indicate colocalization (n = 50 cells per group, 3 independent experiments). Scale bars, 10 mm. (G) Quantification of the fraction of NOTCH1ΔE co-localized with LAMP1 in (F). (H) Scatter plot correlating *UVRAG* expression (Z-score normalized) with *ESCRT* signature genes (*STAM*, *TSG101*, *VPS37A*, *CHMP3*, *VPS4B*) expression in T-ALL samples (dbGaP phs000464 T-ALL, n = 264; phs002276, n = 1335). (I) Scatter plot correlating ESCRT gene signature expression (Z-score normalized) with the ssGSEA scores of NOTCH1 signature (Palomero et al., 2006b) from two T-ALL datasets in (H). Data in (A), (E), and (F) are representative of three independent experiments. Data in (B-D) and (G) represent mean ± s.d. from biologically independent samples, analyzed by one-way ANOVA with Tukey’s *post hoc* test. ****, *p* < 0.0001. See also Figure S7.

To assess clinical relevance, we analyzed The Cancer Genome Atlas (TCGA) datasets across multiple tumor types. A five-gene ESCRT signature (*TSG101, VPS37A, STAM, CHMP3,* and *VPS4B*) was co-expressed with *UVRAG* and *ITCH* and inversely correlated with canonical NOTCH1 target genes *HES1* and *DTX1* (Figures S7F and S7G). High expression of this *ESCRT* module was associated with improved disease stable survival (DSS) in cancers characterized by aberrant NOTCH1 signaling, including kidney renal clear cell carcinoma (KIRC) and low-grade glioma (LGG) (Figure S7H), but not in NOTCH1-independent malignancies (Figure S7I). In T-ALL, ESCRT gene expression positively correlated with *UVRAG* and inversely with NOTCH1 transcriptional output (Figures 4H and 4I). Across the 15 recently classified molecular subtypes of T-ALL (Polonen et al., 2024), those with elevated NOTCH1 activity, such as TLX^+^, exhibited significantly reduced expression of the UVRAG-ITCH-ESCRT module (Figure S7J). Together, these findings define a conserved, post-translational surveillance mechanism in which UVRAG and ITCH direct K27-linked ubiquitination of NOTCH1ΔE, targeting it for ESCRT-mediated lysosomal degradation and limiting pathological NOTCH1 signaling across multiple tissue contexts.

### UVRAG Restrains NOTCH1-driven Leukemogenesis and Leukemia-initiating Cell Function *in vivo*

To assess whether UVRAG-dependent degradation of NOTCH1ΔE constrains NOTCH1-driven malignancy *in vivo*, we focused on T-ALL, where aberrant NOTCH1 activation is a central driver of disease (Kourtis et al., 2018). Analysis of primary T-ALL patient samples revealed significantly reduced UVRAG expression compared to normal T cells, accompanied by an inverse correlation with NICD abundance (Figures 5A and 5B). Moreover, high *UVRAG* expression was associated with decreased leukemic burden (Figure 5C) and improved clinical outcomes in NOTCH1-driven T-ALL (Figure 5D). Based on these observations, we used a genetically engineered mouse model to directly test causality. We generated a doxycycline (Dox)-inducible *Uvrag* mouse by crossing *TRE-Uvrag* with *ROSA26-rtTA2*M2* mice (Belteki et al., 2005), yielding a strain with robust, reversible UVRAG induction upon Dox treatment (Figures S8A-C). T-ALL was initiated by transducing *iUvrag* bone marrow progenitors with *NOTCH1*Δ*E-IRES-GFP*, followed by transplantation into lethally irradiated recipients (Kourtis et al., 2018) (Figure S8D). Upon leukemia establishment, Dox-induced *Uvrag* expression (On_Dox) significantly reduced leukemia burden and extended survival compared to untreated controls (No_Dox) (Figures 5E-G and S8E-G). As in human T-ALL, leukemic cells displayed low endogenous UVRAG, which was restored by Dox and correlated with decreased NICD levels (Figure S8H). Transcriptomic profiling of UVRAG-expressing leukemic cells showed downregulation of canonical NOTCH1 targets (*Hes1*, *Hes5*, *Dtx1*, *Gimap*, and *Myc*) and upregulation of pro-apoptotic genes (*Casp1*, *Casp4*, *Noxa*) (Figures 5H and S9A-C). Pathway analysis revealed suppression of *MYC* targets and NF-kB signaling (Figures 5I and S9C), mirroring transcriptomic changes observed in UVRAG-high patient samples (Figure S9D). UVRAG also promoted differentiation, as evidenced by increased expression of CD4^+^ and CD8^+^ single-positive (SP) markers, whereas control leukemias remained arrested at immature stages (Figure S9E), consistent with previous studies showing that excessive NOTCH1 activity impairs T cell maturation (De Keersmaecker et al., 2008; Izon et al., 2001). To determine whether UVRAG’s tumor-suppressive function depends on NOTCH1ΔE ubiquitination, we compared leukemias driven by WT or K1795R NOTCH1ΔE, the latter lacking the K27-linked ubiquitination site. K1795R-expressing leukemia were refractory to UVRAG induction and progressed more rapidly (Figure S9F), confirming that this post-translational modification is required for UVRAG-mediated control of NOTCH1 signaling *in vivo*.

**Figure 5.**
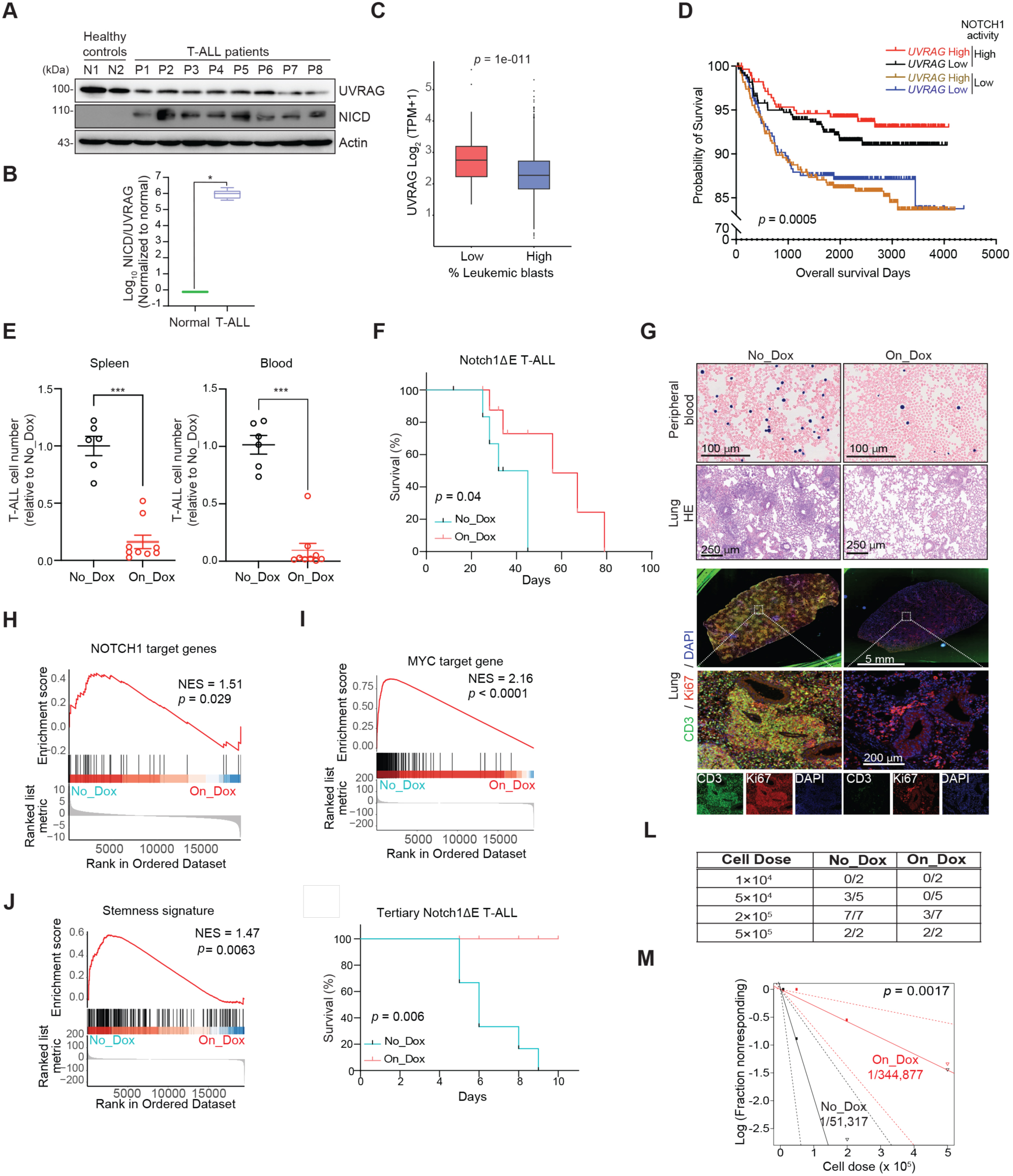
UVRAG Restrains NOTCH1-driven Leukemia Progression and Limits Leukemia-Initiating Cell Function *in vivo*. (A) Protein expression of UVRAG and NICD in T cells from healthy donors (N1, N2) and primary T-ALL patients bone marrow biopsies (P1-P8); representative immunoblots shown. (B) Densitometric quantification of NICD/UVRAG protein levels from (A); n = 3. (C) UVRAG expression in T-ALL samples with high (≥ 70%; n = 1144) and low (< 70%; n = 160) leukemic blast (%) for the T-ALL dataset (dbGaP phs002276). Note the inverse relationship between UVRAG expression and leukemic burden. (D) Kaplan-Meier survival curves comparing the overall survival between T-ALL patients with a high *vs.* low *UVRAG* expression and/or NOTCH1 activity signature (Palomero et al., 2006b), using the median as threshold, in T-ALL patient cohorts from dbGaP (accession number phs002276). The association of the four groups with overall survival was determined using Cox hazards proportional regression analysis. Log-rank *p*-value for trend is indicated. (E) Quantification of GFP^+^ i*Uvrag* T-ALL cells (transduced with Notch1ΔE) in indicated tissues 4 weeks post-treatment with Dox (On_Dox; n = 9) or without Dox (No_Dox; n = 6). See also Figure S8E and S8F. (F) Kaplan-Meier survival curve of recipient mice transplanted with i*Uvrag* BM transduced with NOTCH1ΔE and treated with or without Dox. n = 5. (G) Wright-Giemsa stained peripheral blood smears (top row), H&E-stained lung sections (2^nd^ row), and fluorescent confocal microscopy of lungs stained for CD3 (green), Ki67 (red), and DAPI (blue) harvested 4 weeks post-Dox treatment (3-5 rows). Scale bars as indicated. See also Figure S8G. (H-J) GSEA of NOTCH1 target gene signature (Palomero et al., 2006b) (H), MYC target gene signature (Liberzon et al., 2015) (I), and stemness signature (Ramalho-Santos et al., 2002) (J) suppressed in T-ALL with induced *Uvrag* expression (On_Dox). NES, normalized enrichment score. (K) Kaplan-Meier survival curves of tertiary recipients transplanted with BM from i*Uvrag* cells transduced with Notch1ΔE, treated with or without Dox. (L) Fraction of secondary recipient mice developing leukemia after transplantation with limiting dilution of GFP^+^ splenocytes from Dox-treated or untreated i*Uvrag* T-ALL mice. (M) Log-log plot and LIC frequency calculated *via* extreme limiting dilution analysis in Dox-treated (1/344,877; red) and untreated (1/51,317; black) mice. Boxplot data in (B and C): the center line indicates median, boxes indicate interquartile range (IQR, 25^th^-75^th^ percentiles), and whiskers extend to minimum and maximum values; analyzed using Mann-Whitney U test (B) or Student’s two-tailed *t* test (C). Data in (E) represent mean ± s.e.m. and were analyzed using the Mann-Whitney U test. Data in (D), (F), and (K) analyzed using log-rank (Mantel-Cox) test. *, *p* < 0.05; ****, *p* < 0.0001. See also Figure S8 and S9.

Given the role of NOTCH1 in sustaining leukemia-initiating cell (LIC) (Armstrong et al., 2009; Cox et al., 2004), we next examined whether UVRAG impacts LIC function. GSEA of leukemic cells from untreated mice revealed enrichment of stem cell-associated transcriptional programs (Ramalho-Santos et al., 2002) (Figure 5J). Serial transplantation demonstrated that UVRAG induction significantly impaired leukemia reconstitution in tertiary recipients (Figure 5K). In limiting dilution assays, UVRAG reduced leukemia-initiating potential: only 43% of mice injected with 2 x 10^5^ UVRAG-expressing cells developed leukemia compared to 100% of controls, and no disease developed from 5 x 10^4^ cells *versus* 60% in controls (Figure 5L). Estimated LIC frequency was reduced more than six-fold upon UVRAG induction (Figure 5M). Correspondingly, UVRAG-expressing leukemias downregulated LIC-associated gene signatures (Kourtis et al., 2018) (Figure S9G). These findings establish UVRAG as a critical regulator of oncogenic NOTCH1 signal termination *in vivo*. By promoting lysosomal degradation of the NOTCH1ΔE intermediate, UVRAG limits leukemic expansion and impairs the self-renewal capacity of LIC, defining a previously unrecognized post-translational checkpoint in NOTCH1-driven malignancy.

### UVRAG Sensitizes Cancer Cells to γ-secretase Inhibition

Given its role in promoting lysosomal degradation of membrane-tethered NOTCH1ΔE, we next asked whether UVRAG influences the cellular response to γ-secretase inhibitors (GSIs), which block NICD release and downstream signaling (Habets et al., 2019). Although GSIs show preclinical activity in T-ALL, their clinical use has been limited by dose-limiting gastrointestinal (GI) toxicity (Doody et al., 2013). Nonetheless, a subset of patients achieves durable responses (Knoechel et al., 2015; Papayannidis et al., 2015), prompting efforts to identify modulators of GSI sensitivity. We hypothesized that UVRAG, by depleting NOTCH1ΔE upstream of γ-secretase, could sensitize leukemic cells to GSI therapy. Consistent with this model, *UVRAG* knockdown reduced sensitivity to MRK-560—a PSEN1-selective GSI with reduced GI toxicities (Habets et al., 2019)—across multiple T-ALL cell lines, a phenotype that was recapitulated by *ITCH* depletion (Figures S10A and S10B). *NOTCH1* silencing in *UVRAG*-deficient cells re-sensitized them to MRK-560 (Figure S10C), implicating accumulated NOTCH1ΔE as a mediator of resistance. Conversely, ectopic expression of WT UVRAG enhanced MRK-560 sensitivity by nearly two orders of magnitude (Figure 6A), accompanied by reduced NICD abundance, MYC suppression, and induction of apoptosis (Figure 6B). This effect required NOTCH1 binding, as the ΔCCD3 mutant failed to enhance sensitivity. Notably, UVRAG expression rendered otherwise resistant RPMI-8402 cells responsive to MRK-560, suggesting that it can overcome intrinsic GSI resistance (Figure 6A).

**Figure 6.**
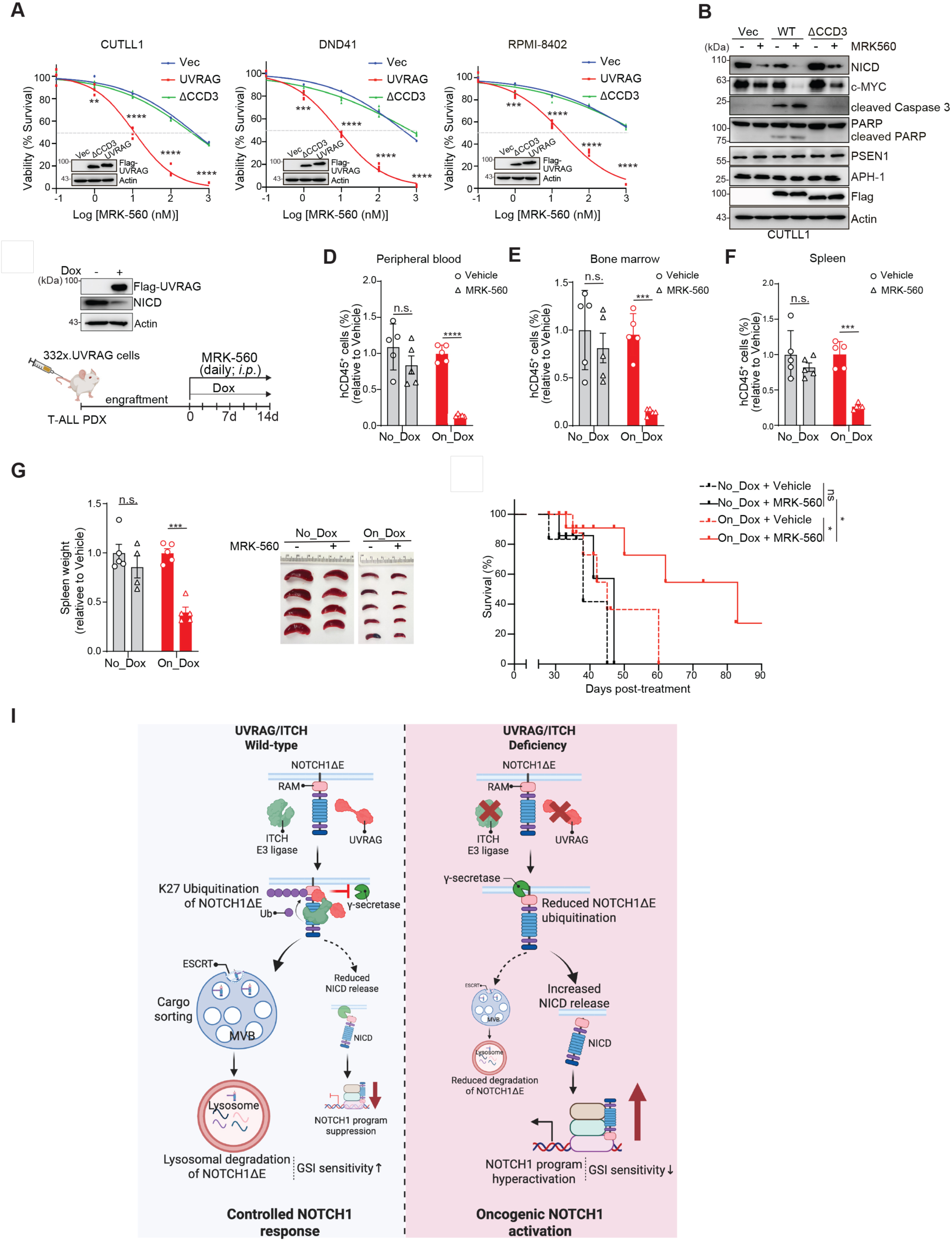
UVRAG Enhances T-ALL Sensitivity to γ-Secretase Inhibition by Depleting NOTCH1ΔE. (A) Cell survival assay of indicated T-ALL cell lines stably expressing Vec, UVRAG WT, or ΔCCD3 mutant after 7-day treatment with indicated concentrations of MRK-560 (n = 3 replicates for each condition per cell line). Immunoblots show Flag-UVRAG expression, with actin as loading control. (B) IB of indicated proteins in CUTLL1 cells from (A), with or without MRK-560 treatment (10 mM, 7 days). (C) Schematic overview of experiment design. NSG mice bearing primary T-ALL PDX 332x (*via* tail vein injection) were treated with or without Dox to induce Flag-UVRAG expression after disease establishment, followed by daily *i.p.* injections of MRK-560 (30 mmol kg^-1^) for 14 days. Immunoblots show Flag-UVRAG and NICD expression in PDX 332x cells (top). (D-G) Percentages of human CD45^+^ cells in peripheral blood (D), bone marrow (E), and spleen (F), and overall spleen weight (G) from Dox-treated (On_Dox) or untreated (No_Dox) mice transplanted with patient sample 332x and treated with DMSO or MRK-560 for 14 days. Fold changes comparing MRK-560 *vs*. vehicle within each group are shown (n = 5 per group). Representative spleen images shown (right). (H) Kaplan-Meier survival curves for xenotransplanted mice ± Dox treatment, administered Vehicle or MRK-560 as described in (C). (I) Proposed working model illustrating K27-linked ubiquitination checkpoint controlling membrane-bound NOTCH1ΔE *via* the UVRAG-ITCH-ESCRT pathway. UVRAG, together with ITCH E3 ligase, promotes K27-linked ubiquitination of NOTCH1ΔE, directing its lysosomal degradation and reducing NOTCH1 signaling upstream of γ-secretase cleavage. Dysregulation of this checkpoint enhances NOTCH1 signaling, promoting T-ALL progression and dampening therapeutic responsiveness. Data in (B) are representative of three independent experiments. Data in (A) and (D-G) represent mean ± s.e.m. analyzed by Student’s two-tailed *t* test for two-group comparisons or one-way ANOVA followed by Tukey’s *post hoc* test. **, *p* < 0.01; ***, *p* < 0.001; ****, *p* < 0.0001; ns, not significant. See also Figure S10.

To validate these findings *in vivo*, we xenografted NSG mice with CUTLL1 cells stably expressing vector, WT UVRAG, or ΔCCD3, and treated them with MRK-560 at two doses (Figure S10D). Mice bearing WT UVRAG-expressing cells showed significantly reduced leukemia burden and prolonged survival, even at lower GSI doses, whereas vector and ΔCCD3 groups were largely unresponsive (Figures S10E and S10F). No GI toxicity, including goblet cell hyperplasia, was observed (Figure S10G), consistent with the selective activity of MRK-560 (Habets et al., 2019). To test this strategy in a clinically relevant setting, we employed a T-ALL patient-derived xenograft (PDX) harboring an activating *NOTCH1* mutation (Figure 6C). Dox-induced expression of WT UVRAG in PDX cells reduced NICD levels (Figure 6C) and markedly enhanced MRK-560 efficacy, resulting in rapid leukemia regression (Figures 6D-G) and improved survival (Figure 6H). Collectively, these findings demonstrate that UVRAG-mediated degradation of NOTCH1ΔE sensitizes T-ALL cells to γ-secretase inhibition. By targeting the signaling-committed receptor intermediate upstream of NICD release, UVRAG enhances therapeutic response and provides a mechanistically distinct strategy to improve the efficacy of NOTCH1-directed therapies.

## DISCUSSION

This study defines a membrane-proximal degradation checkpoint that attenuates NOTCH1 signaling by selectively targeting the signaling-committed intermediate NOTCH1ΔE. We identify UVRAG as a scaffold and co-activator that engages the E3 ligase ITCH to catalyze K27-linked ubiquitination of NOTCH1ΔE at lysine 1795, directing its ESCRT-mediated trafficking to the lysosome. By limiting substrate availability for γ-secretase, this pathway operates upstream of NICD release and dampens the amplitude of downstream transcriptional responses (Figure 6I).

Although the proteolytic cascade of NOTCH activation has been well characterized, the regulatory logic upstream of γ-secretase remains incomplete. Our findings reveal that NOTCH1ΔE is not simply a passive precursor destined for activation, but instead a regulatory node subject to selective turnover. UVRAG binds NOTCH1ΔE at endosomes—whether generated by ligand stimulation or pathological activation—and facilitates its degradation independently of autophagy. This distinguishes our mechanism from earlier reports linking general autophagic flux to receptor turnover (Wu et al., 2016), and underscores a dedicated role for UVRAG in modulating activated NOTCH1 intermediates.

The mechanism by which UVRAG targets NOTCH1ΔE involves UVRAG recruitment and activation of the HECT-type E3 ligase ITCH; ITCH-mediated K27-linked ubiquitination of membrane-bound NOTCH1ΔE at K1795; and subsequent ESCRT-dependent endo-lysosomal trafficking and degradation. Although other E3 ubiquitin ligases, including NEDD4 (Sakata et al., 2004) and FBXW7 (Thompson et al., 2007), have been implicated in the regulation of NOTCH1, FBXW7 selectively targets nuclear NICD for proteasomal degradation (Thompson et al., 2007), while NEDD4—often recruited by the adaptor protein NUMB—has been shown to promote ubiquitination of un-liganded NOTCH1 receptor at the plasma membrane (Sakata et al., 2004). Notably, we found that neither NEDD4 nor NUMB expression inversely correlated with NOTCH gene signatures in T-ALL. We further found that depletion of *ITCH*, but not *NEDD4* nor *NUMB*, diminished NOTCH1 ubiquitination induced by UVRAG overexpression. Consistently, suppression of ITCH phenocopied the consequences of UVRAG loss, boosting NOTCH1 signaling by accrual of membranous NOTCH1ΔE. These data support that ITCH is a specific E3 ubiquitin ligase for activated NOTCH1ΔE, with UVRAG being a critical substrate adaptor for ITCH-mediated NOTCH1ΔE ubiquitination. These findings align with the previous work highlighting an association between ITCH deficiency and augmented NOTCH activation particularly in cells in the immune system (Matesic et al., 2006; Rathinam et al., 2011).

Previous studies suggest that HECT-type E3 ligases, including ITCH, generally adopt an auto-inhibited conformation through intramolecular interactions (Zhu et al., 2017). We show that UVRAG binding to the WW domains of ITCH relieves its autoinhibition, enabling formation of a ternary UVRAG-ITCH-NOTCH1ΔE complex. This interaction promotes site-specific K27-linked ubiquitination of NOTCH1ΔE at lysine 1795. This modification is functionally distinct from FBXW7-mediated K48-linked degradation of NICD (Yeh et al., 2016) or previously proposed K29-linked turnover of full-length NOTCH1 (Chastagner et al., 2008). Blocking K27-linked ubiquitination, either through *UVRAG* or *ITCH* loss or by introducing a K1795R mutation in NOTCH1, leads to accumulation of NOTCH1ΔE and heightened transcriptional output. These findings support a model in which K27-linked ubiquitination acts as a molecular signal for ESCRT engagement and lysosomal routing. Consistently, ESCRT disruption caused accumulation of ubiquitinated NOTCH1ΔE in endosomes and elevated NOTCH1 target gene expression, explaining prior genetic observations showing that ESCRT dysfunction results in ectopic NOTCH activity in *Drosophila* and mammals (Vaccari and Bilder, 2005; Yamamoto et al., 2010).

Our work supports the idea that NOTCH1 signaling is highly sensitive to dosage (Chiang et al., 2008; Izon et al., 2001). By regulating the turnover of NOTCH1ΔE, the UVRAG-ITCH-ESCRT axis acts as a rheostat that dampens signal intensity. In T-ALL, where this axis is frequently suppressed, elevated NOTCH1ΔE levels may sustain leukemic self-renewal and prevent differentiation. Indeed, restoring UVRAG expression reduced leukemic burden, depleted LICs, and induced transcriptional programs associated with T cell maturation. These effects were abolished when NOTCH1ΔE could not be ubiquitinated, highlighting the functional relevance of this degradation checkpoint.

More broadly, this work underscores the importance of kinetic checkpoints for tuning the amplitude of proteolytically driven signaling pathways, which lack reversibility once signaling is initiated. By introducing a mechanism that actively surveils and eliminates these intermediates, our study reframes endosomes as dynamic quality control hubs that tune irreversible signaling cascades. Notably, NOTCH1 plays critical roles in developmental patterning, tissue renewal, and stem cell maintenance across multiple epithelial and hematopoietic contexts. The ability to selectively degrade signaling-competent intermediates may represent a conserved strategy to calibrate signaling thresholds in progenitor niches, where both receptor availability and endocytic activity fluctuate. Whether similar degradation checkpoints operate on other membrane-tethered intermediates in different signaling pathways warrants further investigation.

Our findings also suggest a conceptual framework for therapeutic modulation of signal strength through selective degradation of active receptor intermediates. We show that enhancing UVRAG expression sensitizes cancer cells to γ-secretase inhibition by limiting the pool of substrate available for NICD production. This dual-targeting approach—limiting both substrate and catalytic release—offers a generalizable strategy for improving sensitivity and minimizing toxicity in pathways regulated by sequential proteolysis.

In sum, this study identifies a previously unrecognized endolysosomal checkpoint that regulates NOTCH1 signaling homeostasis upstream of nuclear response. By coupling K27-linked ubiquitination of NOTCH1ΔE to ESCRT-mediated degradation, UVRAG and ITCH establish a membrane-level control mechanism that safeguards signal fidelity. These findings expand our understanding of how ubiquitin linkages, endosomal trafficking, and receptor dynamics intersect to modulate pathway activation—and reveal a tunable axis for controlling signal intensity at the interface of membrane and nucleus.

## ACKNOWLEDGEMENTS

We appreciate the assistance from the members of the Liang laboratory who were involved in this work. We thank the Genomics, Flow Cytometry, Imaging, Bioinformatics, and Animal Facility at The Wistar Institute for their technical support. We are grateful to Ross Tomaino (Taplin Mass Spectrometry Facility, Department of Cell Biology, Harvard Medical School) for assistance with mass spectrometry analysis. We thank A. Hartley, Dr. W. Wells, and Dr. R. Subramanyan (Department of Cardiothoracic Surgery, Children’s Hospital Los Angeles) for their help in collecting thymic tissue. We acknowledge the Therapeutically Applicable Research to Generate Effective Treatments (TARGET) program for generating the dbGaP dataset used in this study. We are especially grateful to the participants and investigators involved in the TARGET initiative. The interpretations presented here are those of the authors and do not necessarily reflect the views of the original investigators or the NIH. All graphic illustrations were created using BioRender.com under an institutional license. B. Mansoori was supported in part by The Wistar Institute Training Program (T32 CA009171). I.A. was supported by the Vogelstein Family Foundation, the St. Baldrick’s Foundation and the NIH (P01 CA229086, R01 CA298153, and R01 CA252239). This work was supported by NIH awards R01 CA140964, R01 CA289624, and R01 CA262631 to C. Liang (PI).

## AUTHOR CONTRIBUTIONS

C.L. conceived and supervised the project, designed experiments, and secured funding. C.L. and C.P. developed the conceptual framework for the limiting dilution and thymus studies. I.A. contributed to the bone marrow transplantation T-ALL model. B.M., Y.S., and T.Z. performed most experiments. Z.Z., M.R., A.V.K., and N.A. carried out bioinformatics analyses. J.C. and Q.Z. assisted with imaging. S.L., J.L., C.P., and J.S.M. supported biochemical and animal experiments. B.M., Y.S., T.Z., Z.Z., M.R., R.M., and Q.L. analyzed data and assembled figures. C.L. wrote the original manuscript. B.M, M.E.M., M.M., I.A., C.P., C.L. reviewed and edited the manuscript. Y-K.H., G.B.W., Y.K., M.G., I.A., C.P., and C.L. provided resources.

## SUPPLEMENTAL INFORMATION

Figures S1-S10, Table S1 and S2

## STAR★METHODS

### KEY RESOURCES TABLE

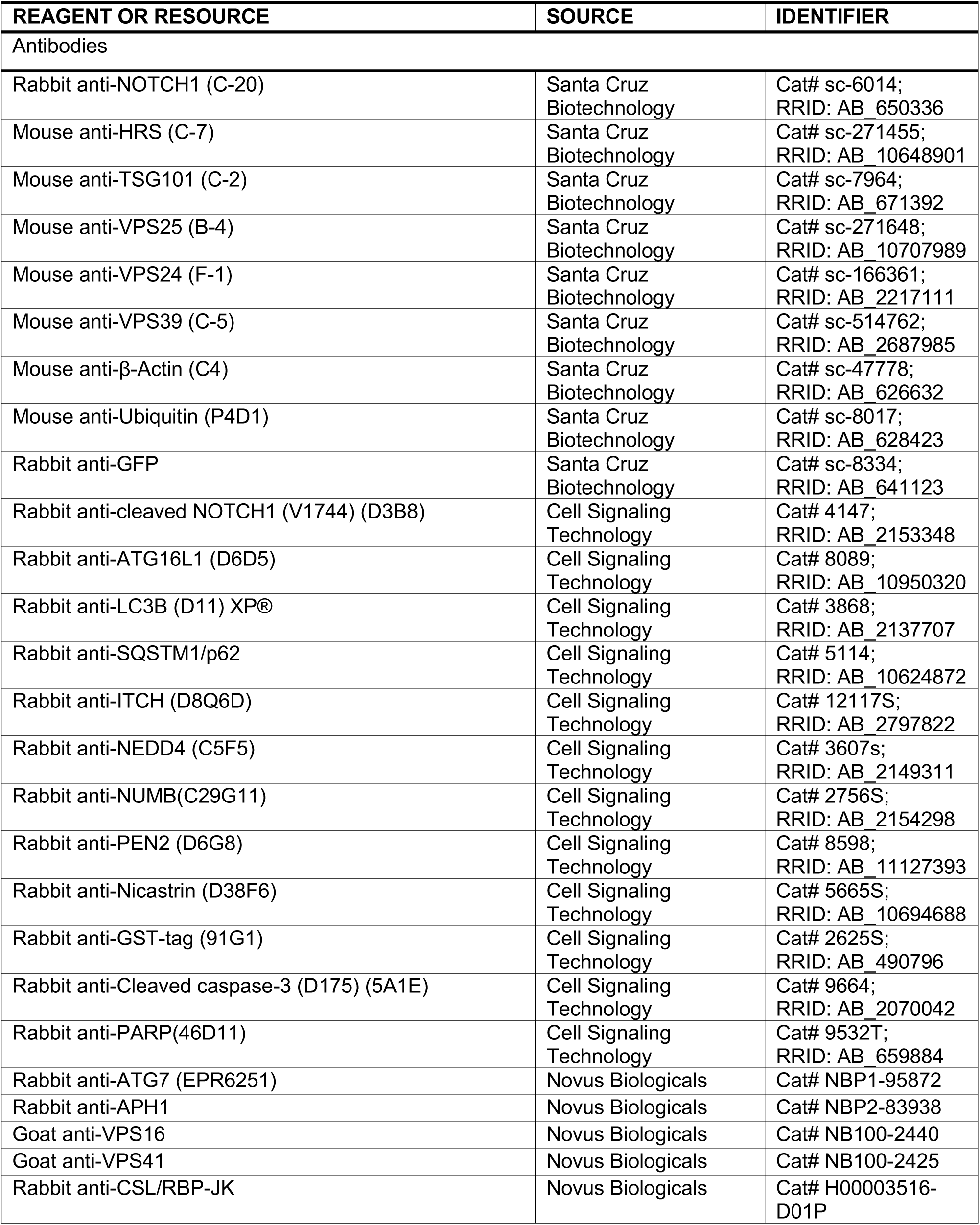
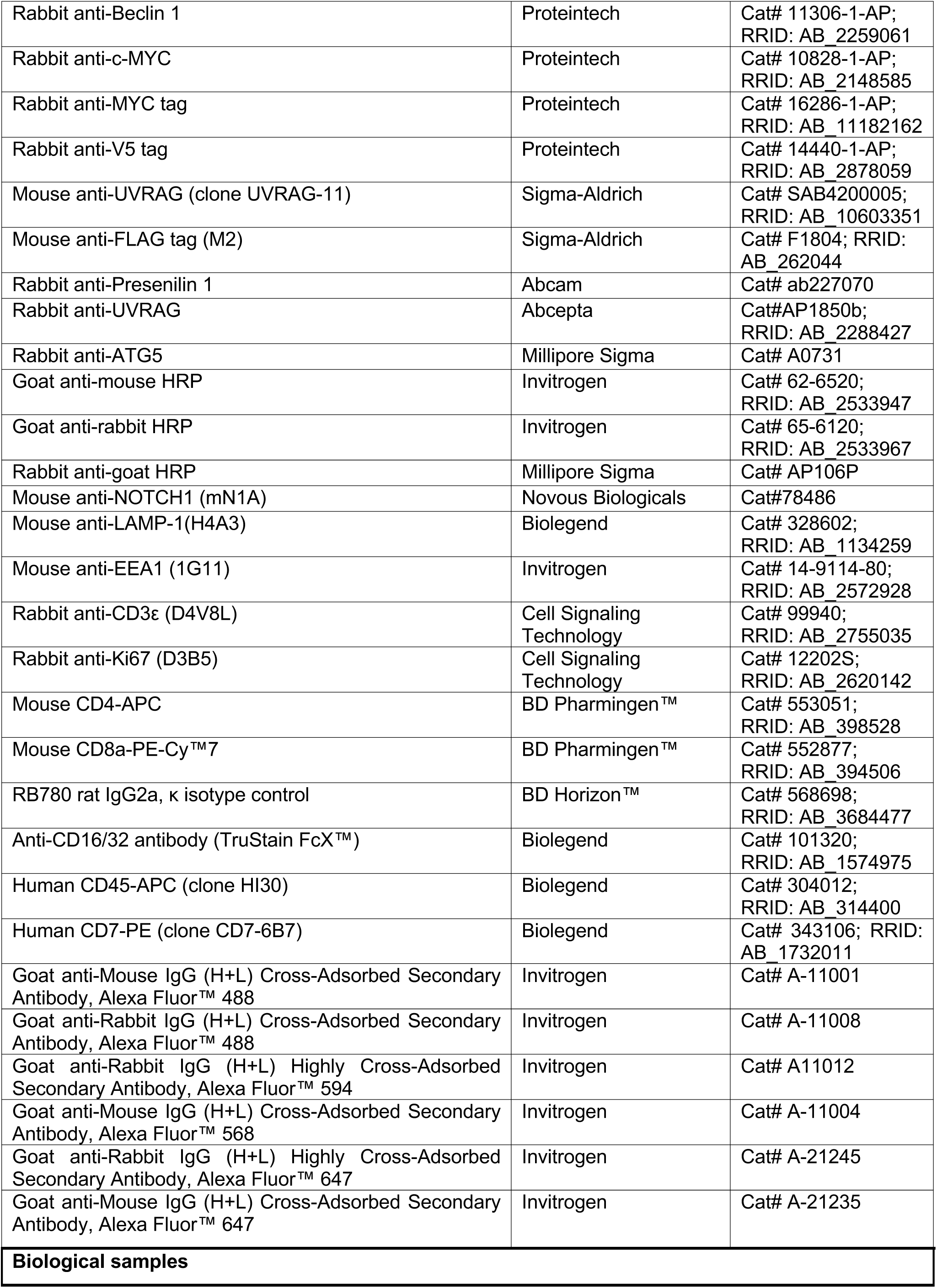
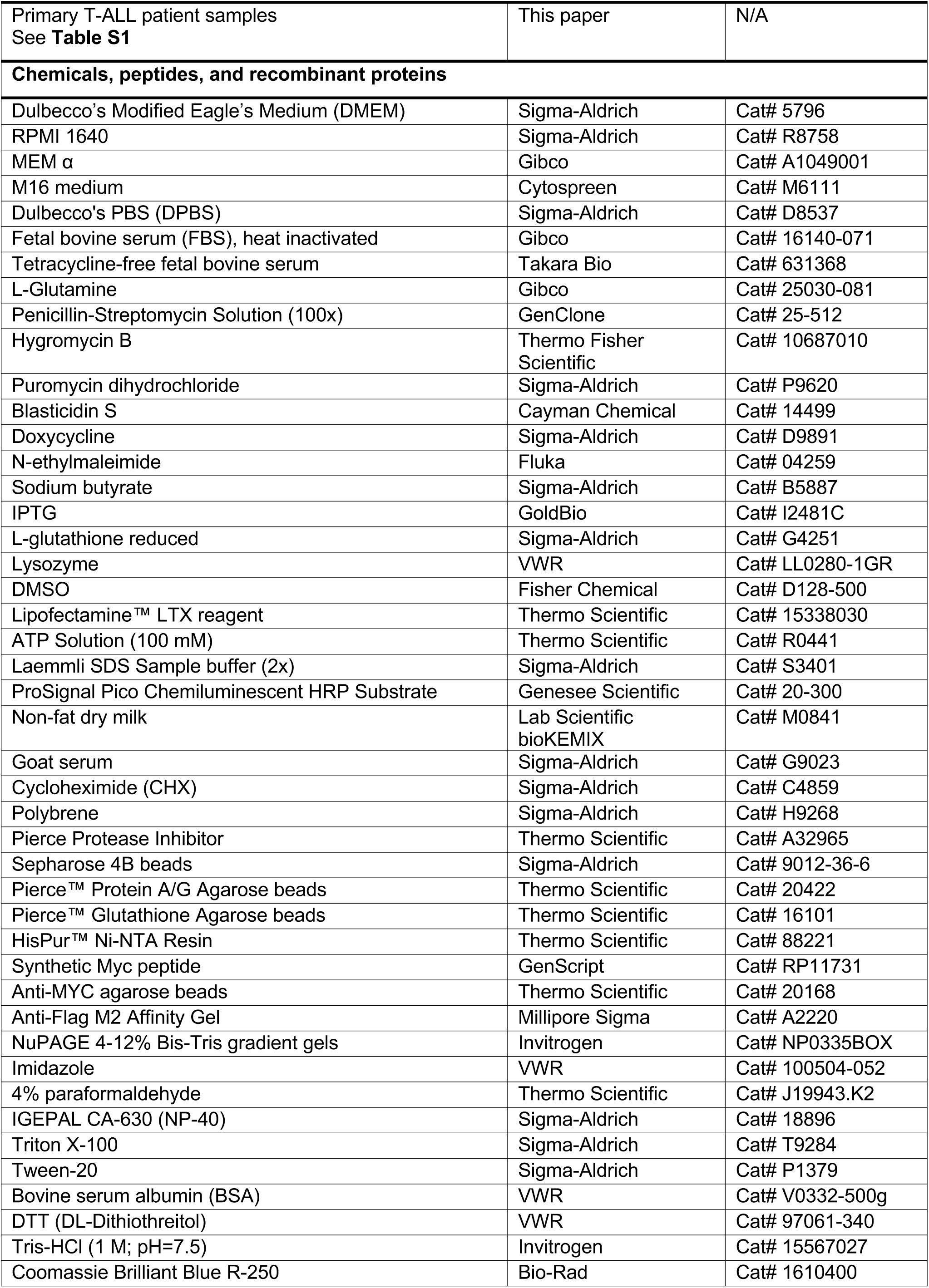
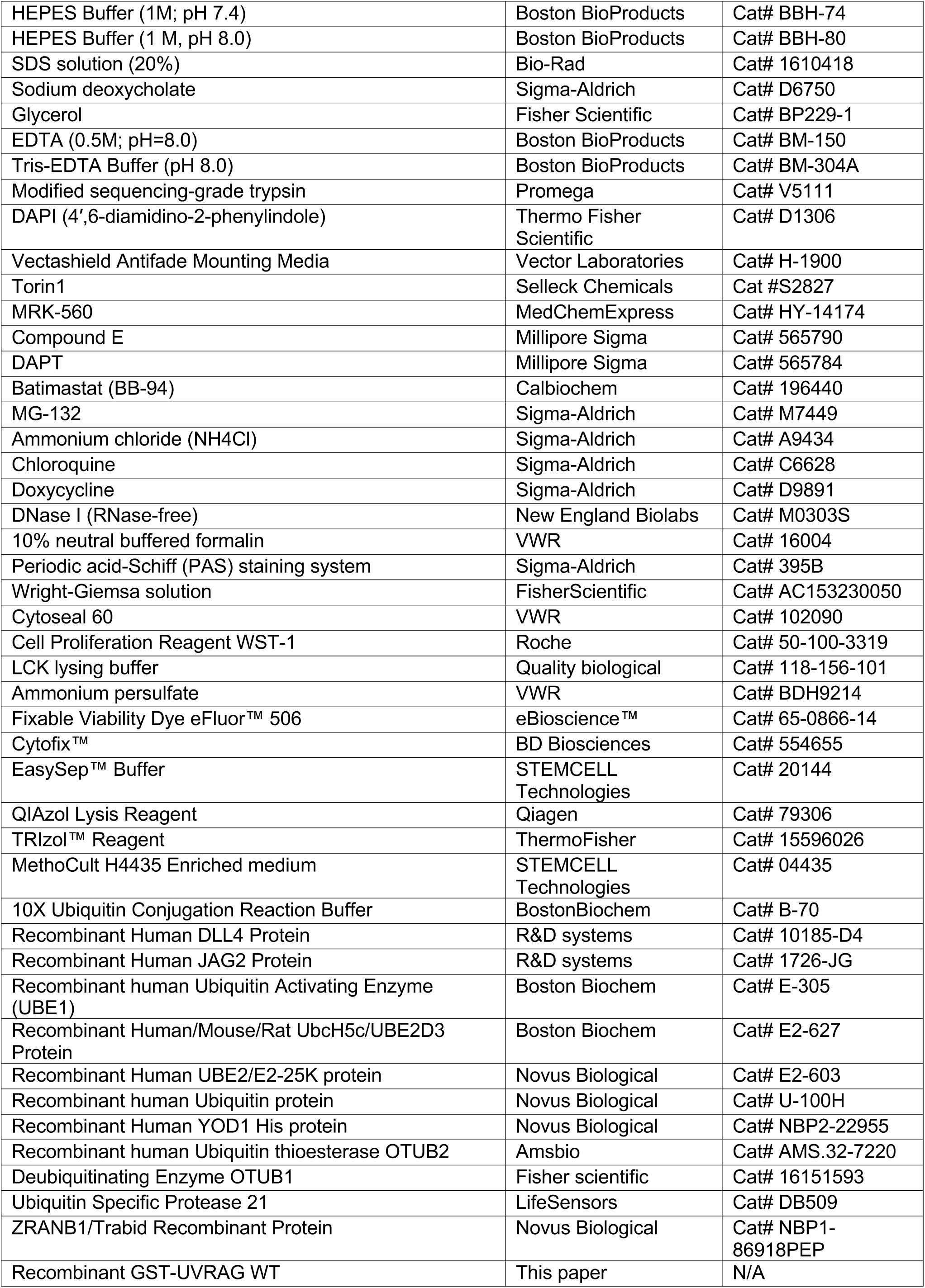
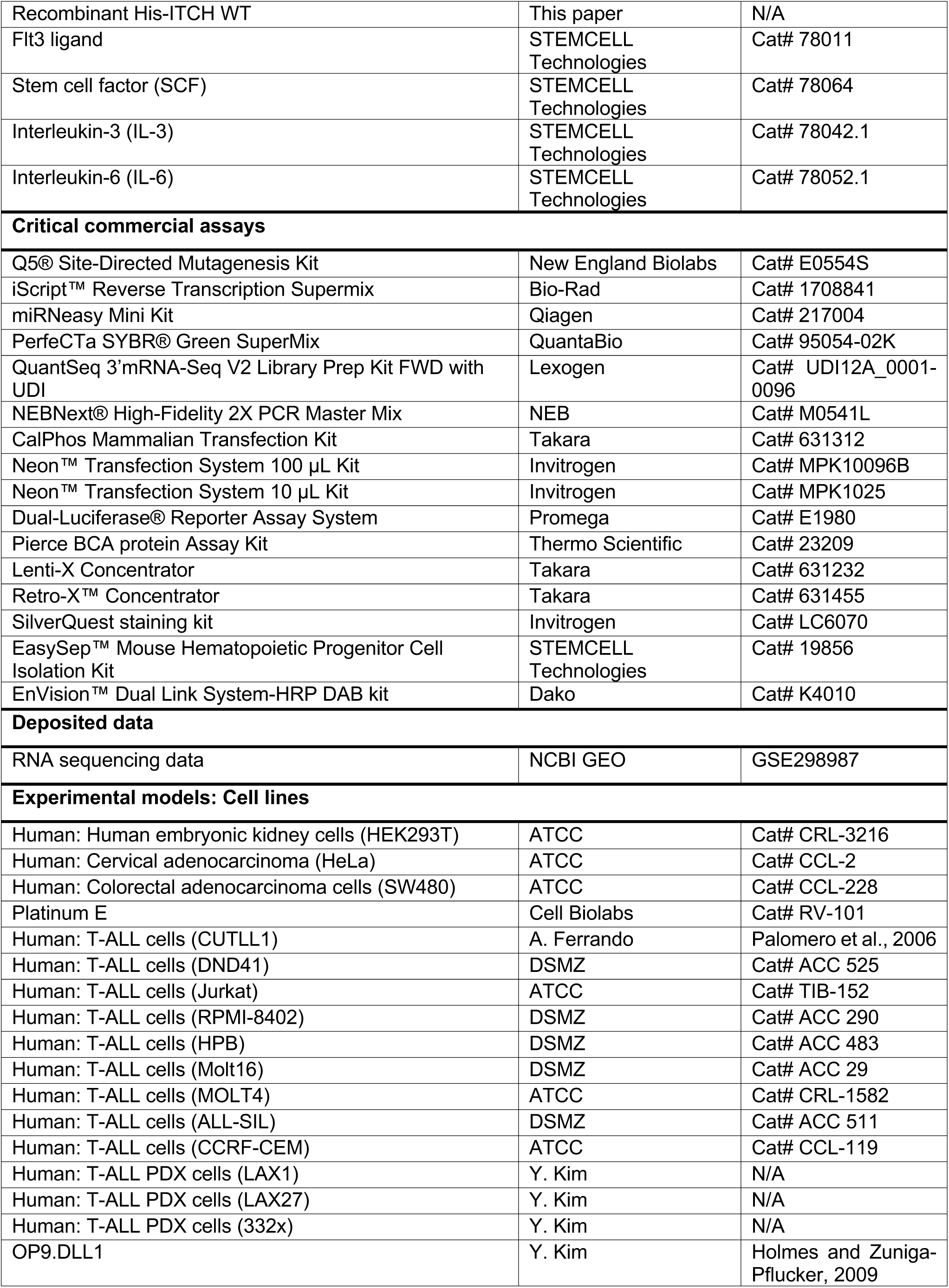
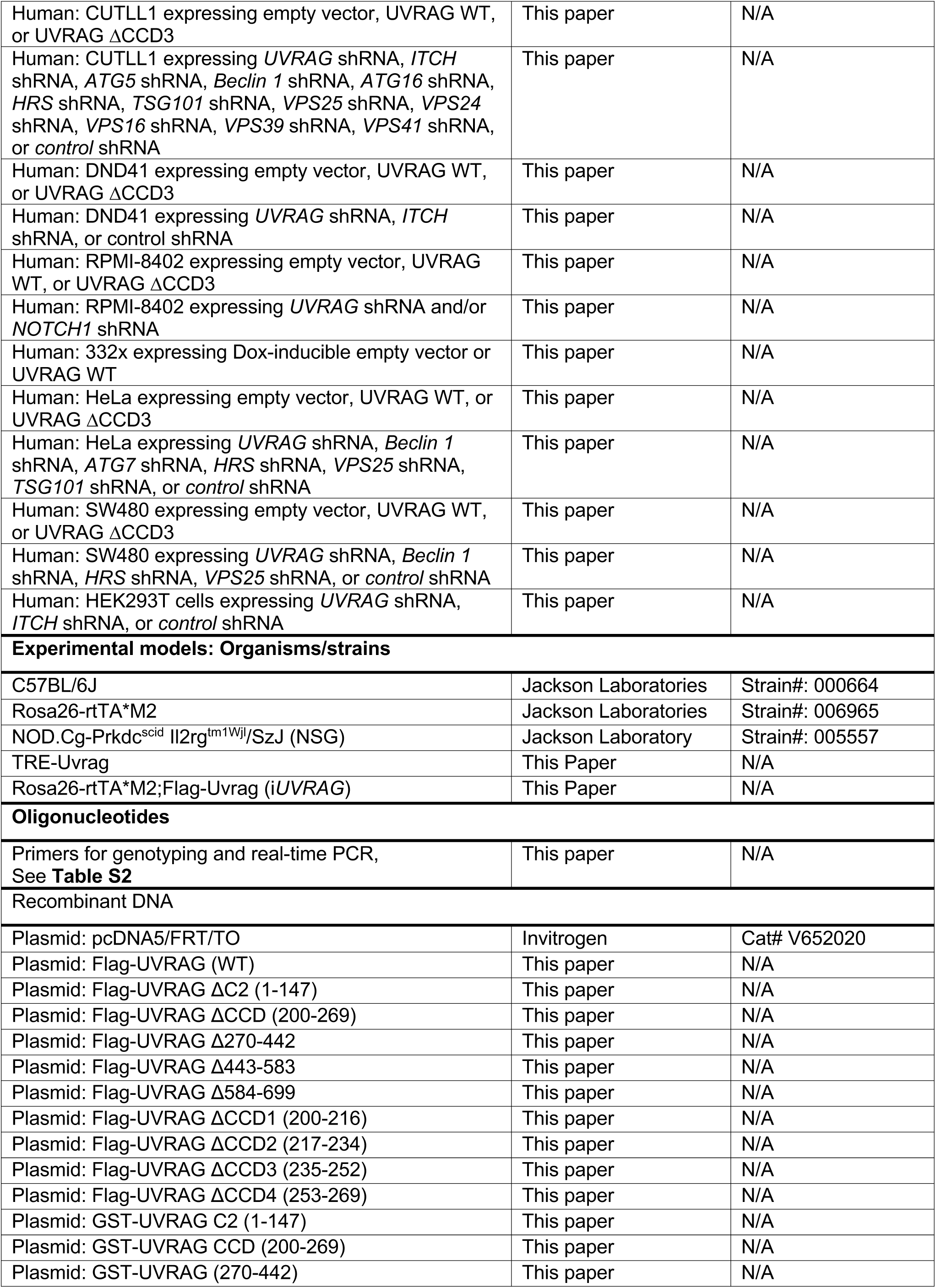
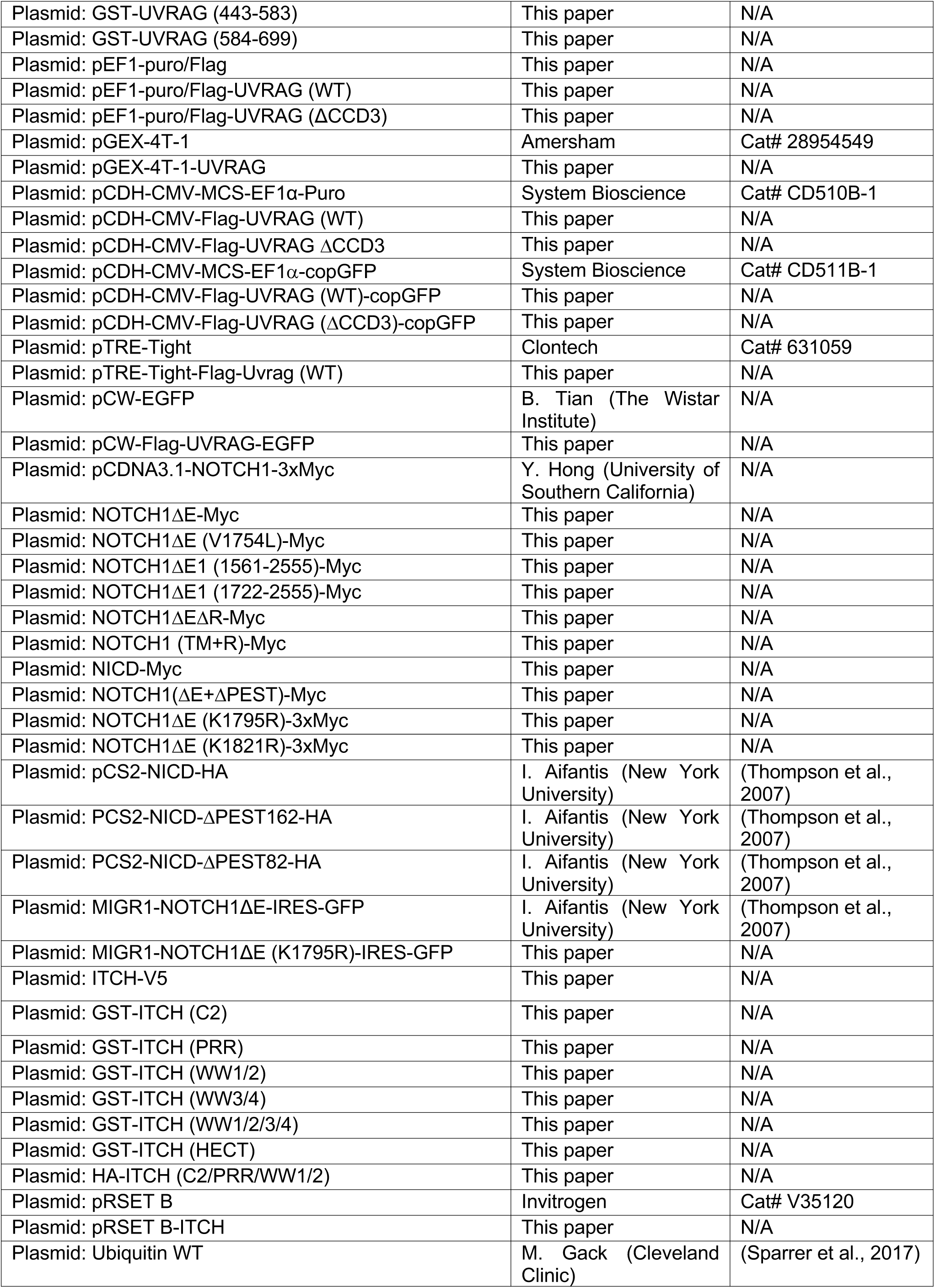
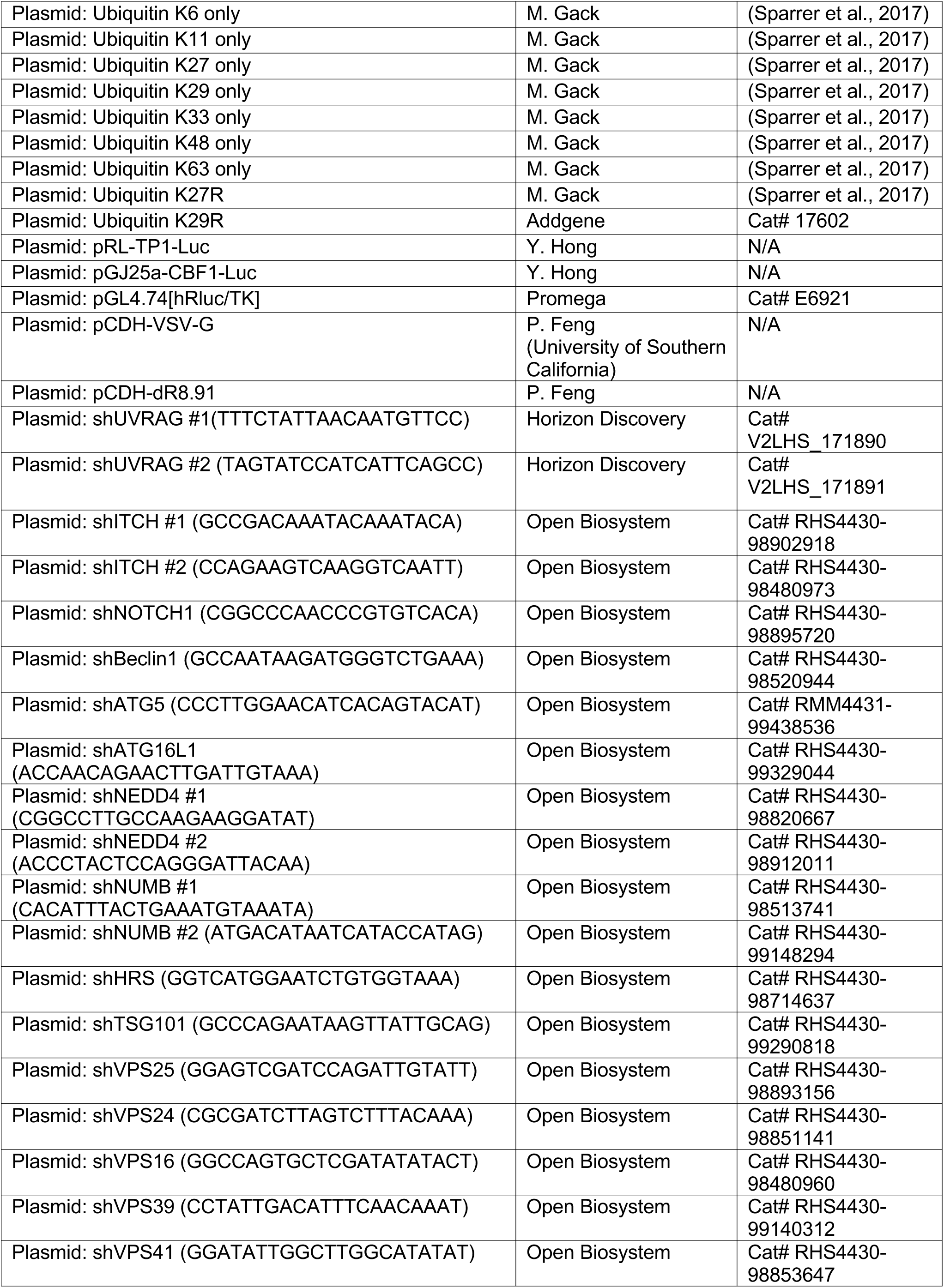
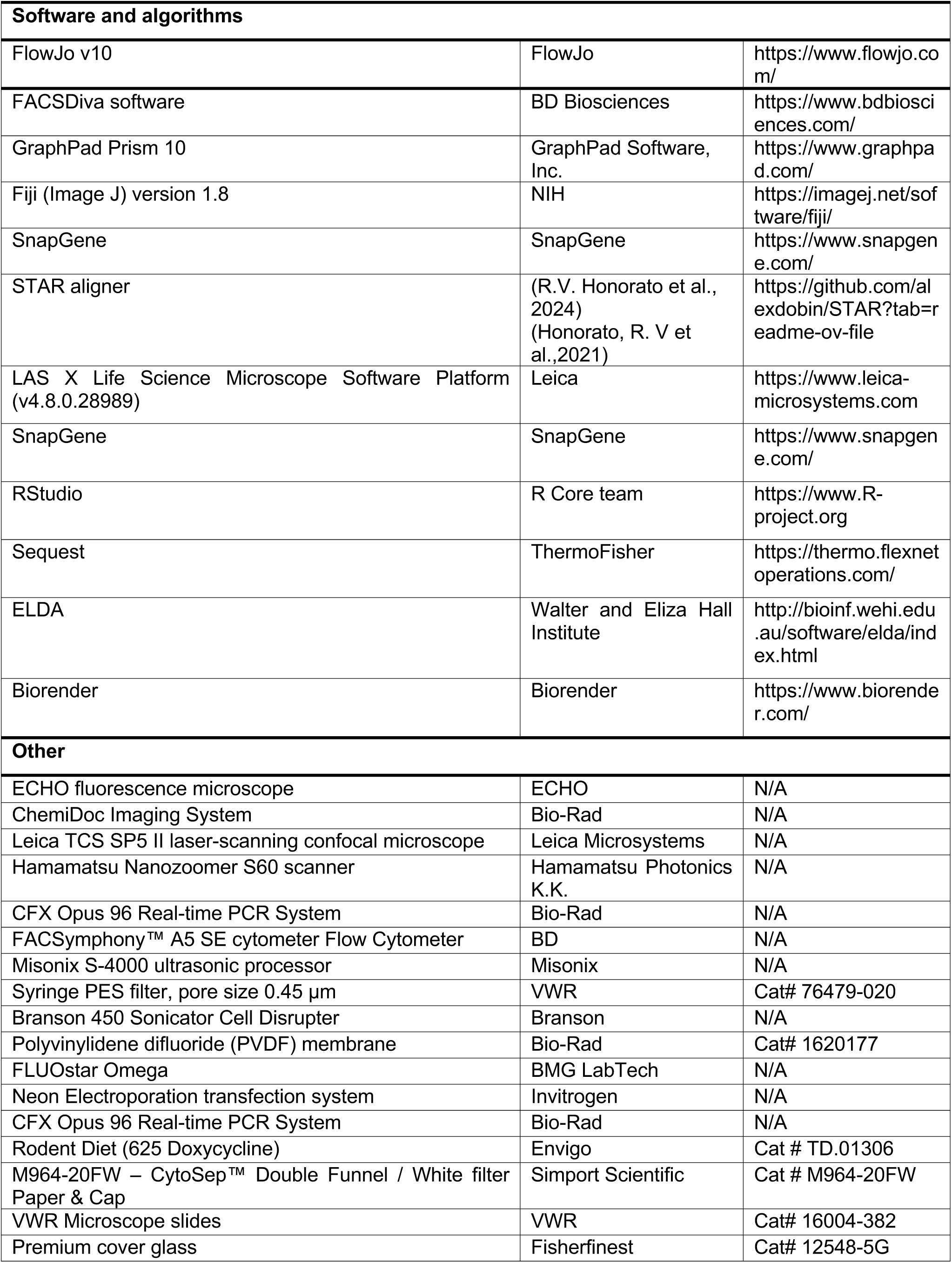

### RESOURCE AVAILABILITY

#### Lead contact

Further information and requests for resources and reagents should be directed to and will be fulfilled by the lead contact, Chengyu Liang.

#### Materials availability

Constructs and reagents in this study will be made available upon request, but a completed Materials Transfer Agreement may be required if there is potential for commercial application.

#### Data and code availability

- This paper does not report original code.
- The raw RNA sequencing data generated for this study have been deposited at NCBI GEO database under accession number GSE298987. The results shown here are in whole or part based upon data generated by the TCGA Research Network: http://cancergenome.nih.gov/. Previously published datasets that were reanalyzed during the study include microarray data from patients with T-ALL (GSE13159), the database of genotypes and phenotypes (dbGaP: http://www.ncbi.nlm.nih.gov/gap) under accession number phs000218 (TARGET) and sub-study specific accession phs000464 (TARGET ALL Expansion Phase 2) and phs002276. Any additional information are available from the corresponding author upon request. Source data will be provided with this paper.

### EXPERIMENTAL MODEL AND SUBJECT DETAILS

#### Cells culture and primary cell samples

Human embryonic kidney cells (HEK293T), HeLa cells, human colorectal adenocarcinoma cells (SW480), and Platinum E (Plat-E) cells for retroviral packaging were cultured in Dulbecco’s Modified Eagle’s Medium (DMEM) supplemented with 10% (v/v) fetal bovine serum (FBS), 2 mM L-glutamine (Gibco), and 100 U ml^-1^ penicillin-streptomycin (Pen-Strep). The human T-ALL cell lines MOLT16, ALL-SIL, RPMI-8402, HPB, DND41, Jurkat, CCRF-CEM, and MOLT4, and CUTLL1 were cultured in RPMI 1640 medium supplemented with 15% (v/v) FBS, 2 mM L-glutamine, and 100 U ml^-1^ Pen-Strep. The primary relapsed T-ALL samples 332x, LAX1, and LAX27 were provided by Y. Kim; The primary relapsed T-ALL samples 332x cells were maintained in Eagle’s Minimum Essential Medium (EMEM); LAX1 and LAX27 were maintained on the bone marrow-derived stromal cell line OP9 expressing the NOTCH ligand DLL1(Holmes and Zuniga-Pflucker, 2009) in IMDM 1640 (Gibco). Both media were supplemented with 20% (v/v) FBS, 2 mM L-glutamine, and 100 U ml^-1^ penicillin-streptomycin. Anonymized de-identified thymic tissue from children undergoing cardiac surgery was collected at Children’s Hospital Los Angeles (CHLA) through a CHLA institutional review board approved protocol. Primary T-ALL patient samples (**Table S1**) were collected at CHLA following informed consent under a CHLA Institutional Review Board (IRB) approved protocol. All cells were maintained at 37°C in a humidified atmosphere with 5% CO₂. Commercially obtained cells (ATCC) were authenticated by the vendor and were not further validated by our laboratory. Cells obtained from other laboratories, where these lines were previously published, were also not re-authenticated. All cell lines were routinely tested for mycoplasma contamination by PCR. Where indicated, cells were treated with 20 mM chloroquine (CQ; Sigma-Aldrich), 10 mM NH4Cl (Sigma-Aldrich), 10 mM MG132 (Sigma-Aldrich), 1 mg ml^-1^ doxycycline (Sigma-Aldrich), 1 mM Torin 1 (Selleckchem), 30 mM Batimastat (Calbiochem), 10 mM DAPT (Millipore Sigma), 1 mM compound E (Millipore Sigma), and MRK-560 (MedChemExpress) for the indicated time points before collection.

#### Animal Studies

##### *iUvrag* mice model

For generation of *iUvrag* mice, the *pTRE-Tight-Flag-Uvrag* plasmid was digested with *Xho*I to release the transgenic cassette. The gel-purified cassette was microinjected into the pronuclei of fertilized one-cell stage embryos (B6D2F1 background) with standard procedures. Injected embryos were cultured overnight in M16 medium (Cytospreen) at 37°C in 5% CO₂. All the two-cell stage embryos were then transferred into oviducts of the pseudopregnant CD-1 female mice at 0.5 dpc by Norris Comprehensive Cancer Center Transgenic Mice Core Facility (USC). Integration of the construct was confirmed by PCR (**Table S2**). Two independent founder lines were identified and back-crossed for more than 20 generations to C57BL/6 mice (Jackson Laboratories). *TRE-Uvrag* transgenic mice were crossed with *Rosa26-rtTA*M2* mice (Jackson Laboratories) in a pure C57BL/6 background to generate the double-transgenic mice (*Rosa26-rtTA*M2;Flag-Uvrag),* denoted as *iUVRAG*. Animals were maintained on the C57BL/6 background. To turn on the expression of *UVRAG*, *iUvrag* mice were administered a doxycycline (Dox) diet (Envigo). Both male and female control and *iUvrag* mice were used in these experiments.

##### Bone marrow (BM) transduction and transplantation

Bone marrow transplantation to generate the mouse model for NOTCH1ΔE-GFP was performed as previously described (King et al., 2013). Briefly, BM from Dox-treated *iUvrag* mice was enriched for c-Kit⁺ hematopoietic stem and progenitor cells (HSPCs) by the magnetic selection using the EasySep™ Mouse Hematopoietic Progenitor Cell Isolation Kit (STEMCELL Technologies). Isolated cells were cultured in RPMI with 20% FBS supplemented with 50 ng ml^-1^ Flt3 ligand, 50 ng ml^-1^ stem cell factor (SCF), 10 ng ml^-1^ interleukin-3 (IL-3), and 10 ng ml^-1^ interleukin-6 (IL-6), all from STEMCELL Technologies. On days 1 and 2 post-isolation, BM progenitor cells were transduced by spin inoculation (1,500 x *g*, 30°C, 90 min) with concentrated retroviral supernatant (*NOTCH1ΔE-IRES-GFP*) in the presence of 8 μg ml^-1^ polybrene. Transduction efficiency was assessed by GFP fluorescence at 96 h. 5×10^5^ GFP⁺ cells were transferred *via* retro-orbital injection into irradiated (5.5 Gy × 2, total 11 Gy) congenic 6-8 weeks old recipient mice (C57BL/6) along with 2×10⁵ unfractionated BM mononuclear cells for hematopoietic support (King et al., 2013). To induce UVRAG expression in NOTCH1-induced leukemia, mice were treated with Dox diet (Envigo) after engraftment. Mice were monitored daily after transfer over the course of the disease and analyzed for leukemia as described. Survival analysis was performed using the Mantel-Cox test.

For the secondary transplants, GFP⁺ leukemic cells transduced with NOTCH1ΔE were isolated from the spleen of primary recipient mice using fluorescence-activated cell sorting (FACS Symphony S6 SE). A total of 5×10^5^ purified GFP⁺ leukemic cells were transplanted into sub-lethally irradiated (4.5 Gy) C57BL/6 recipient mice (8-10 weeks old) *via* retroorbital injection. Tertiary transplantation was performed by repeating the same procedure using splenic GFP⁺ leukemic cells isolated from secondary recipients. Mice were monitored regularly for weight loss, morbidity, and overall survival.

##### *In vivo* limiting dilution assay

GFP⁺ leukemic cells transduced with NOTCH1ΔE were isolated from the spleen of primary recipient mice treated with or without Dox. Cells were enriched by fluorescence-activated cell sorting (FACS Symphony S6 SE) and serially diluted to final doses of 1×10⁴, 5×10⁴, 2×10^5^, to 5×10^5^ GFP⁺ cells per mouse. Each dilution was injected into groups of sub-lethally irradiated (4.5 Gy) 8-10-week-old C57BL/6 recipient mice *via* retro-orbital injection. To support hematopoietic recovery and engraftment, each recipient also received 2×10⁵ freshly isolated WT bone marrow cells. Mice were monitored for signs of leukemia development over a four-week period following transplantation. The frequency of LICs and the 95%-confidence interval (CI) were calculated using extreme limiting dilution (ELDA) software (http://bioinf.wehi.edu.au/software/elda/index.html), which applies a single-hit Poisson model to estimate stem cell frequency and statistical significance across experimental conditions.

##### T-ALL xenograft studies

For CUTLL1 xenograft study, 6-8-week-old *NOD.Cg-Prkdc^scid^ Il2rg^tm1Wjl^/SzJ* (NSG) mice (The Jackson Laboratory) were sub-lethally irradiated (2.5 Gy) and injected retro-orbitally with 5 × 10⁶ CUTLL1 cells stably expressing GFP and/or the indicated UVRAG constructs. Mice were monitored regularly for signs of disease and peripheral blood was collected at defined intervals for flow cytometric analysis of hCD45^+^hCD7^+^ T-ALL cells. Mice were euthanized at defined time points or upon signs of morbidity, and tissues including bone marrow, spleen, and liver were collected for flow cytometry, immunohistochemistry, and histopathology. Spleen weights were recorded at necropsy. Survival studies were analyzed using Kaplan–Meier methodology.

For drug treatment studies, xenografted mice were monitored by peripheral blood analysis for hCD45^+^hCD7^+^ T-ALL cell engraftment. When peripheral blood leukemia burden reached approximately 10%, mice were randomized to receive DAPT (γ-secretase inhibitor, 10 mg kg^-1^ per day, *i.p.*), Torin 1 (mTOR inhibitor 20 mg kg^-1^ per day, *i.p.*), or Vehicle control for 21 consecutive days. Mice were euthanized upon signs of morbidity or at pre-defined endpoints.

To measure leukemia response to MRK-560 therapy, mice with CUTLL1 xenografts or patient-derived xenografts (PDX 332x) harboring a Dox-inducible UVRAG construct were randomized after hCD45^+^hCD7^+^ T-ALL cells reached approximately 10% in peripheral blood to receive MRK-560 (3 or 30 mmol kg^-1^, *i.p.* daily) or Vehicle control for 14 days. In the 332x PDX model, mice were concurrently placed on a Dox diet to induce UVRAG expression, starting at the same time as MRK-560 treatment and continuing until experimental endpoint. Peripheral blood, spleen, bone marrow, and intestine were collected at experimental endpoints for flow cytometric analysis of hCD45^+^hCD7^+^ T-ALL cells or histopathology, and spleen weight measurements. Survival was assessed using Kaplan–Meier curves and analyzed by the log-rank (Mantel–Cox) test.

All mice were maintained in a pathogen-free facility with *ad libitum* access to food and water. All animal procedures were performed in accordance with institutional guidelines and with approval from the Institutional Animal Care and Use Committee (IACUC).

### METHOD DETAILS

#### Plasmids

Wild-type (WT) *UVRAG* and *UVRAG* mutants were constructed by cloning the cDNA of the full-length or truncated mutants into the *Kpn*I/*Not*I sites of the pcDNA5/FRT/TO vector (Invitrogen, USA) with an N-terminal GST tag or Flag tag; or into the *Afl*II/*Not*I sites of the pEF/puro-Flag vector; or into the *Nhe*I/*Not*I Sites of the pCDH-CMV-MCS-EF1a-Puro and pCDH-CMV-MCS-EF1a-copGFP vectors; or into the *BamH*I/*Not*I sites of pGEX-4T-1 with a GST tag; or into the *Nhe*I/*Sal* I sites of pCW-EGFP vector (provided by B. Tian, The Wistar Institute); or into the *Mlu*I/*Not*I sites of the pTRE-Tight vector. NOTCH1ΔE WT and mutants in **Figure S3F** were constructed by subcloning the cDNA of the full-length or mutant NOTCH1ΔE amplified from the pcDNA3.1-NOTCH1-3xMyc vector into the *BamH*I/*Xho*I sites of the pcDNA3.1 vector with a C-terminal 3xMyc tag. Site-directed mutagenesis of NOTCH1ΔE was performed using Q5^®^ Site-Directed Mutagenesis Kit, according to manufacturer instruction. WT *ITCH* and *ITCH* mutants were constructed by cloning the cDNA of the full-length or truncated mutants into the *Kpn*I/*Not*I sites of the pcDNA5/FRT/TO vector with an N-terminal HA tag or GST tag or C-terminal V5 tag; or into pRSET B between *BamH*I and *EcoR*I sites. All constructs were verified by Sanger sequencing (ABI PRISM 377).

#### Transfection and RNA interference

Plasmids were transiently transfected using the Calcium Phosphate Transfection Kit (Takara) or Lipofectamine™ LTX reagent (Thermo Scientific) for HEK293T cells, according to manufacturer protocols. For the transfection of T-ALL cells, 10 μg plasmid DNA was electroporated into 2×10^6^ cells in 100 ml Neon^TM^ transfection reagent (Invitrogen, MPK1025 and MPK10096) using the Neon^TM^ Electroporation transfection system (Invitrogen; 1,400 V, 10 ms pulse width, 3 pulses), followed by 48 h of culture. For retrovirus production, Plat-E packaging cells were transfected with MIGR1-NOTCH1ΔE (WT, K1795R)-GFP plasmid using the Calcium Phosphate Transfection method. For lentivirus production, HEK293T cells were transfected with the lentiviral expression or shRNA plasmids along with packaging constructs pCDH-VSV-G and pCDH-dR8.91, using the Calcium Phosphate Transfection method. The virus-containing medium was collected 60 h after transfection, filtered through 0.45-μm PES filter (VWR), concentrated with Retro-X™ Concentrator (Takara Bio) for retrovirus or Lenti-X Concentrator (Takara Bio) for lentivirus, and supplemented, with 8 mg ml^-1^ polybrene (Sigma-Aldrich). Mouse c-Kit^+^ hematopoietic stem cells (HSCs), isolated with EasySep™ Mouse Hematopoietic Progenitor Cell Isolation Kit (STEMCELL Technologies), were infected by replacing their culture medium with the retrovirus-containing medium for 8 h after centrifugation at 1,500 x *g* for 90 minutes. Other target cells were infected by replacing their culture medium with the lentivirus-containing medium for 12 h. Selection of transduced cells was carried out using either 1 mg ml^-1^ puromycin (Sigma-Aldrich), 5 mg ml^-1^ blasticidin S (Cayman Chemical), depending on the vector selection marker. Gene expression and knockdown efficiency were determined by immunoblot using gene-specific antibodies.

#### Ligand stimulation of cells

Lyophilized recombinant human DLL4 and JAG2 were purchased from R&D Systems and reconstituted at 100 mg mL^-1^ in PBS containing 0.1% bovine serum albumin. For stimulation of cultured HeLa or SW480 cells, culture dishes were coated with 5 mg mL^-1^ DLL4 (for HeLa cells) or 5 mg mL^-1^ JAG2 (for SW480) in PBS and incubated overnight at 4°C. Cells were seeded onto the coated dishes the following day for subsequent NOTCH1 stimulation assays.

#### Cell viability analysis

T-ALL cells were seeded at 1 × 10⁴ cells per well in 96-well plates. After overnight incubation, cells were treated with MRK-560 (MedChemExpress) at the indicated concentrations for 7 days. Viability was assessed using the WST-1 reagent (Roche) following the manufacturer’s protocol. Briefly, the medium was aspirated, and 10 μL of WST-1 diluted in complete medium was added to each well, followed by incubation for 0.5–4 h at 37°C with a brief 1-minute (min) agitation to ensure proper mixing. Absorbance was then measured using a FLUOStar Optima microplate reader.

#### *In vitro* colony-forming assays

Colony forming assays were performed as previously described (King et al., 2013) with minor modifications. Briefly, 1,000 CUTLL1 cells were suspended in 1 mL of methylcellulose-based medium (MethoCult H4435; STEMCELL Technologies) and plated in triplicate into 6-well plates. Colonies were counted on day 10, harvested, and replated at 1,000 cells per well for subsequent passages. This serial replating was performed for four rounds (P1-P4), with colony numbers recorded at the end of each passage. Colony images were acquired using an Echo Revolve microscope (Echo Laboratories). Colony-forming units (CFUs) were quantified from triplicate wells at each passage.

#### Luciferase reporter assay

Luciferase reporter assays were performed with the Dual-Luciferase Reporter Assay System (Promega E1980) and read on a FLUOStar Optima microplate reader (BMG LabTech), according to the manufacturer’s instructions. Briefly, 48 h after co-transfection with NOTCH-responsive firefly luciferase reporters, the constitutive Renilla luciferase control plasmid pGL4.74[hRluc/TK] (Promega], and the indicated plasmids, cells were lysed and luciferase activity was measured. For shRNA experiments, cells were first transduced with lentiviral shRNAs and, 24-36 h later, transfected with the NOTCH luciferase reporters and the Renilla control plasmid. For experiments in which NOTCH activity was induced by DLL4 or JAG2, transfected cells were replated onto DLL4- or JAG2-coated dishes 6 h post-transfection, and luciferase activity was measured 24 h later. Firefly luciferase activity was normalized to Renilla luciferase to control transfection efficiency, and results are presented relative to the control.

#### RT-qPCR analysis

Total RNA from T-ALL samples was extracted using QIAzol Lysis Reagent and purified using the miRNeasy Mini Kit, following the manufacturer’s instructions. RNA from other cells was extracted using Trizol reagent (Invitrogen), according to the manufacturer’s instructions. RNA quality and quantity were examined using a NanoDrop Lite spectrophotometer. cDNA synthesis was performed using the iScript™ cDNA synthesis kit (Bio-Rad), and 50 ng cDNA was used for qPCRs with PerfeCTa SYBR Green Master SuperMix (Quantabio) on a CFX Opus 96 Real-time PCR System (Bio-Rad). Relative mRNA expression was calculated using the comparative threshold (ΔΔC_T_) method with *GAPDH* as the internal control. All reactions were performed in at least triplicates. See **Table S2** for primer sequence details.

#### Immunoprecipitation and immunoblotting

Immunoprecipitation of endogenous proteins or overexpressed epitope-tagged proteins was performed as previously described (Yang et al., 2016), Briefly, transfected cells were lysed in lysis buffer (25 mM Tris-HCl pH 7.5, 300 mM NaCl, 1 mM EDTA, and 1-2% (v/v) IGEPAL CA-630 (NP-40) (Sigma-Aldrich) or Triton X-100 (Sigma-Aldrich), and protease inhibitor cocktail (Thermo Scientific^TM^), or in RIPA buffer [(50 mM Tris-HCl pH 8.0, 150 mM NaCl, 1 mM EDTA, 2% Triton X-100, 0.5% sodium deoxycholate, 1mM DTT, 0.1% SDS, protease inhibitor cocktail, and 10 mM N-ethylmaleimide] for ubiquitination detection of NOTCH1. Lysates underwent freeze-thaw cycles and sonication (15% amplitude, 10 s process time, 5 s push-on, and 1 s push-off). Cell debris was pelleted by centrifugation (14,500 × *g*, 15 min, 4 °C). Protein concentrations were determined using the Pierce BCA Protein Assay Kit (Thermo Scientific^TM^). An aliquot of the cleared lysate was reserved as whole cell lysate (WCL) control. After preclearing with Sepharose 4B beads (Sigma-Aldrich) for 2 h at 4 °C with gentle rotation, cell extracts were subjected to immunoprecipitation using primary antibodies (1-2 mg ml^-1^) as indicated overnight (12-16 h) at 4°C. The immune complexes were captured using protein A/G agarose beads (Thermo Scientific^TM^) for 2 h at 4°C. Beads were washed extensively with lysis buffer, and immunoprecipitates were eluted by heating in Laemmli Sample Buffer (Sigma-Aldrich) at 95°C for 5 min. Proteins were resolved by SDS-PAGE, transferred to PVDF membranes (Bio-Rad), blocked with 5% non-fat milk (Lab Scientific bioKEMIX) in PBST, and probed with the indicated antibodies. Horseradish peroxidase (HRP)-conjugated secondary antibodies (1:3000) were used for detection with ProSignal Pico Chemiluminescent HRP Substrate (Genesee Scientific), and signals were visualized using a ChemiDoc Imaging System (Bio-Rad). Immunoblot images were acquired with a linear or sigmoid gradation conversion curve.

#### Protein purification and GST pull-down assay

Recombinant proteins were expressed in BL21(DE3) cells transformed with pGEX-4T-1-UVRAG (for GST-UVRAG) or pRSET B-ITCH (for His-ITCH). Protein expressions were induced with 1 mM isopropyl *b*-D-1-thiogalactopyranoside (IPTG) at 20°C overnight. Cells were harvested, subjected to three freeze-thaw cycles, and re-suspended in pre-chilled lysis buffer (125 mM Tris, pH 9.0, 300 mM NaCl, 10% glycerol, 0.05% Triton X-100, 1x protease inhibitors, 1mg mL^-1^ lysozyme, and 1µg mL^-1^ DNase I). Lysates were sonicated (3 x 30 s on ice at 60% amplitude with 15 min intervals) and centrifuged to remove debris. The clarified supernatant was incubated with equilibrated glutathione agarose beads (for GST-UVRAG) or HisPur Ni-NTA resin (for His-ITCH) for 1 h at 4 °C with rotation. After three washes with ice-cold PBST (PBS with 1% Triton X-100), GST-UVRAG was eluted using 10 mM reduced L-glutathione in 50 mM Tris-HCl, 150 mM NaCl, pH 8.0, and His-ITCH was eluted with 100 mM imidazole (VWR) in 20 mM NaH_2_PO4, 300 mM NaCl, pH 8.0, followed by desalting to remove imidazole. Protein concentrations were determined using the Pierce BCA Protein Assay kit (Thermo Scientific^TM^).

For purification of NOTCH1ΔE+ΔPEST protein, HEK293T cells were transiently transfected with pcDNA3.1-NOTCH1ΔE+ΔPEST-3xmyc using the Calcium Phosphate method. 48 h post-transfection, cells were harvested by centrifugation at 500 x *g* for 5 min at 4 °C and re-suspended in ice-cold lysis buffer (25 mM Tris pH 7.5, 150 mM NaCl, 1 mM EDTA, and 1% NP-40) supplemented with a protease inhibitor cocktail (Thermo Scientific^TM^). Cells were lysed by sonication on ice (15% amplitude, 10 s processing time, 5 s push-on, 1 s push-off). The clarified supernatant was incubated with anti-c-Myc agarose beads (Thermo Scientific^TM^) overnight at 4 °C with gentle rotation. Beads were washed 3 times with high-stringency RIPA buffer (50 mM Tris-HCl pH 7.4, 150 mM NaCl, 1 mM EDTA, 1% NP-40, 0.5% sodium deoxycholate, 0.1% SDS, and protease inhibitor cocktail) to remove nonspecific bound and interacting proteins. The bound NOTCH1ΔE+ΔPEST protein was eluted using synthetic Myc peptide (GenScript) according to the manufacturer’s protocol. Protein concentration was measured using the Pierce BCA Protein Assay Kit following the manufacturer’s instructions.

For the GST pull-down assay, GST-UVRAG bound to glutathione agarose beads was incubated with 5 μg purified His-ITCH in Triton X-100 buffer (150 mM NaCl, 5 mM EDTA, 1% (v/v) Triton X-100, 50 mM HEPES pH 7.4 plus protease inhibitor cocktail) at 4 °C for 4 h. Beads were washed extensively and bound proteins were eluted by heating in laemmli sample buffer at 95°C for 5 min, followed by immunoblotting.

#### *In Vitro* ubiquitination assay

GST-UVRAG and His-ITCH proteins were purified from bacteria; NOTCH1ΔE+ΔPEST was purified from HEK293T cell extract following the protocol described in “protein purification”. *In vitro* ubiquitination of NOTCH1ΔE+ΔPEST by ITCH was carried out in a reaction buffer containing 50 mM Tris pH 7.5, 5 mM MgCl2, 1 mM DTT, 1 mM ATP, 100 nM UBE1 (E1 enzyme, Boston Biochem), 500 nM UbcH5c (E2 enzyme, Boston Biochem), 1 μg purified ITCH E3 ligase, 1 μg purified NOTCH1ΔE+ΔPEST protein substrate bearing a C-terminal 3xMyc tag, with or without 1 μg purified GST-UVRAG. Reactions were initiated by addition of ubiquitin (Boston Biochem, 5-10 μg final) and incubated at 30°C for 1 h in a total volume of 30 μl. The reaction was stopped by addition of Laemmli SDS sample buffer and heating at 95°C for 5 min. Samples were resolved by SDS-PAGE and analyzed by immunoblotting using anti-Myc antibody.

#### *In vitro* DUB restriction assay

DUB restriction assay was performed as previously described (Sparrer et al., 2017). Briefly, HEK293T cells overexpressing NOTCH1ΔE-Myc, FLAG-UVRAG, and HA-Ub WT were lysed in NP-40 buffer (150 mM NaCl, 1% (v/v) NP-40, 50 mM HEPES pH 7.4, and protease inhibitor cocktail) at 60 h post-transfection. Cell debris was pelleted by centrifugation (14,500 × *g*, 15 min, 4 °C). Post-centrifuged WCLs were incubated with anti-c-Myc antibody overnight at 4°C, and then captured using protein A/G agarose beads (Thermo Scientific^TM^) for 2 h at 4°C. Myc-precipitates were rigorously washed with 500 mM NaCl-containing NP40 lysis buffer and then incubated in DUB reaction buffer (25 mM Tris, pH 7.5, 150 mM NaCl, and 4 mM dithiothreitol (DTT) together with 2μg of recombinant DUB enzymes including USP 21 (LifeSensors), OTUB1 (Fisher scientific), OTUB2 (Amsbio), TRABID (Novus Biological), or YOD1 (Novus Biological) at 30°C for 2 h. The reaction was terminated by adding Laemmli SDS sample buffer and heating at 95 °C for 5 minutes. Samples were resolved by SDS-PAGE and polyubiquitination of NOTCH1ΔE was determined by IB with anti-HA antibody.

#### Mass spectrometry

To identify UVRAG-binding partners in T-ALL cells, twenty 10 cm dishes of CUTLL1 cells expressing Vec or Flag-UVRAG were lysed with lysis buffer (25 mM Tris-HCl, pH 7.5, 300 mM NaCl, 1 mM EDTA, 1% Triton X-100 [v/v]) supplemented with a complete protease inhibitor cocktail, followed by centrifuge at 14,500 × *g* for 15 min at 4 °C. Post-centrifuged lysates were pre-cleared using Sepharose 4B beads (Sigma-Aldrich) to remove cell debris. Lysates were mixed with a ~50% slurry of anti-Flag M2 Affinity Gel (Millipore Sigma) and incubated for 8 h at 4°C. After extensive washing of the beads with lysis buffer, bound proteins were eluted in Laemmli SDS sample buffer for 5 min at 95°C and separated on a NuPAGE 4-12% Bis-Tris gradient gels (Invitrogen). To stain co-immunoprecipitated proteins, a SilverQuest staining kit (Invitrogen) was used according to the manufacturer’s instructions. Bands that were specifically present in the Flag-UVRAG sample, but not empty vector, were excised and analyzed by ion-trap mass spectrometry analysis at the Harvard Taplin Biological Mass Spectrometry facility (Boston). Amino acid sequences were determined by tandem mass spectrometry and database searches.

To identify the lysine residues in NOTCH1ΔE that are ubiquitinated by UVRAG/ITCH, twenty 15 cm dishes of HEK293T cells were each transfected with NOTCH1ΔE-Myc and Flag-UVRAG. At 48 h post-transfection, cells were lysed in lysis buffer (25 mM Tris-HCl, pH 7.5, 300 mM NaCl, 1 mM EDTA, 1% Triton X-100 [v/v]) supplemented with a complete protease inhibitor cocktail, followed by centrifuge at 14,500 × *g* for 15 min at 4 °C. Post-centrifuged lysates were pre-cleared twice using Sepharose 4B beads (Sigma-Aldrich) to remove cell debris. Clarified lysates were mixed with a ~50% slurry of anti-MYC agarose beads (Thermo Scientific), incubated for 8 h at 4 °C, and extensively washed with lysis buffer. Purified proteins were eluted in Laemmli SDS sample buffer for 5 min at 95°C and separated on a NuPAGE 4–12% Bis-Tris gradient gels (Invitrogen). The gel was stained with Coomassie Brilliant Blue (NuSep, Coomassie Electrophoresis Stain). Bands corresponding to unmodified and ubiquitinated forms of NOTCH1ΔE-Myc were excised and analyzed by ion trap mass spectrometry for post-translational modification at the Harvard Taplin Biological Mass Spectrometry facility (Boston).

Excised gel bands were cut into approximately 1 mm^3^ pieces. Gel pieces were then subjected to a modified in-gel trypsin digestion procedure (Shevchenko et al., 1996). Gel pieces were washed and dehydrated with acetonitrile for 10 min. followed by removal of acetonitrile. Pieces were then completely dried in a speed-vac. Rehydration of the gel pieces was with 50 mM ammonium bicarbonate solution containing 12.5 ng µl^-1^ modified sequencing-grade trypsin (Promega, Madison, WI) at 4°C. After 45 min., the excess trypsin solution was removed and replaced with 50 mM ammonium bicarbonate solution to just cover the gel pieces. Samples were then placed in a 37°C room overnight. Peptides were later extracted by removing the ammonium bicarbonate solution, followed by one wash with a solution containing 50% acetonitrile and 1% formic acid. The extracts were then dried in a speed-vac (~1 h). The samples were then stored at 4°C until analysis. On the day of analysis the samples were reconstituted in 5-10 µl of HPLC solvent A (2.5% acetonitrile, 0.1% formic acid). A nano-scale reverse-phase HPLC capillary column was created by packing 5 µm C18 spherical silica beads into a fused silica capillary (125 μm inner diameter x ~20 cm length) with a flame-drawn tip(Peng and Gygi, 2001). After equilibrating the column each sample was loaded via a Famos auto sampler (LC Packings, San Francisco CA) onto the column. A gradient was formed and peptides were eluted with increasing concentrations of solvent B (97.5% acetonitrile, 0.1% formic acid). As peptides eluted they were subjected to electrospray ionization and then entered into an LTQ Velos ion-trap mass spectrometer (ThermoFisher, San Jose, CA). Peptides were detected, isolated, and fragmented to produce a tandem mass spectrum of specific fragment ions for each peptide. Peptide sequences (and hence protein identity) were determined by matching protein databases with the acquired fragmentation pattern by the software program, Sequest (ThermoFisher, San Jose, CA)(Eng et al., 1994). Spectral matches were manually examined and multiple identified peptides per protein were required. The differential modifications of 114.0429 Daltons on lysine was used for identifying ubiquitinated residues. The data was filter to a 1% peptide false discovery rate.

#### Autophagy analysis

For assessment of autophagy, HEK293T cells expressing Vec, UVRAG (WT or mutant) were treated with 50 nM Torin 1 (Selleck Chemicals) for 3 h. Cells were lysed in ice-cold RIPA lysis buffer (50 mM Tris-HCl pH 8.0, 150 mM NaCl, 1 mM EDTA, 2% Triton X-100, 0.5% sodium deoxycholate, 0.1% SDS) supplemented with protease inhibitor cocktail for 30 min, followed by centrifugation at 16, 000 x *g* for 10 min at 4°C. Cleared lysates were diluted in 2 x Laemmli sample buffer and analyzed by immunoblotting for LC3 and p62.

#### Immunofluorescence (IF) and confocal microscopy

Cells seeded on coverslips were washed with PBS and fixed with 4% (v/v) paraformaldehyde (Thermo Fisher Scientific) for 20 min at RT. Cells were then permeabilized in 0.2% (v/v) Triton X-100 in PBS for 10 min and blocked with PBS containing 10% goat serum for 1 h at room temperature (RT). Primary antibodies diluted in 1% goat serum were applied to the cells for 1.5 h at RT or overnight at 4°C. After three washes with PBS containing 0.25% Tween-20, cells were incubated with Alexa 488-, Alexa 546-, Alexa 568-, and/or Alexa 647-conjugated secondary antibodies (Invitrogen, 1:800) in 1% goat serum for 1 h at RT for confocal microscopy analysis. Following another round of PBS washes, nuclei were counterstained with DAPI (Thermo Scientific) and coverslips were mounted using VECTASHIELD® PLUS Antifade Mounting Medium (Vector Laboratories).

For immunofluorescent staining of tissue sections, paraffine-embedded sections were deparaffinized in xylene, rehydrated through a graded ethanol series, and rinsed in distilled water. Antigen retrieval was performed by incubating slides in DAKO target retrieval solution (EDTA, pH 9.0) using a pressure cooker at 95°C for 20 minutes, followed by cooling at RT for 15 min. Slides were washed 3 times in PBS containing 0.1 M glycine (IF wash buffer) and then blocked in 10% goat serum (in IF wash buffer) for 1 h at RT. Sections were incubated overnight at 4°C with primary antibodies diluted in blocking solution. After 3 washes in IF wash buffer, slides were incubated with fluorescently conjugated secondary antibodies (Invitrogen, 2μg/ml) for 30 min at RT in the dark. Slides were then washed and incubated with an autofluorescence quenching reagent followed by nuclear staining with DAPI (1: 10,000 in PBS) for 5 min. For sequential staining, coverslips were carefully removed, and antibody stripping was performed by incubating slides in citrate buffer (pH 6.0) in a steamer for 20 minutes. After cooling, slides were re-blocked and incubated with the second primary antibody overnight at 4°C, followed by incubation with the corresponding secondary antibody for 30 min. After final washes, slides were counterstained with DAPI, washed, and mounted using VECTASHIELD Vibrance mounting medium (Vector Laboratories). Slides were imaged at 40X magnification using a Hamamatsu Nanozoomer S60 scanner (Hamamatsu Photonics K.K., Shizuoka, Japan).

Confocal microscopy was performed on a Leica TCS SP5 II laser-scanning confocal microscope (Leica Microsystems) equipped with HyD detectors and Leica PMT detectors; four laser (405 nm, 488 nm, 561 nm, and 633 nm) were used for fluorescence imaging in the blue, green, red, and far-red channels, respectively, combined with their respective standard emission filter sets. Images were acquired using the HCX PL APO CS (63x, 1.4NA, oil immersion) objective and the Leica LAS-X software (v4.8.0.28989). Confocal z-stacks were acquired with optimal z-intervals according to the Nyquist criterion, covering the whole volume of cells from the basal to the apical region. Images were then compiled by ‘max projection’ before analysis in ImageJ.

#### Image processing and analysis

All image analyses were conducted using ImageJ Fiji (v.2.9.0, National Institutes of Health). Maximum intensity projections were generated from deconvoluted z-stack confocal images. Fluorescence intensity was quantified by drawing line scans across regions of interest and applying the ‘Plot Profile’ function. Intensity values for each channel were normalized to their respective maximum value to enable cross-sample comparisons. For individual cell measurements, cells were manually cropped and background signals were set to zero. The cropped images were converted into binary format, and nuclei were segmented using the DAPI signal *via* the Intensity Threshold method. Settings for segmentation and filtering were applied consistently across all datasets

For the measurement of NOTCH1ΔE colocalization with UVRAG and/or endosomal markers, intensity threshold segmentation was applied to individual cells to identify regions corresponding to NOTCH1ΔE, UVRAG, and/or endosomes (EEA1, LAMP1). Colocalization analysis was performed using the “JACoP” plugin in ImageJ to calculate Pearson’s correlation coefficient (to assess the linear intensity correlation between UVRAG and ligand-stimulated NOTCH1, Figure 2C) or Manders’ coefficient (to quantify the proportion of NOTCH1ΔE that overlapped with LAMP1, Figure 4F). At least 30 cells, randomly selected from 5-10 HPF across three independent experiments, were analyzed.

#### Flow cytometric analysis and cell sorting

Single-cell suspensions were prepared from bone marrow (femur and tibia), spleen, and peripheral blood from adult mice and red blood cells were lysed with ACK lysing buffer (Quality Biological). To block nonspecific antibody binding, cells were incubated with anti-CD16/32 antibody (TruStain FcX™, BioLegend; 1:200) for 15 min on ice. Cell staining was performed using fluorescently conjugated antibodies against mouse CD4-APC (BD Pharmingen^TM^; 1:400), mouse CD8a-PE-Cy™7 (BD Pharmingen^TM^; 1:400), or RB780 rat IgG2a, κ isotype control (BD Horizon™; 1:400), and gated on GFP^+^. GFP-positive leukemic cells were also isolated from spleen using a FACSymphony S6 SE cell sorter (BD Biosciences) for limiting dilution experiments. For human T-ALL xenograft analysis, single cells were stained with antibodies against human CD45-APC (clone HI30, BioLegend; 1:400) and/or human CD7-PE (clone CD7-6B7, BioLegend; 1:400). After staining, cells were fixed using BD Cytofix™ fixation buffer (BD Biosciences); live cells were gated and analyzed on a BD FACSymphony™ A5 SE cytometer and FACSDiva software (BD Biosciences). Data were further analyzed with FlowJo v.10.

#### Histology and immunohistochemistry

Peripheral blood smears were fixed in methanol and stained with Wright-Giemsa solution. Slides were rinsed with distilled water, air-dried, and mounted with Cytoseal 60 (VWR), and coverslipped. Tissue specimens were harvested from mice, fixed overnight in 10% neutral buffered formalin (VWR), dehydrated, and embedded in paraffin. Paraffine blocks were sectioned at 5 μm thickness and stained with hematoxylin and eosin (H&E) using a Dakewe automated stainer. For analysis of intestinal goblet cells, sections were stained using the periodic acid-Schiff (PAS) staining system (Sigma-Aldrich). Briefly, deparaffinized sections were oxidized in 0.5% periodic acid for 15 min, rinsed and incubated for another 5 min in distilled water, treated with Schiff’s reagent for 15 min, and counterstained with hematoxylin. Slides were dehydrated, cleared, and mounted. PAS-positive goblet cells were quantified per crypt using bright-field microscopy in a blinded manner.

For immunochemistry, tissue slides were deparaffinized by immersion in xylene and rehydrated by serial immersion in 100%, 95%, 70% alcohol and distilled water. Endogenous peroxidase was blocked with 3% hydrogen peroxide. Antigen retrieval was performed by incubating the sections in boiling 10 mM sodium citrate pH 6.0, followed by incubation with the indicated primary antibody overnight at 4°C. Antibody binding was detected with EnVision^TM^ Dual Link System-HRP DAB kit (Dako). Sections were then counterstained with hematoxylin. For negative controls, the primary antibody was excluded. For evaluation and scoring of immunohistochemical data, we randomly selected 10 fields within the tumor area under high power magnification (40x) for evaluation. The investigators conducted blind counting for all quantification.

#### RNA-Seq

Total RNA was extracted from splenic leukemia cells isolated from bone marrow-transplanted mice using Trizol reagent (Invitrogen), according to the manufacturer’s protocol. Extracted RNA was treated with DNase I for 30 min. Subsequently, 500 ng of RNA was used to generate 3’mRNA-Seq libraries using the QuantSeq 3’mRNA-Seq V2 Library Prep Kit FWD with UDI (UDI12A_0001-0096; Lexogen) for Illumina, according to the manufacturer’s protocol. Library quality and fragment size were assessed using an Agilent Tapestation 4150 (Agilent Technologies) and quantified with a Qubit 3.0 fluorometer (Thermo Scientific). Libraries were pooled and sequenced on an Illumina NextSeq 2000 using high-output, single-end 75-bp reads. Sequencing reads were aligned to the mouse reference genome (GRCm38) using the STAR aligner (Dobin et al., 2013) with default parameters. Gene-level counts were obtained using the *featurecounts* function from the *R*subread package in *R* (Liao et al., 2019). Differential gene expression analysis was performed using DESeq2 (Anders and Huber, 2010). Genes with an average read count below ten or flagged as outliers were excluded. Differential expression plots for significant genes (*p* <0.05, [log_2_FC]>1) were generated using the limma and ggplot2 packages in *R*.

#### Gene set enrichment analysis (GSEA)

GSEA was performed using the clusterProfiler *R* package (Wu et al., 2021). Genes were ranked by the metric Sign(log_2_FC)x[(-log(p-value)], derived from differential expression analysis. Enrichment scores and statistical significance were calculated using 1,000 phenotype permutation. Gene sets used in this study were identified from the Molecular Signatures Database (MSigDB v3.0 Curated Collection) and previously published sources (Li et al., 2021; Liberzon et al., 2015; Palomero et al., 2006b; Ramalho-Santos et al., 2002).

Single-sample GSEA (ssGSEA) was performed using the GSVA *R* package (v1.50.0) on normalized expression matrices (log₂TPM or variance-stabilized counts). Enrichment scores were computed for each sample using a predefined NOTCH1 gene signature composed of canonical NOTCH1 transcriptional targets^21^. ssGSEA scores were Z-score normalized for visualization and correlation analyses. For heatmaps or scatter plot visualization, samples were grouped by median thresholds of ssGSEA scores or relevant gene expression levels. Correlations with pathway components or clinical parameters were assessed using *Pearson* or *Spearman* correlation, depending on data distribution.

Cross-species pathway analysis was conducted to compare GSEA results between human and mouse datasets. Specifically, we analyzed the human T-ALL dataset (GSE13159) and our UVRAG-inducible T-ALL mouse model using GSEA applied independently to each species. Enrichment were performed on hallmark gene sets (MSigDB Hallmark (H) collection, and results were filtered to retain only those pathways detected in both datasets. For each overlapping pathway, the normalized enrichment scores (NES) and adjusted *p*-values (*padj*) were extracted. To quantify pathway significance across species, −log_10_(*padj*) values were calculated for both human and mouse data and plotted in a two-dimensional correlation scatter plot using the ggplot2 *R* package. Pathways were categorized as “upregulated” if NES > 0 and *padj* < 0.05 in both species, “downregulated” if NES < 0 and *padj* < 0.05 in both species, and “not consistent” if they did not meet the above criteria. Selected hallmark pathways were manually annotated in the plots to highlight relevant biological processes.

#### Identification of ESCRT gene signature correlated with UVRAG/ITCH and NOTCH1 activity

TCGA RNA-sequencing expression was obtained from the Xena browser (https://xenabrowser.net; ref.(Goldman et al., 2020)) and the pan-cancer clinical data through a recent harmonization project (Liu et al., 2018). Cancer types with at least 200 samples with both gene expression and clinical information were included (BRCA, LUAD, LGG, HNSC, PRAD, THCA, LUSC, SKCM, STAD, BLCA, KIRC, LIHC, CESC, KIRP, COAD, and SARC). To evaluate NOTCH activity status, the average z-score of *HES1* and *DTX1* was used as an indirect readout. In addition, the average z-score of *UVRAG* and *ITCH* was used to evaluate UVRAG/ITCH levels. For *ESCRT* expression, 30 genes were considered*: MVB12A, CHMP6, VPS4B, TSG101, VPS28, HGS, CHMP1A, CHMP3, VPS37C, UBAP1, CHMP2A, SNF8, STAM, CHMP4A, CHMP4B, CHMP5, VPS37A, CHMP2B, IST1, CHMP4C, STAM2, VPS36, VPS4A, MVB12B, VPS37B, CHMP1B, CHMP7, VPS25, VTA1, VPS37D.* The expression of each ESCRT gene was correlated with NOTCH activity and UVRAG/ITCH levels, in every cancer type (through *Spearman* rank correlation). ESCRT genes with negative median correlation across cancer types with NOTCH activity, and a positive median correlation across cancer types with UVRAG/ITCH, were selected for ESCRT signature. This resulted in a list of 13 ESCRT genes: *STAM2*, *CHMP3*, *STAM*, *TSG101*, *CHMP5*, *VPS36*, *UBAP1*, *CHMP1B*, *CHMP2B*, *VPS4B*, *VPS37A*, *CHMP4C*, and *VTA1*. We selected five genes (*TSG101*, *VPS37A*, *STAM*, *CHMP3*, and *VPS4B*) with the highest average correlation coefficient across different cancer types and intersection with T-ALL results. The five-gene score was correlated with survival across TCGA cancer types. The log-rank test (evaluated using log rank_test function from the lifelines statistics package) was computed, comparing survival based on ESCRT gene signature (average z-score), with median value to separate the curves. Significant cancer types were selected as those having a log-rank FDR corrected *q* < 0.1. Kaplan-Meier (KM) curves were plotted using Kaplan Meier Fitter from the lifelines Python package.

### QUANTIFICATION AND STATISTICAL ANALYSIS

Statistical analysis was performed using GraphPad Prism 10. Data are expressed as mean ± standard deviation (s.d.) or standard error of the mean (s.e.m.) for the number of replicates, as indicated in the figure legend. All experiments were independently repeated at least three times. The Student’s *t*-test was used for comparisons between the two groups. For multiple groups, analysis of variance (ANOVA) with Tukey’s *post hoc* test or Kruskal-Wallis test with *post hoc* Dunn’s test were applied, as specified in the respective figure legends. A two-sided *p* value of ≤ 0.05 was considered statistically significant (\**p* < 0.05, \*\**p* < 0.01, \*\*\**p* < 0.001, ****, *p* < 0.0001, and n.s., not significant).

## REFERENCES

Aifantis, I., Raetz, E., and Buonamici, S. (2008). Molecular pathogenesis of T-cell leukaemia and lymphoma. Nat Rev Immunol 8, 380–390.

Anders, S., and Huber, W. (2010). Differential expression analysis for sequence count data. Genome Biol 11, R106.

Armstrong, F., Brunet de la Grange, P., Gerby, B., Rouyez, M.C., Calvo, J., Fontenay, M., Boissel, N., Dombret, H., Baruchel, A., Landman-Parker, J., et al. (2009). NOTCH is a key regulator of human T-cell acute leukemia initiating cell activity. Blood 113, 1730–1740.

Belteki, G., Haigh, J., Kabacs, N., Haigh, K., Sison, K., Costantini, F., Whitsett, J., Quaggin, S.E., and Nagy, A. (2005). Conditional and inducible transgene expression in mice through the combinatorial use of Cre-mediated recombination and tetracycline induction. Nucleic Acids Res 33, e51.

Chapman, G., Major, J.A., Iyer, K., James, A.C., Pursglove, S.E., Moreau, J.L., and Dunwoodie, S.L. (2016). Notch1 endocytosis is induced by ligand and is required for signal transduction. Biochim Biophys Acta 1863, 166–177.

Chastagner, P., Israel, A., and Brou, C. (2008). AIP4/Itch regulates Notch receptor degradation in the absence of ligand. PLoS One 3, e2735.

Chiang, M.Y., Xu, L., Shestova, O., Histen, G., L’Heureux, S., Romany, C., Childs, M.E., Gimotty, P.A., Aster, J.C., and Pear, W.S. (2008). Leukemia-associated NOTCH1 alleles are weak tumor initiators but accelerate K-ras-initiated leukemia. J Clin Invest 118, 3181–3194.

Cox, C.V., Evely, R.S., Oakhill, A., Pamphilon, D.H., Goulden, N.J., and Blair, A. (2004). Characterization of acute lymphoblastic leukemia progenitor cells. Blood 104, 2919–2925.

De Keersmaecker, K., Lahortiga, I., Mentens, N., Folens, C., Van Neste, L., Bekaert, S., Vandenberghe, P., Odero, M.D., Marynen, P., and Cools, J. (2008). In vitro validation of gamma-secretase inhibitors alone or in combination with other anti-cancer drugs for the treatment of T-cell acute lymphoblastic leukemia. Haematologica 93, 533–542.

Dobin, A., Davis, C.A., Schlesinger, F., Drenkow, J., Zaleski, C., Jha, S., Batut, P., Chaisson, M., and Gingeras, T.R. (2013). STAR: ultrafast universal RNA-seq aligner. Bioinformatics 29, 15–21.

Doody, R.S., Raman, R., Farlow, M., Iwatsubo, T., Vellas, B., Joffe, S., Kieburtz, K., He, F., Sun, X., Thomas, R.G., et al. (2013). A phase 3 trial of semagacestat for treatment of Alzheimer’s disease. N Engl J Med 369, 341–350.

Dovey, H.F., John, V., Anderson, J.P., Chen, L.Z., de Saint Andrieu, P., Fang, L.Y., Freedman, S.B., Folmer, B., Goldbach, E., Holsztynska, E.J., et al. (2001). Functional gamma-secretase inhibitors reduce beta-amyloid peptide levels in brain. J Neurochem 76, 173–181.

Eng, J.K., McCormack, A.L., and Yates, J.R. (1994). An approach to correlate tandem mass spectral data of peptides with amino acid sequences in a protein database. Journal of the American Society for Mass Spectrometry 5, 976–989.

Ferrando, A.A. (2009). The role of NOTCH1 signaling in T-ALL. Hematology Am Soc Hematol Educ Program, 353–361.

Fortini, M.E., and Bilder, D. (2009). Endocytic regulation of Notch signaling. Curr Opin Genet Dev 19, 323–328.

Fuwa, T.J., Hori, K., Sasamura, T., Higgs, J., Baron, M., and Matsuno, K. (2006). The first deltex null mutant indicates tissue-specific deltex-dependent Notch signaling in Drosophila. Mol Genet Genomics 275, 251–263.

Goldman, M.J., Craft, B., Hastie, M., Repecka, K., McDade, F., Kamath, A., Banerjee, A., Luo, Y., Rogers, D., Brooks, A.N., et al. (2020). Visualizing and interpreting cancer genomics data via the Xena platform. Nat Biotechnol 38, 675–678.

Habets, R.A., de Bock, C.E., Serneels, L., Lodewijckx, I., Verbeke, D., Nittner, D., Narlawar, R., Demeyer, S., Dooley, J., Liston, A., et al. (2019). Safe targeting of T cell acute lymphoblastic leukemia by pathology-specific NOTCH inhibition. Sci Transl Med 11.

Henne, W.M., Buchkovich, N.J., and Emr, S.D. (2011). The ESCRT pathway. Dev Cell 21, 77–91.

Holmes, R., and Zuniga-Pflucker, J.C. (2009). The OP9-DL1 system: generation of T-lymphocytes from embryonic or hematopoietic stem cells in vitro. Cold Spring Harb Protoc 2009, pdb prot5156.

Hurley, J.H., and Emr, S.D. (2006). The ESCRT complexes: structure and mechanism of a membrane-trafficking network. Annu Rev Biophys Biomol Struct 35, 277–298.

Itakura, E., Kishi, C., Inoue, K., and Mizushima, N. (2008). Beclin 1 forms two distinct phosphatidylinositol 3-kinase complexes with mammalian Atg14 and UVRAG. Molecular biology of the cell 19, 5360–5372.

Izon, D.J., Punt, J.A., Xu, L., Karnell, F.G., Allman, D., Myung, P.S., Boerth, N.J., Pui, J.C., Koretzky, G.A., and Pear, W.S. (2001). Notch1 regulates maturation of CD4+ and CD8+ thymocytes by modulating TCR signal strength. Immunity 14, 253–264.

Kandachar, V., and Roegiers, F. (2012). Endocytosis and control of Notch signaling. Curr Opin Cell Biol 24, 534–540.

King, B., Trimarchi, T., Reavie, L., Xu, L., Mullenders, J., Ntziachristos, P., Aranda-Orgilles, B., Perez-Garcia, A., Shi, J., Vakoc, C., et al. (2013). The ubiquitin ligase FBXW7 modulates leukemia-initiating cell activity by regulating MYC stability. Cell 153, 1552–1566.

Knoechel, B., Bhatt, A., Pan, L., Pedamallu, C.S., Severson, E., Gutierrez, A., Dorfman, D.M., Kuo, F.C., Kluk, M., Kung, A.L., et al. (2015). Complete hematologic response of early T-cell progenitor acute lymphoblastic leukemia to the gamma-secretase inhibitor BMS-906024: genetic and epigenetic findings in an outlier case. Cold Spring Harb Mol Case Stud 1, a000539.

Kopan, R., and Ilagan, M.X. (2009). The canonical Notch signaling pathway: unfolding the activation mechanism. Cell 137, 216–233.

Kourtis, N., Lazaris, C., Hockemeyer, K., Balandran, J.C., Jimenez, A.R., Mullenders, J., Gong, Y., Trimarchi, T., Bhatt, K., Hu, H., et al. (2018). Oncogenic hijacking of the stress response machinery in T cell acute lymphoblastic leukemia. Nat Med 24, 1157–1166.

Lee, G., Liang, C., Park, G., Jang, C., Jung, J.U., and Chung, J. (2011). UVRAG is required for organ rotation by regulating Notch endocytosis in Drosophila. Dev Biol 356, 588–597.

Li, Y., Li, K., Zhu, L., Li, B., Zong, D., Cai, P., Jiang, C., Du, P., Lin, J., and Qu, K. (2021). Development of double-positive thymocytes at single-cell resolution. Genome medicine 13, 49.

Liang, C., Feng, P., Ku, B., Dotan, I., Canaani, D., Oh, B.H., and Jung, J.U. (2006). Autophagic and tumour suppressor activity of a novel Beclin1-binding protein UVRAG. Nature cell biology 8, 688–699.

Liang, C., Lee, J.S., Inn, K.S., Gack, M.U., Li, Q., Roberts, E.A., Vergne, I., Deretic, V., Feng, P., Akazawa, C., et al. (2008). Beclin1-binding UVRAG targets the class C Vps complex to coordinate autophagosome maturation and endocytic trafficking. Nature cell biology 10, 776–787.

Liao, Y., Smyth, G.K., and Shi, W. (2019). The R package Rsubread is easier, faster, cheaper and better for alignment and quantification of RNA sequencing reads. Nucleic Acids Res 47, e47.

Liberzon, A., Birger, C., Thorvaldsdottir, H., Ghandi, M., Mesirov, J.P., and Tamayo, P. (2015). The Molecular Signatures Database (MSigDB) hallmark gene set collection. Cell Syst 1, 417–425.

Liu, J., Lichtenberg, T., Hoadley, K.A., Poisson, L.M., Lazar, A.J., Cherniack, A.D., Kovatich, A.J., Benz, C.C., Levine, D.A., Lee, A.V., et al. (2018). An Integrated TCGA Pan-Cancer Clinical Data Resource to Drive High-Quality Survival Outcome Analytics. Cell 173, 400–416 e411.

Ma, Z., Zeng, Y., Wang, M., Liu, W., Zhou, J., Wu, C., Hou, L., Yin, B., Qiang, B., Shu, P., et al. (2023). N4BP1 mediates RAM domain-dependent notch signaling turnover during neocortical development. EMBO J 42, e113383.

Matesic, L.E., Haines, D.C., Copeland, N.G., and Jenkins, N.A. (2006). Itch genetically interacts with Notch1 in a mouse autoimmune disease model. Hum Mol Genet 15, 3485–3497.

Mevissen, T.E., Hospenthal, M.K., Geurink, P.P., Elliott, P.R., Akutsu, M., Arnaudo, N., Ekkebus, R., Kulathu, Y., Wauer, T., El Oualid, F., et al. (2013). OTU deubiquitinases reveal mechanisms of linkage specificity and enable ubiquitin chain restriction analysis. Cell 154, 169–184.

Mizushima, N., Yoshimori, T., and Levine, B. (2010). Methods in mammalian autophagy research. Cell 140, 313–326.

O’Neil, J., Grim, J., Strack, P., Rao, S., Tibbitts, D., Winter, C., Hardwick, J., Welcker, M., Meijerink, J.P., Pieters, R., et al. (2007). FBW7 mutations in leukemic cells mediate NOTCH pathway activation and resistance to gamma-secretase inhibitors. J Exp Med 204, 1813–1824.

Palomero, T., Barnes, K.C., Real, P.J., Glade Bender, J.L., Sulis, M.L., Murty, V.V., Colovai, A.I., Balbin, M., and Ferrando, A.A. (2006a). CUTLL1, a novel human T-cell lymphoma cell line with t(7;9) rearrangement, aberrant NOTCH1 activation and high sensitivity to gamma-secretase inhibitors. Leukemia 20, 1279–1287.

Palomero, T., Lim, W.K., Odom, D.T., Sulis, M.L., Real, P.J., Margolin, A., Barnes, K.C., O’Neil, J., Neuberg, D., Weng, A.P., et al. (2006b). NOTCH1 directly regulates c-MYC and activates a feed-forward-loop transcriptional network promoting leukemic cell growth. Proc Natl Acad Sci U S A 103, 18261–18266.

Papayannidis, C., DeAngelo, D.J., Stock, W., Huang, B., Shaik, M.N., Cesari, R., Zheng, X., Reynolds, J.M., English, P.A., Ozeck, M., et al. (2015). A Phase 1 study of the novel gamma-secretase inhibitor PF-03084014 in patients with T-cell acute lymphoblastic leukemia and T-cell lymphoblastic lymphoma. Blood Cancer J 5, e350.

Peng, J., and Gygi, S.P. (2001). Proteomics: the move to mixtures. J Mass Spectrom 36, 1083–1091.

Piper, R.C., and Lehner, P.J. (2011). Endosomal transport via ubiquitination. Trends Cell Biol 21, 647–655.

Polonen, P., Di Giacomo, D., Seffernick, A.E., Elsayed, A., Kimura, S., Benini, F., Montefiori, L.E., Wood, B.L., Xu, J., Chen, C., et al. (2024). The genomic basis of childhood T-lineage acute lymphoblastic leukaemia. Nature 632, 1082–1091.

Ramalho-Santos, M., Yoon, S., Matsuzaki, Y., Mulligan, R.C., and Melton, D.A. (2002). “Stemness”: transcriptional profiling of embryonic and adult stem cells. Science 298, 597–600.

Rathinam, C., Matesic, L.E., and Flavell, R.A. (2011). The E3 ligase Itch is a negative regulator of the homeostasis and function of hematopoietic stem cells. Nat Immunol 12, 399–407.

Rossi, M., Rotblat, B., Ansell, K., Amelio, I., Caraglia, M., Misso, G., Bernassola, F., Cavasotto, C.N., Knight, R.A., Ciechanover, A., et al. (2014). High throughput screening for inhibitors of the HECT ubiquitin E3 ligase ITCH identifies antidepressant drugs as regulators of autophagy. Cell Death Dis 5, e1203.

Sakata, T., Sakaguchi, H., Tsuda, L., Higashitani, A., Aigaki, T., Matsuno, K., and Hayashi, S. (2004). Drosophila Nedd4 regulates endocytosis of notch and suppresses its ligand-independent activation. Curr Biol 14, 2228–2236.

Sanchez-Irizarry, C., Carpenter, A.C., Weng, A.P., Pear, W.S., Aster, J.C., and Blacklow, S.C. (2004). Notch subunit heterodimerization and prevention of ligand-independent proteolytic activation depend, respectively, on a novel domain and the LNR repeats. Mol Cell Biol 24, 9265–9273.

Schroeter, E.H., Kisslinger, J.A., and Kopan, R. (1998). Notch-1 signalling requires ligand-induced proteolytic release of intracellular domain. Nature 393, 382–386.

Schweisguth, F. (1995). Suppressor of Hairless is required for signal reception during lateral inhibition in the Drosophila pupal notum. Development 121, 1875–1884.

Shevchenko, A., Wilm, M., Vorm, O., and Mann, M. (1996). Mass spectrometric sequencing of proteins silver-stained polyacrylamide gels. Anal Chem 68, 850–858.

Sparrer, K.M.J., Gableske, S., Zurenski, M.A., Parker, Z.M., Full, F., Baumgart, G.J., Kato, J., Pacheco-Rodriguez, G., Liang, C., Pornillos, O., et al. (2017). TRIM23 mediates virus-induced autophagy via activation of TBK1. Nat Microbiol 2, 1543–1557.

Thompson, B.J., Buonamici, S., Sulis, M.L., Palomero, T., Vilimas, T., Basso, G., Ferrando, A., and Aifantis, I. (2007). The SCFFBW7 ubiquitin ligase complex as a tumor suppressor in T cell leukemia. J Exp Med 204, 1825–1835.

Vaccari, T., and Bilder, D. (2005). The Drosophila tumor suppressor vps25 prevents nonautonomous overproliferation by regulating notch trafficking. Dev Cell 9, 687–698.

van Tetering, G., van Diest, P., Verlaan, I., van der Wall, E., Kopan, R., and Vooijs, M. (2009). Metalloprotease ADAM10 is required for Notch1 site 2 cleavage. J Biol Chem 284, 31018–31027.

Wilkin, M., Tongngok, P., Gensch, N., Clemence, S., Motoki, M., Yamada, K., Hori, K., Taniguchi-Kanai, M., Franklin, E., Matsuno, K., et al. (2008). Drosophila HOPS and AP-3 complex genes are required for a Deltex-regulated activation of notch in the endosomal trafficking pathway. Dev Cell 15, 762–772.

Wilkin, M.B., Carbery, A.M., Fostier, M., Aslam, H., Mazaleyrat, S.L., Higgs, J., Myat, A., Evans, D.A., Cornell, M., and Baron, M. (2004). Regulation of notch endosomal sorting and signaling by Drosophila Nedd4 family proteins. Curr Biol 14, 2237–2244.

Wu, T., Hu, E., Xu, S., Chen, M., Guo, P., Dai, Z., Feng, T., Zhou, L., Tang, W., Zhan, L., et al. (2021). clusterProfiler 4.0: A universal enrichment tool for interpreting omics data. Innovation (Camb) 2, 100141.

Wu, X., Fleming, A., Ricketts, T., Pavel, M., Virgin, H., Menzies, F.M., and Rubinsztein, D.C. (2016). Autophagy regulates Notch degradation and modulates stem cell development and neurogenesis. Nat Commun 7, 10533.

Yamamoto, S., Charng, W.L., and Bellen, H.J. (2010). Endocytosis and intracellular trafficking of Notch and its ligands. Curr Top Dev Biol 92, 165–200.

Yang, G., Zhou, R., Zhou, Q., Guo, X., Yan, C., Ke, M., Lei, J., and Shi, Y. (2019). Structural basis of Notch recognition by human gamma-secretase. Nature 565, 192–197.

Yang, Y., He, S., Wang, Q., Li, F., Kwak, M.J., Chen, S., O’Connell, D., Zhang, T., Pirooz, S.D., Jeon, Y.H., et al. (2016). Autophagic UVRAG Promotes UV-Induced Photolesion Repair by Activation of the CRL4(DDB2) E3 Ligase. Molecular cell 62, 507–519.

Yeh, C.H., Bellon, M., Pancewicz-Wojtkiewicz, J., and Nicot, C. (2016). Oncogenic mutations in the FBXW7 gene of adult T-cell leukemia patients. Proc Natl Acad Sci U S A 113, 6731–6736.

Zhu, K., Shan, Z., Chen, X., Cai, Y., Cui, L., Yao, W., Wang, Z., Shi, P., Tian, C., Lou, J., et al. (2017). Allosteric auto-inhibition and activation of the Nedd4 family E3 ligase Itch. EMBO Rep 18, 1618–1630.

