## Supplemental Information for "A K27-linked Ubiquitin Checkpoint Controls NOTCH Homeostasis"

Supplemental information includes ten supplementary figures and two supplementary tables.

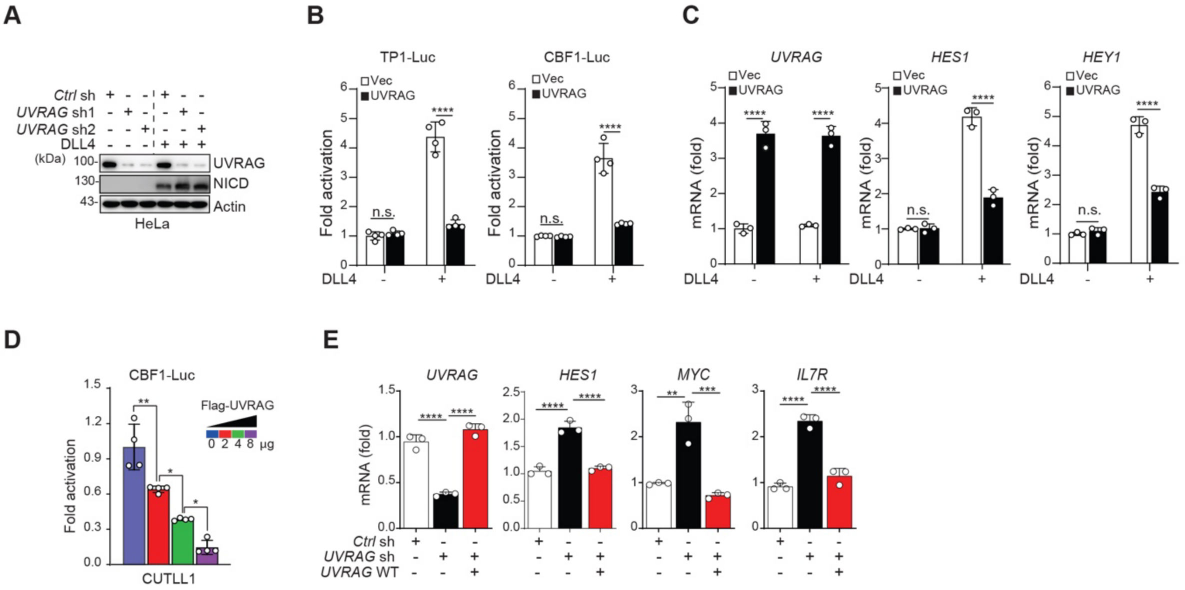


**Figure S1**

**Figure S1. UVRAG suppresses NOTCH1 activation, related to Figure 1**

(A) Immunoblotting (IB) analysis of UVRAG and NICD in HeLa cells transduced with *control* (*Ctrl*) or *UVRAG*-specific shRNAs (sh1 and sh2), with or without DLL4 stimulation. Actin serves as a loading control.

(B) TP1- (left) and CBF1- (right) luciferase reporter activity in HeLa cells stably expressing Vec or Flag-UVRAG, in the absence or presence of DLL4. n = 4.

(C) RT-qPCR quantification of *UVRAG*, *HES1*, and *HEY1* transcripts in cells from (B). n = 3.

(D) Dose-dependent inhibition of NOTCH signaling by UVRAG, measured by CBF1-luciferase activity in CUTLL1 cells transfected with increasing amounts of Flag-UVRAG. n = 4.

(E) RT-qPCR quantification of *UVRAG*, *HES1*, *MYC*, and *IL7R* mRNA in CUTLL1 cells expressing *Ctrl* shRNA or *UVRAG* shRNA, complemented with vector (*UVRAG* sh) or WT UVRAG (*UVRAG* WT). n = 3.

Data in (A) are representative of three independent experiments. Data in (B-E) are presented as mean ± s.d. from biologically independent samples and analyzed using one-way ANOVA with Tukey's *post hoc* test. *, $p < 0.05$; **, $p < 0.01$; ***, $p < 0.001$; ****, $p < 0.0001$; n.s., not significant.

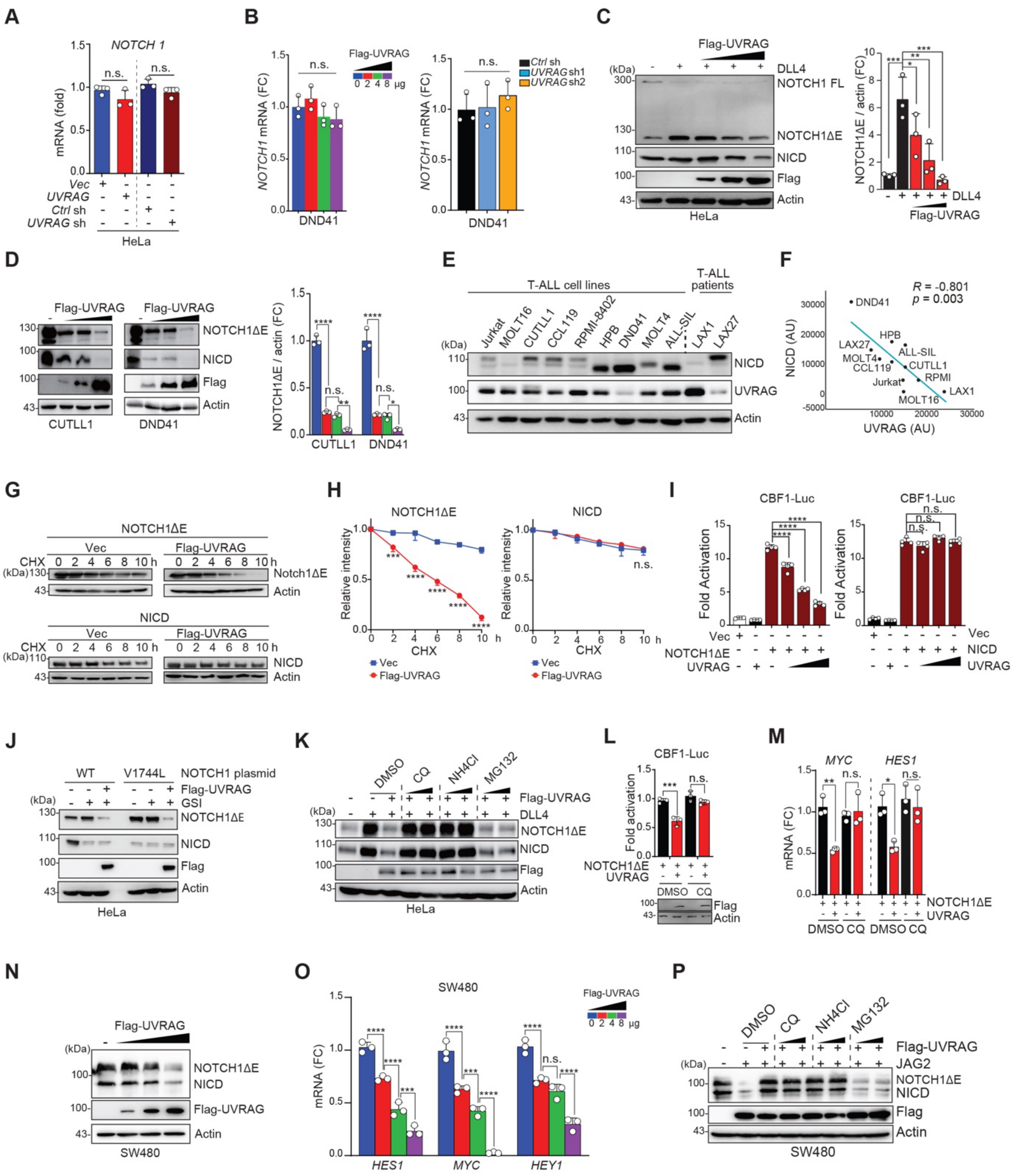


Figure S2

**Figure S2. UVRAG promotes lysosomal degradation of membrane-bound NOTCH1ΔE, related to Figure 1**

(A) RT-qPCR of *NOTCH1* transcripts in cells expressing *Ctrl* or *UVRAG* shRNA, or overexpressing Vector (Vec) or Flag-UVRAG.

(B) *NOTCH1* transcript levels in DND41 cells transfected with increasing amounts of Flag-UVRAG (left) or transduced with *Ctrl* or *UVRAG* shRNAs (sh1 and sh2) (right).

(C) IB (left) of NOTCH1 and densitometric quantification (right; n = 3) of NOTCH1ΔE/actin in HeLa cells transduced with increasing Flag-UVRAG and stimulated with DLL4. FL, full length.

(D) IB (left) of NOTCH1 and densitometric quantification (right; n=3) of NOTCH1ΔE/actin in T-ALL cells transduced with increasing amounts of Flag-UVRAG.

(E) Expression of NICD and UVRAG in human T-ALL cell lines and primary T-ALL bone marrow biopsies (LAX1, LAX27) by IB.

(F) Scatter plot showing correlation between NICD and UVRAG protein levels in samples from (E), quantified by densitometry and analyzed by *Pearson* correlation. AU, arbitrary unit.

(G and H) IB (G) and degradation kinetics (H) of NOTCH1ΔE and NICD in HEK293T cells transduced with indicated plasmids and treated with CHX over time.

(I) CBF1-luciferase activity in HEK293T cells co-transfected with NOTCH1ΔE (left) or NICD (right) with increasing UVRAG.

(J) IB of NOTCH1ΔE and NICD in HeLa cells co-expressing WT or V1744L NOTCH1 with Flag-UVRAG, treated w/ or w/o DAPT (GSI).

(K) IB of NOTCH1ΔE and NICD in HeLa cells expressing Flag-UVRAG, stimulated with DLL4 and treated with chloroquine (CQ), NH4Cl or MG132.

(L) CBF1-luciferase activity in HEK293T cells transfected with the NOTCH1ΔE with Vec or Flag-UVRAG (UVRAG +) and treated with DMSO or CQ.

(M) RT-qPCR of *MYC* and *HES1* transcripts in cells from (K).

(N) IB of NOTCH1ΔE and NICD in SW480 cells transfected with increasing Flag-UVRAG and stimulated with JAG2.

(O) RT-qPCR of indicated NOTCH1 target genes in cells from (N).

(P) IB of NOTCH1ΔE and NICD in SW480 cells treated with CQ, NH4Cl, or MG132 in the presence/absence of Flag-UVRAG and JAG2.

Data in (C-E), (G), (J), (K), (N), and (P) are the representative of three independent experiments. Data in (A-D), (H), (I), (L), (M), and (O) are presented as mean ± s.d. and analyzed using one-way ANOVA followed by Tukey's *post hoc* test. *, $p < 0.05$; **, $p < 0.01$; ***, $p < 0.001$; ****, $p < 0.0001$; n.s., not significant.

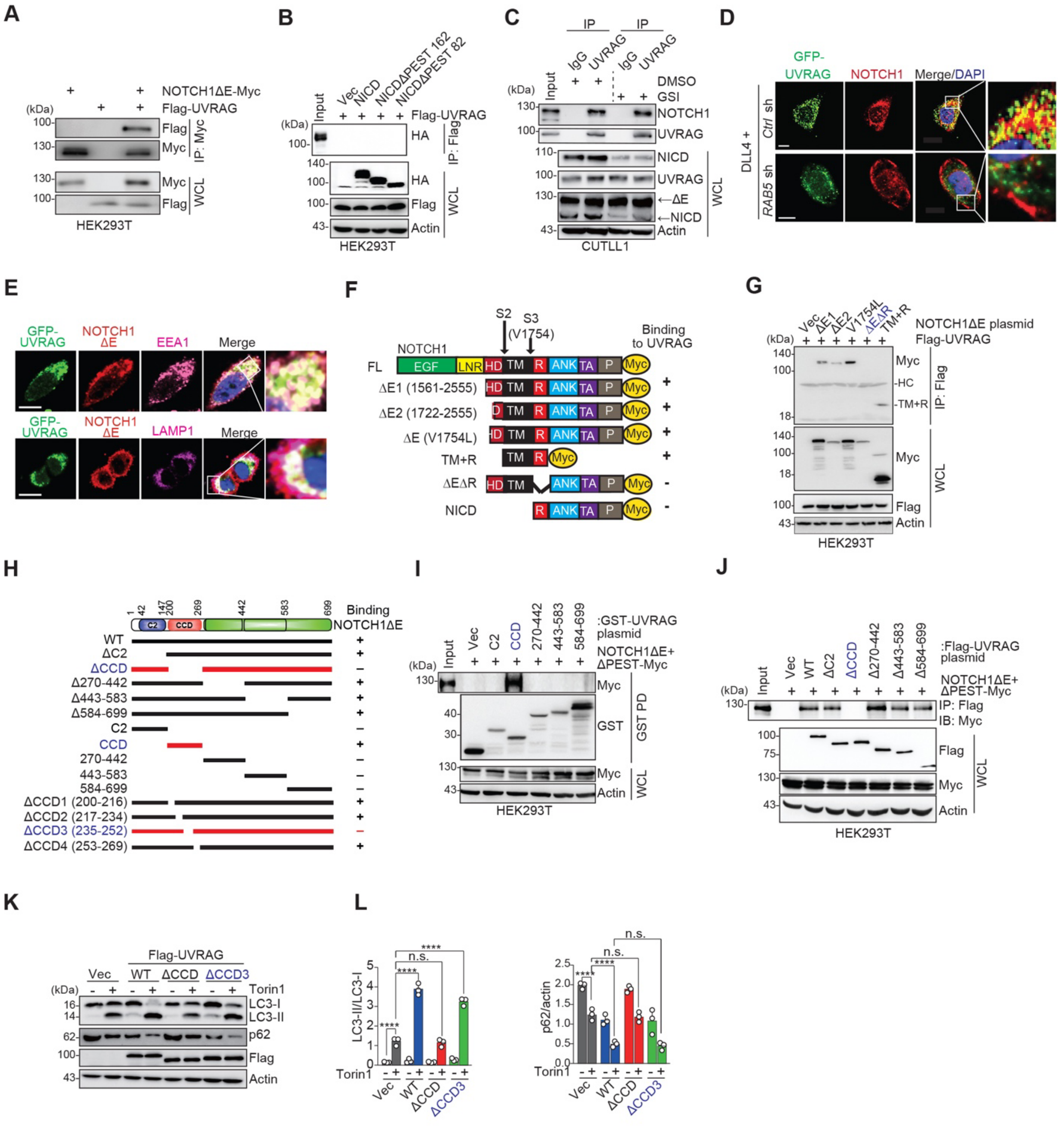


Figure S3

**Figure S3. UVRAG selectively binds membrane-tethered NOTCH1ΔE, related to Figure 2.**
(A and B) Co-IP of Flag-UVRAG with Myc-tagged NOTCH1ΔE (A) or with HA-tagged NICD (B) in cells transfected with the indicated plasmids.
(C) Co-IP of endogenous NOTCH1 with UVRAG in CUTLL1 cells treated with DMSO or DAPI (GSI). IgG, negative control.

(D) Representative confocal micrographs of colocalization of GFP-UVRAG (green) and endogenous NOTCH1 (red) in DLL4-stimulated HeLa cells transduced with *Ctrl* shRNA or *RAB5*-specific shRNA (*RAB5* sh). DAPI marks nuclei (blue). Scale bars, 10 mm.

(E) Representative confocal micrographs showing colocalization of GFP-UVRAG (green) and NOTCH1DE-Myc (red) with early endosomes (EEA1; magenta; top) or late endosomes/lysosomes (LAMP1; magenta; bottom) in HeLa cells. Insets highlight colocalization. Scale bars, 10 mm.

(F) Schematic of NOTCH1 constructs and their interactions with UVRAG. Domains: EGF, epidermal growth-factor-like repeats; LNR, Lin-12/NOTCH repeats; HD, heterodimerization domain; R, RAM domain; ANK, ankyrin repeats; TA, transactivation domain; P, PEST. S2 and S3 cleavage sites are marked. +, positive interaction; -, no interaction.

(G) co-IP of full-length or mutant NOTCH1ΔE constructs with UVRAG in HEK293T cells co-transfected with indicated plasmids. HC, heavy chain.

(H) Domain map of UVRAG constructs and summary of their interactions with NOTCH1ΔE. +, positive interaction; -, negative interaction.

(I and J) GST-pull down (PD) and co-IP of Myc-tagged NOTCH1ΔE+ΔPEST with GST-tagged UVRAG truncation (I) or deletion mutants (J) in HEK293T cells.

(K) IB analysis of LC3-I/LC3-II and p62 levels in HEK293T cells expressing Vec, UVRAG WT, or ΔCCD mutants, with or without Torin1 (3 h).

(L) Densitometric quantification of LC3-II/LC3-I ratio (left) and p62/actin ratio (right) in cells from K.

Data in (A-E), (G), and (I-K) are representative of three independent experiments. Data in (L) represent mean ± s.d. from 3 biologically independent samples, analyzed using one-way ANOVA with *post hoc* multiple comparisons. ****, $p < 0.0001$; n.s., not significant.

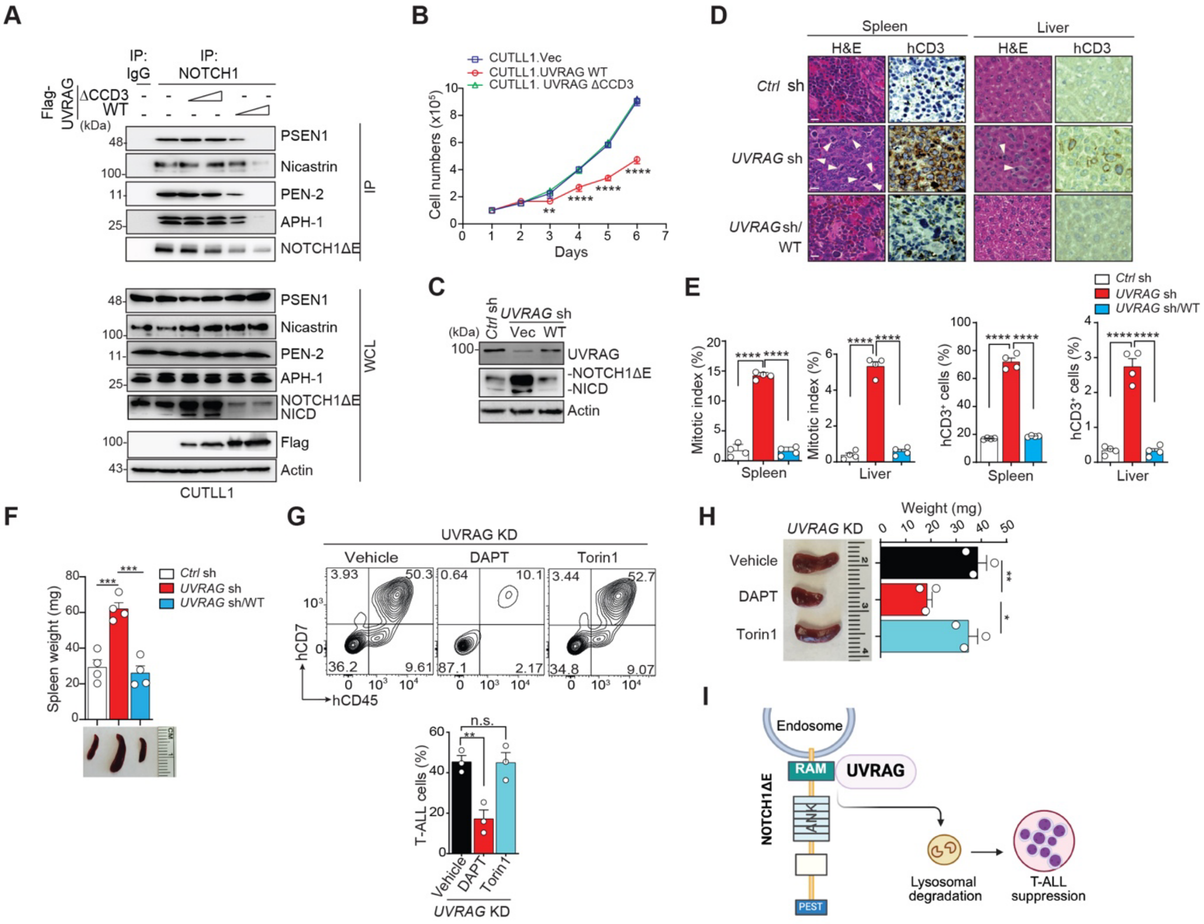


Figure S4

**Figure S4. UVRAG interaction with NOTCH1ΔE suppresses NOTCH1-driven T-ALL progression, related to Figure 2.**

(A) Co-IP of $\gamma$-secretase complex components (PSEN1, Nicastrin, PEN-2, APH-1) with NOTCH1 in CUTLL1 cells expressing Vec or increasing amounts of Flag-UVRAG WT or UVRAG ΔCCD3 mutant. IgG, negative control.

(B) Growth curve of CUTLL1 cells stably expressing Vec, UVRAG WT, or ΔCCD3 (n = 3).

(C) IB of NOTCH1ΔE and NICD in CUTLL1 cells stably expressing *Ctrl* shRNA or *UVRAG* shRNA complemented with Vec (*UVRAG* sh) or WT UVRAG (*UVRAG* sh/WT).

(D) H&E staining and immunohistochemical (IHC) for human CD3 (hCD3) in spleen (left) and liver (right) tissues from NSG mice bearing CUTLL1 xenografts expressing cells from (C). Arrows denote mitotic figures. Scale bars, 20 mm. Images representative of 5-6 mice per group.

(E) Quantification of mitotic figures and hCD3$^{+}$ cells in spleen and liver from (D).

(F) Spleen images and weight measurements at 33 days post-transplantation of mice injected with CUTLL1 cells described in (C) (n = 6).

(G) Flow cytometry plots (top) and quantification (bottom) of T-ALL (hCD45$^{+}$hCD7$^{+}$) cells in bone marrow of mice transplanted with *UVRAG*-knockdown (KD) CUTLL1 cells and treated with Vehicle, DAPT (10 mg kg$^{-1}$ per day, *i.p*), or Torin1 (20 mg kg$^{-1}$ per day, *i.p.*) for 21 days (n = 6-7 mice per group). (H) Spleen images and weight from mice in (G).

(I) Schematic model of UVRAG binding to the RAM domain of NOTCH1ΔE, driving lysosomal degradation and T-ALL suppression.

Data in (A), (C), (D), and (G) are representative of three independent experiments. Data in (B) and (E-H) are shown as mean ± s.d., analyzed using one-way ANOVA with Tukey's *post hoc* test. *, $p < 0.05$; **, $p < 0.01$; ***, $p < 0.001$; ****, $p < 0.0001$; n.s., not significant.

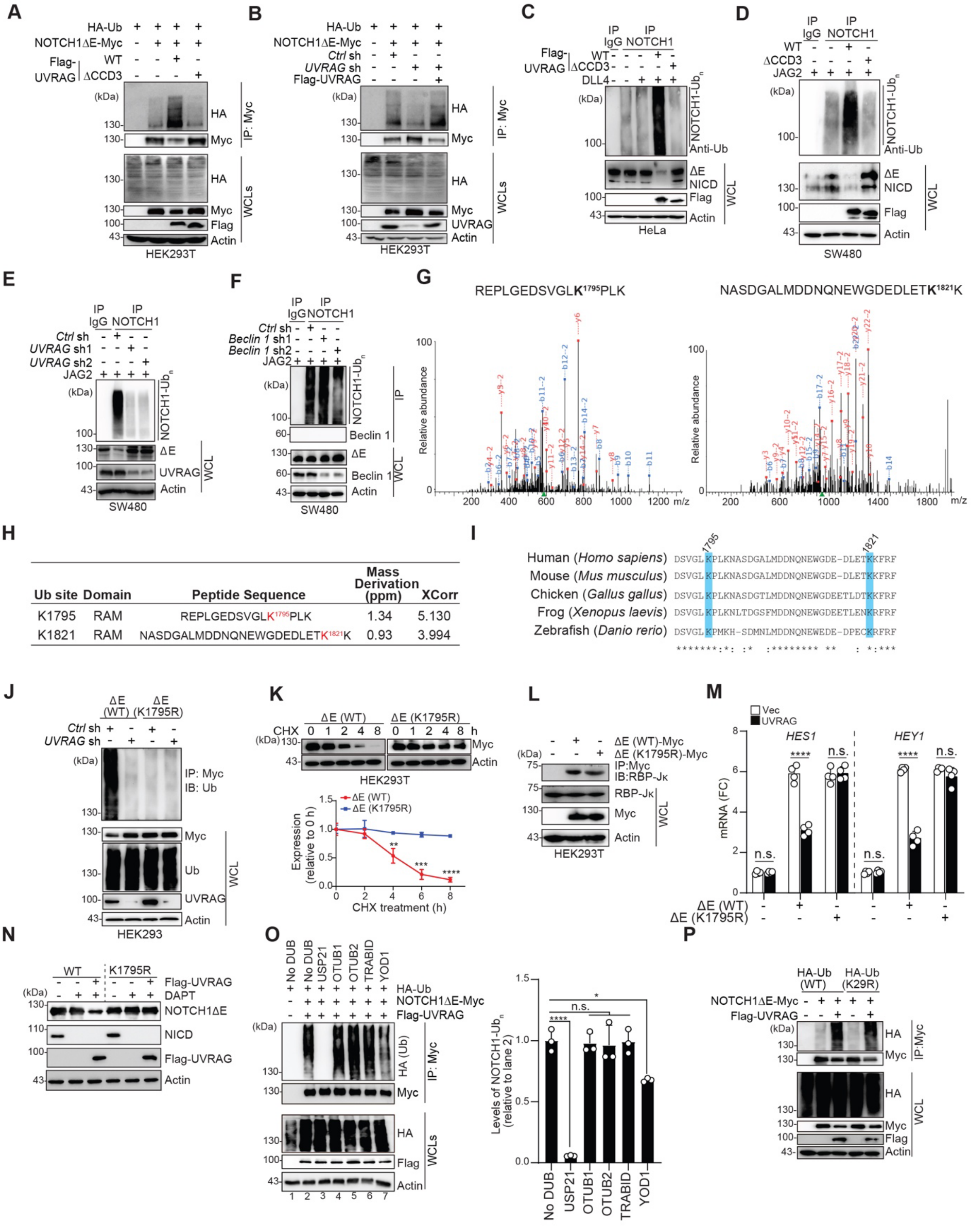


Figure S5

**Figure S5. UVRAG promotes ubiquitination of NOTCH1ΔE at lysine 1795 to drive its degradation, related to Figure 3.**

(A) IP of NOTCH1ΔE ubiquitination in cells transfected with HA-Ub, NOTCH1ΔE-Myc, and either Vec, Flag-UVRAG WT or ΔCCD3.

(B) Ubiquitination of NOTCH1ΔE in cells expressing *Ctrl* or UVRAG shRNAs, w/ or w/o rescue by Flag-UVRAG.

(C and D) IP of NOTCH1 ubiquitination in HeLa (C) or SW480 (D) cells expressing Vec, Flag-UVRAG WT or ΔCCD3, following DLL4 (C) or JAG2 (D) stimulation.

(E) Ubiquitination of NOTCH1 in JAG2-stimulated SW480 cells expressing *Ctrl* or *UVRAG* shRNAs.

(F) IP of NOTCH1 from JAG2-stimulated SW480 cells expressing *Ctrl* or *Beclin 1* shRNAs.

(G) Mass spectrometry spectra identifying K1795 and K1821 as UVRAG-regulated ubiquitination sites in NOTCH1ΔE. Annotated *b*- and *y*-ion are labeled.

(H) Sequences of identified peptides flanking ubiquitinated lysines, including peptide mass accuracy and cross-correlation (XCorr) values.

(I) Sequence alignment of *NOTCH1* orthologs across species showing conserved ubiquitinated lysines (cyan). *, conserved residues.

(J) Ubiquitination of NOTCH1ΔE WT and K1795R from cells expressing *Ctrl* or *UVRAG* shRNA.

(K) Degradation kinetics (top) and densitometric quantification (bottom) of NOTCH1ΔE WT and K1795R in CHX-treated cells (n=3).

(L) Co-IP of NOTCH1ΔE WT or K1795R with RBP-Jk.

(M) RT-qPCR of *HES1* and *HEY1* transcripts in HEK293T cells transfected with NOTCH1ΔE WT or K1795R and either Vec or Flag-UVRAG. n = 4 biological replicates.

(N) IB of NOTCH1ΔE and NICD in HeLa cells expressing WT or K1795R NOTCH1ΔE w/ or w/o DAPT. (O) *In vitro* DUB restriction assay of NOTCH1ΔE ubiquitination using lysates from HEK293T cells expressing HA-Ub, Flag-UVRAG, and NOTCH1ΔE-Myc. Purified proteins were incubated with indicated DUB; ubiquitinated species quantified by densitometry (right). n = 3.

(P) Ubiquitination of NOTCH1ΔE-Myc in HEK293T cells transfected with HA-Ub WT or K29R, NOTCH1ΔE-Myc, and Flag-UVRAG.

Data in (A-F), (J-L), and (N-P) are representative of three independent experiments. Data in (K) and (M) are shown as mean ± s.d., analyzed using analyzed using Student's two-tailed *t* test (K) or one-way ANOVA with Tukey's *post hoc* test (M). *, $p < 0.05$; **, $p < 0.01$; ***, $p < 0.001$; ****, $p < 0.0001$; n.s., not significant.

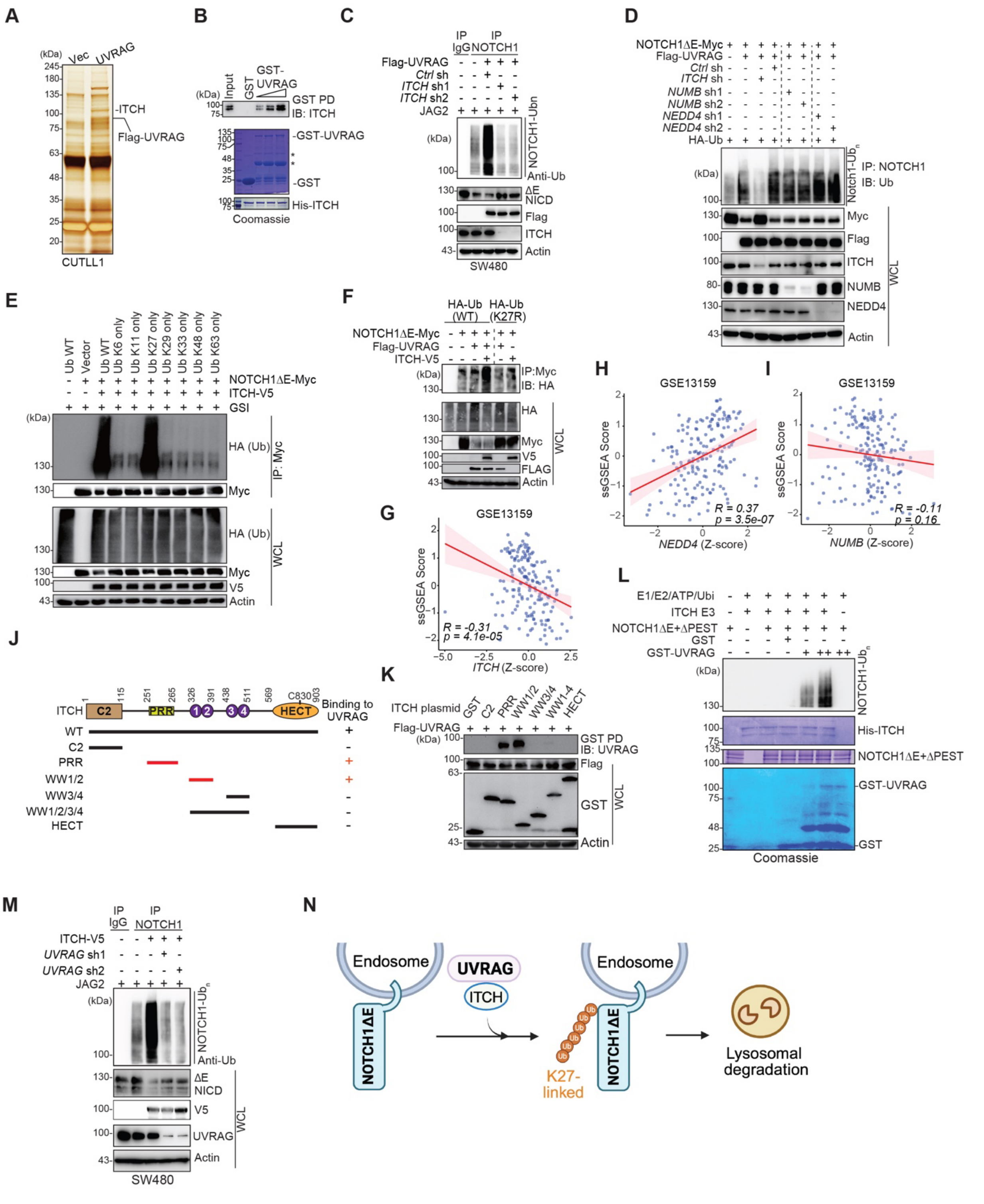


Figure S6

**Figure S6. UVRAG cooperates with ITCH to catalyze K27-linked ubiquitination of NOTCH1ΔE, related to Figure 3**

(A) Identification of UVRAG-interacting proteins by affinity purification. CUTLL1 cells stably expressing Vec or Flag-UVRAG were subjected to IP with anti-Flag agarose beads and the purified proteins were visualized by silver staining and identified by Mass Spectrometry.
(B) *In vitro* GST PD showing direct interaction between GST-UVRAG and His-ITCH. Coomassie staining shows input proteins.
(C) Ubiquitination of NOTCH1 in JAG2-stimulated SW480 cells expressing Flag-UVRAG and transduced with *Ctrl* or *ITCH* shRNAs (sh1 and sh2).
(D) Ubiquitination of NOTCH1ΔE-Myc in HEK293T cells co-transfected with Flag-UVRAG, HA-Ub, and *Ctrl* shRNA or shRNAs targeting *ITCH*, *NUMB*, or *NEDD4*.
(E) Ubiquitination of NOTCH1ΔE-Myc in HEK293T cells co-transfected with ITCH-V5 and either WT or lysine-mutant HA-Ub constructs, treated with compound E and assessed by IP with anti-Myc and IB with anti-HA.
(F) Ubiquitination of NOTCH1ΔE-Myc in HEK293T cells co-transfected with Flag-UVRAG, ITCH-V5, and either HA-Ub WT or the K27R mutant.
(G-I) Scatter plot showing correlation between *ITCH* (G), *NEDD4* (H), and *NUMB* (I) expression (Z-score) and ssGSEA scores of the NOTCH1 gene signature (Palomero et al., 2006b) in T-ALL dataset GSE13159 (n = 174).
(J) Schematic of ITCH domain architecture and truncation mutant tested for UVRAG interaction. Summary of UVRAG-binding capacity is indicated. +, positive interaction; -, no interaction.
(K) GST PD of Flag-UVRAG with GST-ITCH truncation constructs in HEK293T cells.
(L) *in vitro* ubiquitination assay showing UVRAG-dependent enhancement of ITCH-mediated NOTCH1ΔE ubiquitination. Purified GST-UVRAG, His-ITCH, and substrate (NOTCH1ΔE+ΔPEST) were mixed with ubiquitination components as indicated (bottom: Coomassie input; top: anti-Ub blot).
(M) Endogenous NOTCH1 ubiquitination in SW480 cells stably expressing ITCH-V5 and transduced with *Ctrl* or UVRAG shRNAs.
(N) Model depicting UVRAG-mediated recruitment and activation of ITCH to NOTCH1ΔE, promoting K27-linked ubiquitination and degradation.
Data in (B-F) and (K-M) are representative of three independent experiments.

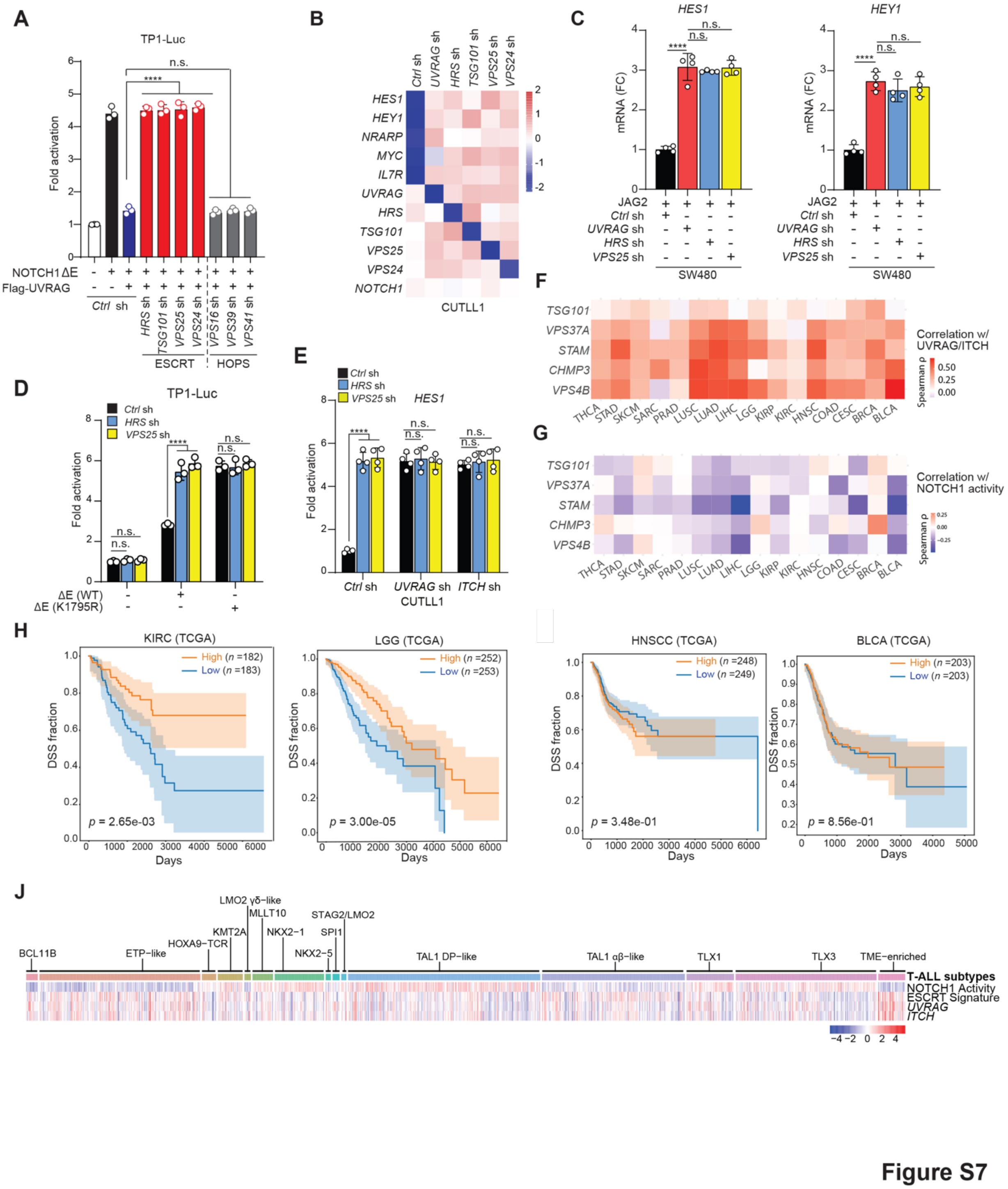


**Figure S7. K27-ubiquitinated NOTCH1ΔE is degraded through the ESCRT-mediated endo-lysosomal pathway, related to Figure 4.**

(A) TP1-luciferase activity in HEK293T cells co-transfected with NOTCH1ΔE-Myc and Flag-UVRAG along with *Ctrl* sh or gene-specific shRNAs as indicated.

(B) Heatmap showing expression changes in selected NOTCH1 target genes in CUTLL1 cells transduced with the indicated shRNAs.
(C) RT-qPCR analysis of *HES1* and *HEY1* in JAG2-stimulated SW480 cells transduced with the indicated shRNAs.
(D) TP1-luciferase activity in HEK293T cells expressing WT or K1795R-mutant NOTCH1ΔE and transduced with *Ctrl*, *HRS*, or *VPS25* shRNA.
(E) RT-qPCR of *HES1* in CUTLL1 cells expressing indicated shRNAs.
(F and G) Heatmap showing *Spearman* rank correlation coefficients for the expression of five core ESCRT genes (*TSG101*, *VPS37A*, *STAM*, *CHMP3*, and *VPS4B*), positively correlated with *UVRAG* and *ITCH* (F) and negatively correlated with NOTCH1 activity (averaging the levels of *HES1* and *DTX1*, G), across TCGA cancer types.
(H and I) Kaplan-Meier survival curves comparing the disease stable survival (DSS) between patients with a high *vs.* low (using the median as threshold) 5-gene ESCRT signature scores (average z-score) in TCGA KIRC (kidney renal clear cell carcinoma; H), LGG (low-grade glioma; H), HNSCC (head and neck squamous cell carcinoma; I), and BLCA (bladder urothelial carcinoma; I). Log-rank *p*-values are indicated.
(J) Heatmap showing the expression of NOTCH1 activity signature (Palomero et al., 2006b), ESCRT signature, *UVRAG*, and *ITCH* across 15 molecular subtypes of T-ALL in dataset dbGaP phs002276. Annotated T-ALL subtype classifications are indicated at the top.
Data in (A-E) are shown as mean ± s.d. of biologically independent samples and were analyzed using one-way ANOVA with Tukey's *post hoc* test. ****, $p < 0.0001$; n.s., not significant.

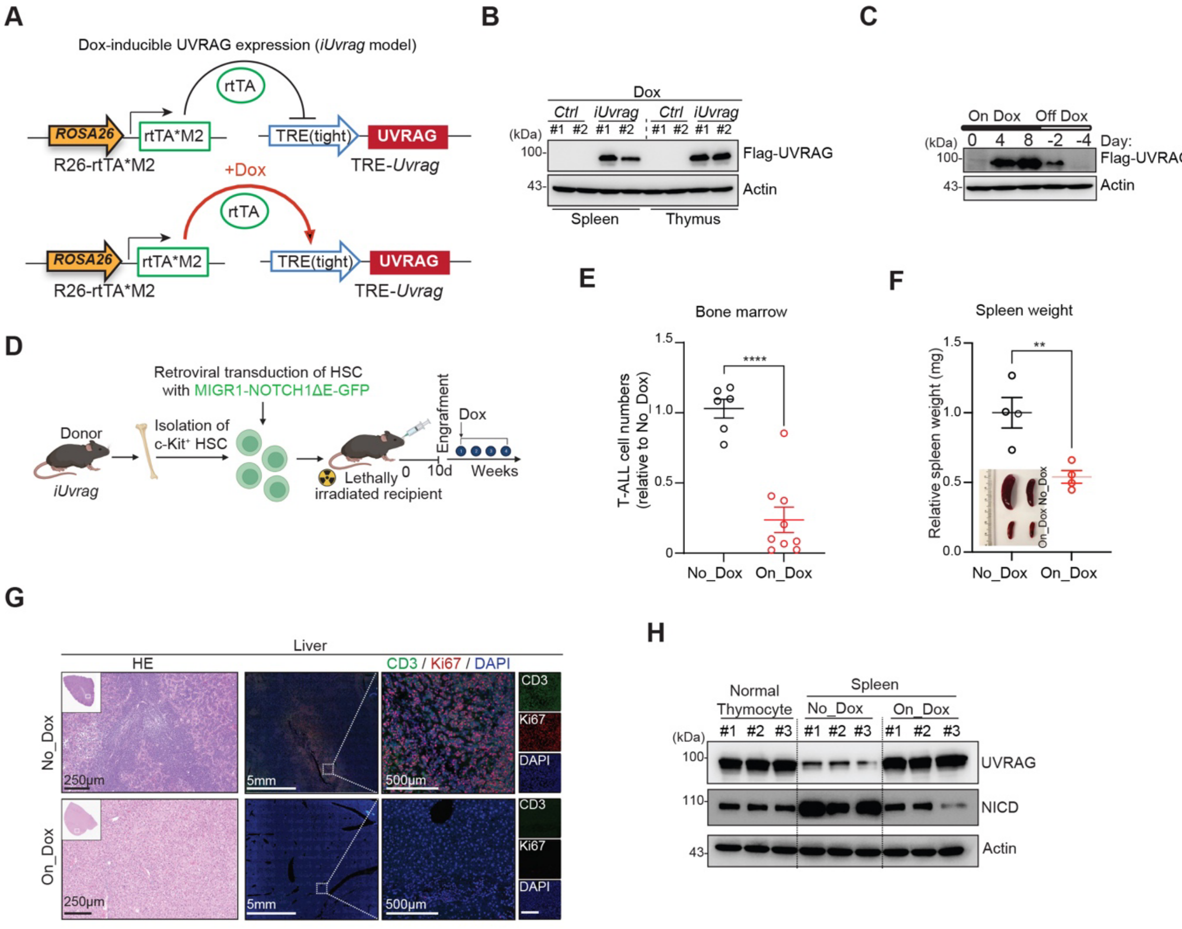


**Figure S8**

**Figure S8. UVRAG restrains NOTCH1-driven leukemia progression *in vivo*, related to Figure 5.**

(A) Schematic of the *Rosa-rtTA*M2* inducible *Uvrag* mouse model (*iUvrag*), where Dox treatment activates TRE-driven Flag-UVRAG expression.

(B) IB of Flag-UVRAG expression in spleens and thymi of control and *iUvrag* mice treated with Dox for 7 days.

(C) Time-course IB of Flag-UVRAG expression in spleens of *iUvrag* mice with Dox turned on or withdrawn (Off Dox) over the indicated days.

(D) Schematic of experimental design of NOTCH1ΔE-induced T-ALL using retroviral transduction of hematopoietic stem cells (HSCs) from *iUvrag* mice, followed by transplantation into irradiated recipients and induction of UVRAG expression by Dox.

(E) Quantification of GFP$^+$ T-ALL cell burden in the bone marrow (BM) of recipient mice treated with Dox (On_Dox; n = 9) or not (No_Dox; n = 6) for 4 weeks.

(F) Spleen weight measurements and representative spleen images from mice in (E) (n = 4 per group). (G) H&E staining and immunofluorescence of liver sections from mice in (E), showing CD3$^+$ T cells (green), Ki67$^+$ proliferating cells (red), and DAPI-stained nuclei (blue). Scale bars as indicated.

(H) IB of UVRAG and NICD expression in thymocytes from normal mice and in splenic T-ALL cells from *iUvrag* recipient mice treated with or without Dox.

Data in (B), (C), (G), and (H) are representative of three independent experiments. Data in (E and F) are shown as mean ± s.e.m. analyzed using Student's two-tailed *t* test. **, $p < 0.01$; ****, $p < 0.0001$.

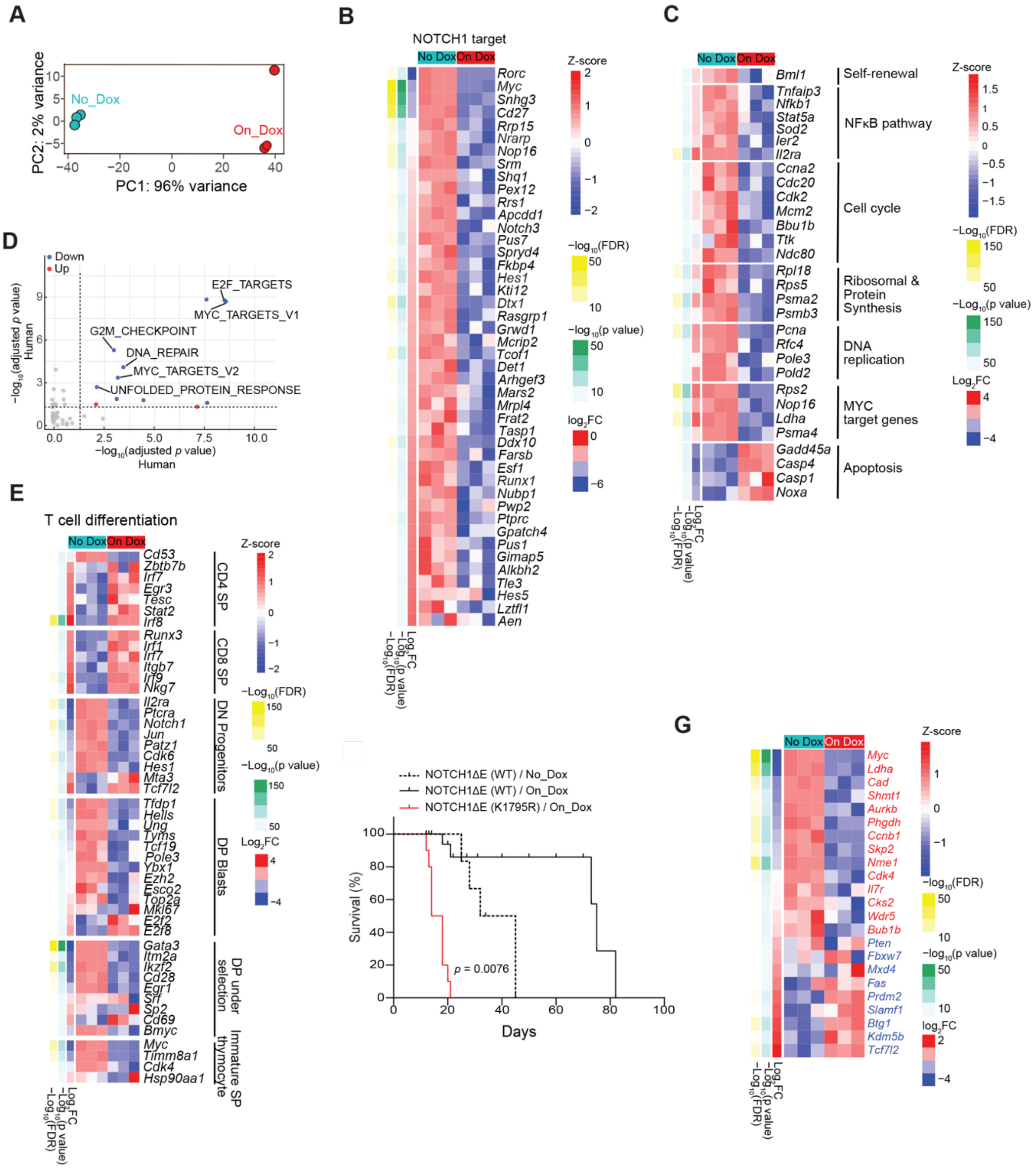


**Figure S9**

**Figure S9. UVRAG expression suppresses NOTCH1-driven transcriptomic programs and leukemia-initiating cell function *in vivo*, related to Figure 5.**

(A) Principal component analysis (PCA) of RNA-seq profiles from *GFP*$^{+}$;*iUvrag* T-ALL cells isolated from spleens of mice treated with Dox (On_Dox) or untreated (No_Dox). n = 3. Fisher's exact and hypergeometric tests with Benjamini-Hochberg correction is used.

(B and C) Heatmaps showing differential expression of canonical NOTCH1 target genes (B) and genes involved in indicated pathways (C) in On_Dox *vs.* No_Dox GFP$^{+}$; i*Uvrag* T-ALL cells. (n =

3). $Log_2$ fold change (FC) and $-Log_{10}$ false discovery rate (FDR) are color-coded.
(D) Enriched term clusters using differentially expressed genes (adjusted *p*-value < 0.05, $Log_2FC > 0.5$) in *UVRAG* high *vs.* *UVRAG* low (using the quantile as threshold) T-ALL subsets with high NOTCH1 activity signature (Palomero et al., 2006b) (using median as threshold) in humans and mice, using Enrichr and MSigdb Hallmark 2020 gene sets. Dashed line indicates adjusted $p = 0.05$.
(E) Heatmaps showing transcriptomic changes in gene sets associated with thymocyte development and T cell differentiation(Li et al., 2021) in On_Dox *vs.* No_Dox cells in (B).
(F) Kaplan-Meier survival curve of mice transplanted with *iUvrag* BM cells transduced with retroviral NOTCH1ΔE WT (black and dotted line) or K1795R mutant plasmid (red line) and treated with or without Dox. n = 5 mice per group. *p* value was calculated using the log-rank test for trend.
(G) Heatmap of LIC-associated gene expression in $GFP^+$; *iUvrag* T-ALL cells from Dox-treated *vs.* untreated mice (n = 3).

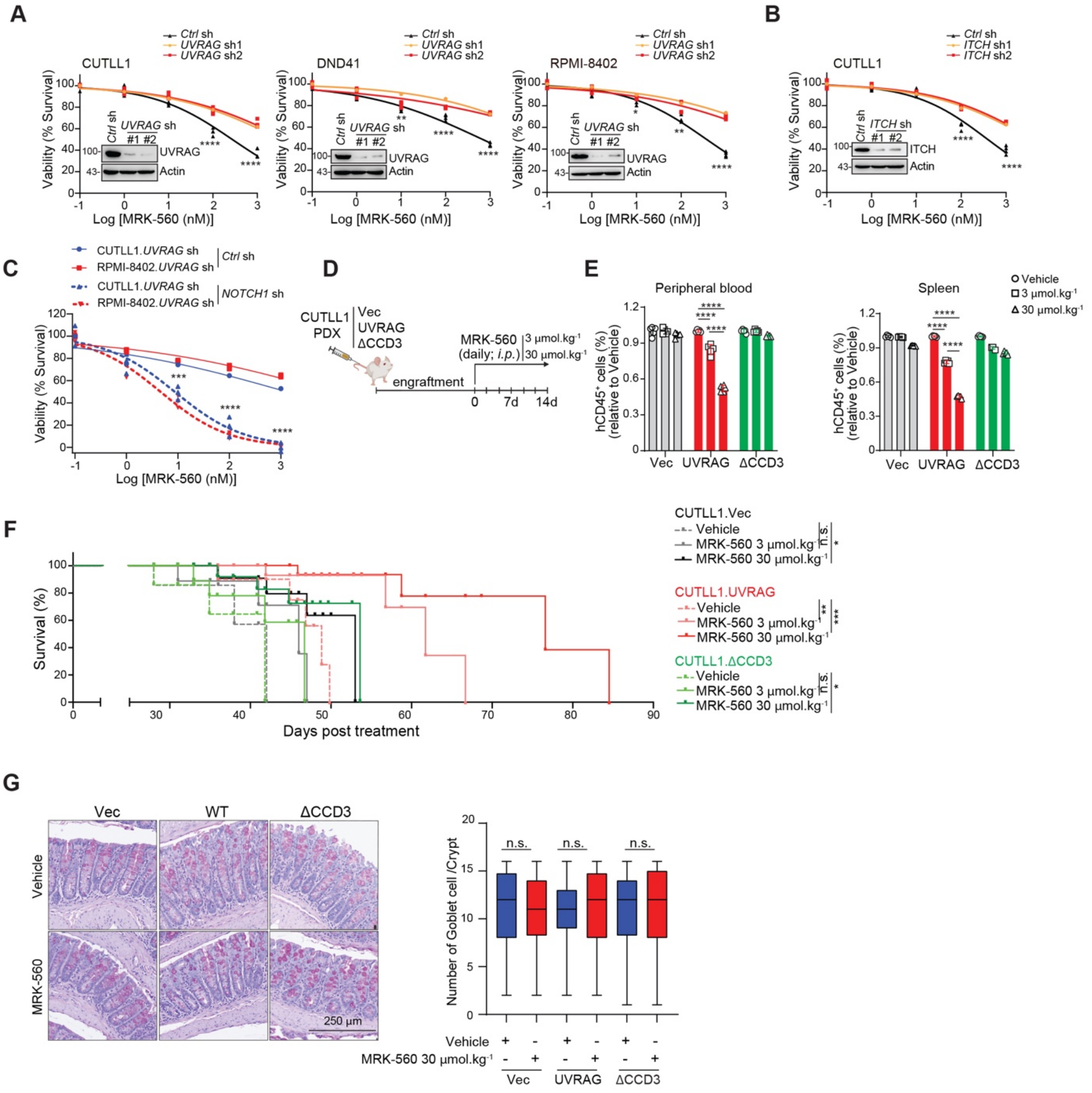


**Figure S10**

**Figure S10. UVRAG enhances T-ALL sensitivity to γ-secretase inhibition by depleting NOTCH1ΔE, related to Figure 6.**

(A and B) Cell viability assays of CUTLL1, DND41, and RPMI-8402 T-ALL cell lines stably expressing *Ctrl* or *UVRAG* shRNAs (A), or expressing *Ctrl*, *ITCH* shRNAs (B), treated with MRK-560 at the indicated concentrations for 7 days. IB confirms protein knockdown.

(C) Depletion of *NOTCH1* restores MRK-560 in *UVRAG*-deficient CUTLL1 and RPMI-8402 cells.

(D) Experiment design for *in vivo* therapy: NSG mice transplanted with CUTLL1 cells stably expressing Vec, UVRAG WT or ΔCCD3 (*via* tail vein injection) were treated with Vehicle or MRK

560 (3 or 30 mmol $kg^{-1}$, *i.p.* daily for 14 days) after disease establishment (human $CD45^+$ cells reached ~10% in peripheral blood lymphocytes).
(E) Quantification of human $CD45^+$ ($hCD45^+$) cells in peripheral blood (left) and spleens (right) of treated mice (n = 5 per group). Fold changes reflect MRK-560 vs. Vehicle within each genotype.
(F) Kaplan-Meier survival curve of mice from (D), with treatment stratified by genotype and MRK-560 dose. *p* values calculated by log-rank (Mantel-Cox) test.
(G) Representative images of PAS staining of intestinal sections from mice treated with vehicle or MRK-560. Goblet cell numbers per crypt were quantified across treatment groups (n = 5 per group). Scale bar, 250 mm.
Data in (A-C) and (E) represent mean ± s.d. from 3 biologically independent experiments, analyzed by two-tailed Student's *t* test or one-way ANOVA followed by Tukey's *post hoc* test. Boxplot data in G: the center line indicates median, boxes indicate interquartile range (IQR, $25^{th}$-$75^{th}$ percentiles), and whiskers extend to minimum and maximum values; analyzed using Kruskal-Wallis with *post hoc* Dunn's test. *, $p < 0.05$; **, $p < 0.01$; ***, $p < 0.001$; ****, $p < 0.0001$; n.s., not significant.

**Table S1: T-ALL patient sample information**

| Patient | Age (Y) | Gender | Sample | Blast (%) | Cytogenetic |
|---|---|---|---|---|---|
| P1 | 5 | M | Peripheral blood | 90 | *FBXW7* mutation, *PHF6* mutation, *CDK2NA* deletion |
| P2 | 8 | M | Bone marrow | 88 | *CDK2NA* and *PTEN* mutation |
| P3 | 11 | M | Bone marrow | 95 | *NOTCH1* (HD domain) and *PTEN* mutation, *CDK2NA* deletion |
| P4 | 9 | M | Bone marrow | 81 | *FBXW7* mutation, *TP53* mutation, *CDK2NA* mutation, *USP7* mutation |
| P5 | 9 | M | Peripheral blood | 94 | *STIL-TAL1* fusion, *CDK2NA* and *CDK2NB* deletion |
| P6 | 15 | M | Peripheral blood | 82 | *PTEN* deletion (typically associated with lack of NOTCH1 mutations,), MYB duplication |
| P7 | 13 | M | Peripheral blood | 96 | *STIL-TAL1* fusion, *CDK2NA* and *CDK2NB* deletion |
| P8 | 4 | M | Bone marrow | 90 | *STIL-TAL1* fusion, *NOTCH1* mutation, *PHF6* mutation, *STAT5B* mutation |

Y, year; M, male; %, percentage

**Table S2: Primers for genotyping and RT-qPCR**

| REAGENT or RESOURCE | SOURCE | IDENTIFIER |
|---|---|---|
| Oligonucleotides for mouse strain genotyping | | |
| *R26-rtTA*M2*<br>Forward: AAAGTCGCTCTGAGTTGTTAT (common) | Integrated DNA Technologies | N/A |
| *R26-rtTA*M2*<br>Reverse: GGAGCGGGAGAAATGGATATG (wt)<br>GCGAAGAGTTTGTCCTCAACC (mut) | Integrated DNA Technologies | N/A |
| *Uvrag*<br>Forward: AAGCAGAGGAAATCATCG | Integrated DNA Technologies | N/A |
| *Uvrag*<br>Reverse: TATGTTTCAGGTTCAGGG | Integrated DNA Technologies | N/A |
| *TRE*<br>Forward: AGATCGCCTGGAGAATTCG | Integrated DNA Technologies | N/A |
| *TRE*<br>Reverse: AGGTAGACTTTCCACTCT | Integrated DNA Technologies | N/A |
| Oligonucleotides for RT-qPCR | | |
| *Human UVRAG*<br>Forward: AGGATTACTTTGTATGCGGTGTC | Integrated DNA Technologies | N/A |
| *Human UVRAG*<br>Reverse: CAGGTTGGGAAGGGTTTGC | Integrated DNA Technologies | N/A |
| *Human MYC*<br>Forward: CTGAGGAGGAACAAGAAGATGAG | Integrated DNA Technologies | N/A |
| *Human MYC*<br>Reverse: TGTGAGGAGGTTTGCTGTG | Integrated DNA Technologies | N/A |
| *Human IL7R*<br>Forward: TCGCTCTGTTGGTCATCTTG | Integrated DNA Technologies | N/A |
| *Human IL7R*<br>Reverse: GGAGACTGGGCCATACGATA | Integrated DNA Technologies | N/A |
| *Human NRARP*<br>Forward: ACCAACTGCGAGTTCAAC | Integrated DNA Technologies | N/A |
| *Human NRARP*<br>Reverse: ATGAGATAGAGCACGATGTC | Integrated DNA Technologies | N/A |
| *Human HES1*<br>Forward: CGTTCGCTTGGTTAGCAGTG | Integrated DNA Technologies | N/A |
| *Human HES1*<br>Reverse: GCAGATAATGGATTCCCTCAAA | Integrated DNA Technologies | N/A |
| *Human HEY1*<br>Forward: GCGCACGCCCTTGCT | Integrated DNA Technologies | N/A |
| *Human HEY1*<br>Reverse: GCCAGGCATTCCCGAAAT | Integrated DNA Technologies | N/A |
| *Human GAPDH*<br>Forward: CCCATCACCATCTTCCAGG | Integrated DNA Technologies | N/A |
| *Human GAPDH*<br>Reverse: CCATCACGCCACAGTTTCC | Integrated DNA Technologies | N/A |
